# Why do hippocampal mossy cells matter? It depends on the frequency and context

**DOI:** 10.1101/2021.12.04.467951

**Authors:** Katarina Kolaric, Christina Strauch, Yingxin Li, Sasha Woods, Marinho A. Lopes, Natasha Sivarajah, Catia M. Teixeira, Jason Lerch, Paul W. Frankland, Mark Henkelman, Zuner A Bortolotto, E. Clea Warburton, Zafar I Bashir, Denise Manahan-Vaughan, Denize Atan

## Abstract

The discrimination of similar episodes and places, and their representation as distinct memories, depend on a process called pattern separation that relies on the circuitry of the hippocampal dentate gyrus (DG). Mossy cells (MCs) are key neurons in the circuitry, but how they influence DG network dynamics, function, and seizure risk has not been fully elucidated. We found the net impact of MCs was inhibitory at physiological frequencies connected with learning and behaviour, and their absence associated with deficits in pattern separation and spatial memory; at higher frequencies, their net impact was excitatory, and their absence protected against seizures. Thus, MCs influence DG outputs in a highly dynamic manner that varies with frequency and context.

**One-Sentence Summary:** Hippocampal mossy cells are required for learning and memory; but their absence protects against seizures.

## Main Text

The discrimination of similar episodes and places, and their representation as distinct memories, depends on a process called pattern separation. Several human, animal, and computational studies have implicated the hippocampal dentate gyrus (DG) as a pattern separator (*1–4*); and pathologies involving the DG, like temporal lobe epilepsy (TLE) and mild cognitive impairment, are associated with pattern separation deficits (*5, 6*). To function as a pattern separator, DG circuitry must rapidly transform neural inputs with overlapping spatiotemporal firing patterns into distinct outputs (*3*); but despite receiving thousands of excitatory perforant path (PP) inputs from the entorhinal cortex (EC), only ∼5% of the major DG output neurons, the granule cells (GCs), are active at any time (*7*). This sparse GC firing pattern reduces the probability of collisions between similar DG inputs, an advantage for pattern separation, yet how this is achieved by DG circuitry is not clear. In this regard, glutamatergic mossy cells (MCs) found in the DG hilus are ideally connected for a role in pattern separation; firstly, MCs drive network excitation through direct synapses with GCs along the septotemporal axes of the ipsilateral and contralateral hippocampus (*8*), thereby providing a system to orthogonalize information to distant GCs; secondly, MCs contribute to the strong lateral inhibition and winner-takes-all competition between GCs through their connections to the dense intrinsic network of inhibitory interneurons (INT) within the DG (*9, 10*).

However, a distinct disadvantage of the intrinsic properties and connectivity of MCs is their vulnerability to excitotoxic cell death. Firstly, MCs have characteristically high rates of spontaneous excitatory postsynaptic potentials that are potentiated by short periods of depolarisation (*11*). Secondly, MCs both receive and send glutamatergic axons directly to and from GCs, which can potentially create an escalating feedback loop of excitation. Yet how MC death influences network dynamics and susceptibility to uncontrolled electrical activity (seizures) is still debated. Some studies have indicated the net impact of MCs on GCs is inhibitory via their indirect MC→INT→GC connections, so seizure susceptibility increases after their death (*12, 13*); conversely, seizure risk could be reduced by the loss of their direct MC→GC connections (Figure 1A)(*14*). One possible explanation is that the balance between GC excitation and inhibition by MCs varies depending on the stimulation frequency and context. We therefore used magnetic resonance imaging (MRI), immunofluorescent microscopy, *in vitro* and *in vivo* electrophysiology, behavioural tasks, and computational modelling to investigate how MCs influence the neural dynamics of DG circuity in mice at different frequencies and their impact on pattern separation, learning, memory and seizures.

**Figure 1.**
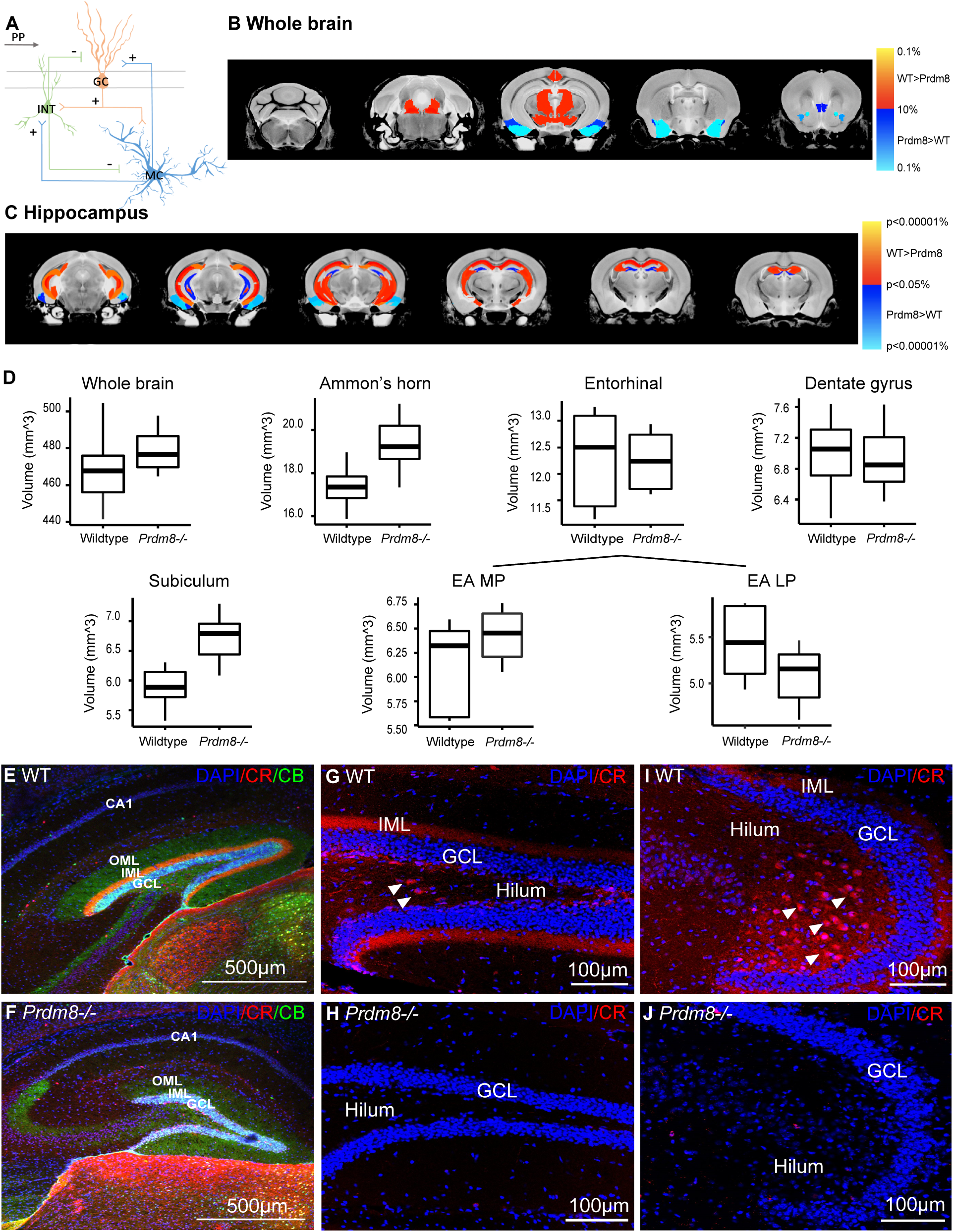
The *Prdm8^-/-^* dentate gyrus (DG) is smaller than wild type (WT) on MRI scans because mossy cells (MC) are absent. (A) Schematic of direct MC→granule cell (GC) connections and indirect MC→interneuron (INT)→GC connections. (B-D) Differences in volume between *Prdm8^-/-^* and WT mice (n=6|7) of (B) whole brain at a false discovery rate (FDR) threshold of 10%, and (C) hippocampus at an unthresholded p<0.05 and superimposed on MRI brain images. (D) Overall, *Prdm8^-/-^* whole brain and hippocampus was larger than WT (FDR<10%). Hippocampal subfield analysis showed the DG was disproportionately smaller (p<0.05) in *Prdm8^-/-^* mice compared with WT. (E-J) Representative confocal micrographs of sagittal hippocampal sections from adult *Prdm8^-/-^* and WT mice, counterstained with DAPI (4’-6-diamidino-2-dihydrochloride). (E, F) Calbindin^+^ (CB) GC bodies, dendrites and projections were intact in *Prdm8^-/-^* hippocampus but calretinin^+^ (CR) MC projections were absent from the inner molecular layer (IML). Note: CR labels some INT and immature GCs. (G-J) CR^+^ MC bodies (white arrow heads) and axons in the IML were absent from dorsal (G, H) and ventral (I, J) *Prdm8^-/-^* DG. OML=outer molecular layer; ML=molecular layer; EA=entorhinal; MP=medial perforant path.

### MCs are selectively absent from the DG of *Prdm8^-/-^* mice

We have previously reported *Prdm8^eGFP/eGFP^* mice (hereafter called *Prdm8^-/-^* mice) have a night blindness phenotype and their hippocampal commissure is almost completely absent (*15, 16*). As the hippocampal commissure mainly comprises axonal projections from MCs and DG interneurons to the contralateral hippocampus, we investigated *Prdm8^-/-^* mice for hippocampal abnormalities on MRI scans. Overall, *Prdm8^-/-^* brains were larger than wild type (WT); and hippocampal subfield analysis showed the CA1, CA2 and CA3 regions were also larger, while the DG was disproportionately smaller (Figure 1B-D). Because the DG comprises GCs, MCs and several interneuron populations, identifiable from their morphology and protein expression (*17*), we next used immunofluorescence microscopy to look for changes in these cell populations in the *Prdm8^-/-^* DG. We used antibodies to calbindin (CB) to label mature GC bodies, dendrites and mossy fibres traversing the hilar region to CA3, whereas antibodies to AMPA receptor subunits GluR2/3 label MC bodies and calretinin (CR) antibodies label MC bodies and projections to the inner molecular layer (IML) of the ipsilateral and contralateral hippocampus. Most DG interneurons are parvalbumin^+^ (PV) basket and axo-axonic cells or somatostatin^+^ (SOM) hilar perforant pathway (HIPP) cells, which form recurrent inhibitory connections between MCs and GCs; because of their connectivity, PV^+^ and SOM^+^ interneurons are the most likely candidates for mediating the indirect di-synaptic inhibition of GCs by MCs (Figure 1A)(*3*). Although PRDM8 was expressed in GCs and MCs but not PV^+^ or SOM^+^ interneurons or astrocytes of WT DG (Figure S1), we found GluR2/3 and CR immunoreactivity was almost completely absent from the *Prdm8^-/-^* DG while the morphology and organisation of GCs, PV^+^ and SOM^+^ interneurons were intact (Figures 1E-J, S2,3, Table S1). Overall, these results showed that the *Prdm8^-/-^* DG is smaller than WT DG due to the selective absence of MCs.

### MCs regulate the gamma frequency-dependent inhibition of GCs resulting from PP stimulation

GCs receive inputs from the EC via the medial (MPP) and lateral (LPP) perforant pathways: the MPP conveys spatial information like object location, whereas the LPP carries non-spatial information like olfactory and visual cues (*18*). Low frequency theta oscillations (5-10Hz) in the hippocampus are modulated by locomotor speed, whereas slow gamma frequency oscillations (20-50Hz) correlate with object exploration and associative memory for object encounters (*19*). Knowing that MCs were absent from the *Prdm8^-/-^* DG but the MPP was intact (Figures 1B-D, S2), we examined the effect of MPP stimulation at physiological frequencies (5, 20, 50Hz) on GC activity *in vitro* recorded in acute hippocampal slices. We first confirmed that basal GC responses to MPP stimulation were normal in the *Prdm8^-/-^* DG and found short-term plasticity was intact (Figure S4, Tables S2-5). We next investigated the short-term dynamics of GC responses to MPP stimulation in the *Prdm8^-/-^* DG to determine whether GC activity was reduced (because direct excitatory MC→GC connections were absent) or increased (because indirect inhibitory MC→INT→GC connections were no longer driven by MCs) (Figure 1A).

Our results showed the peak amplitudes of field excitatory post-synaptic potentials (fEPSPs) recorded in the GC population of the *Prdm8^-/-^* DG were less depressed compared with WT during 20Hz and 50Hz stimulation, but not during 5Hz stimulation (Figure 2A-F, Tables S6&7, S14-16). To confirm the net impact of MCs on GC activity was inhibitory via the di-synaptic MC→INT→GC pathway, we recorded GC responses to MPP stimulation in the presence of GABAA receptor antagonist picrotoxin (PTX) to block synaptic transmission via inhibitory GABAergic interneurons. Like GC responses recorded from *Prdm8^-/-^* hippocampal slices, we found GC responses in PTX-treated WT slices were disinhibited during 20Hz and 50Hz stimulations, but not during 5Hz stimulations (Figure 2G-L, Tables S8&9, S14-16). Furthermore, GC responses recorded from PTX-treated *Prdm8^-/-^* and WT slices were similar (Figure 2M-R, Tables S10-16). These results suggested the absence of MCs from DG circuitry reduces the inhibitory drive of the MC→INT→GC pathway at low gamma frequencies usually associated with spatial memory processing.

**Figure 2.**
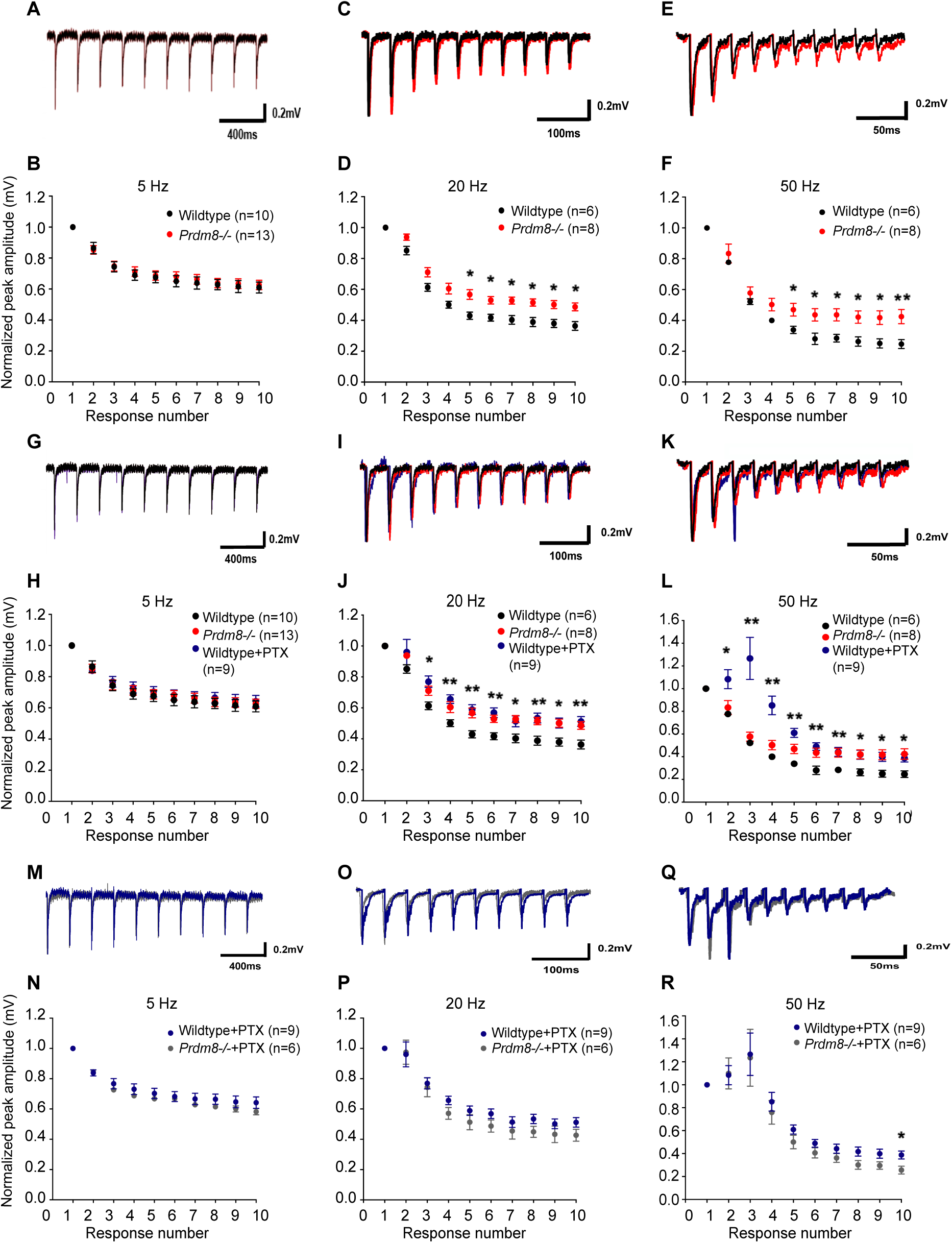
Granule cell (GC) responses are less depressed to 20Hz and 50Hz medial perforant path (MPP) stimulation in the *Prdm8^-/-^* dentate gyrus (DG) compared with wild type (WT). (A-R) Example waveforms and normalized peak field excitatory post-synaptic potential (fEPSPs) amplitudes recorded in GCs during 5Hz, 20Hz and 50Hz stimulations of the MPP in *Prdm8^-/-^* and WT hippocampal slices. (A, B) There were no differences in peak fEPSP amplitudes between genotypes (n=13|10) during 5Hz stimulations (p=0.760), but significant differences were observed between *Prdm8^-/-^* and WT GC responses (n=8|6) (C, D) during 20Hz stimulations (p=0.006) and (E, F) 50Hz stimulations (p=0.025). (G-L) Blocking GABAergic inhibition in WT hippocampal slices with picrotoxin (PTX) recapitulated the disinhibition of GC responses recorded in *Prdm8^-/-^* slices (n=9|8) (I, J) during 20Hz (p=0.507) and (K, L) 50Hz stimulations (p=0.008). (M-R) GC responses to MPP stimulation in *Prdm8^-/-^* and WT hippocampal slices (n=6|9) in the presence of PTX were similar (M, N) at 5Hz (p=0.339), (O, P) 20Hz (p=0.536) and (Q, R) 50Hz (p=0.770). Statistical comparisons are summarized in Tables S6-16.

### MCs are required for long-term potentiation (LTP) of synaptic transmission at MPP-GC synapses after high frequency stimulation (HFS)

LTP of synaptic transmission in the hippocampus is generally accepted to be the electrophysiological correlate of hippocampal-dependent learning; LTP persists over several days to weeks and enables encoding of long-term spatial representations in the hippocampus (*20*). Having investigated the short-term dynamics of GC responses to MPP stimulation *in vitro*, we next tested the long-term dynamics of GC responses to HFS of the MPP *in vivo* that normally induces LTP of synaptic transmission at MPP-GC synapses. Baseline input-output population spike (PS) amplitudes and fEPSP slope of GC responses were larger in recordings from *Prdm8^-/-^* mice than WT mice (Figure 3A-E, Table S17&18), although there was little difference in fEPSP slope between genotypes *in vitro* (Figure S4A,B, Table S2). Additionally, there were no differences between genotypes in their GC responses (PS amplitude and fEPSP slope) to low frequency test pulses (0.025Hz) after normalization to baseline (Figure 3F-H, Tables S19&20). Yet HFS of the MPP had the reverse outcome: the PS amplitude of GC responses were larger in WT mice than *Prdm8^-/-^* mice. This effect was first evident immediately after HFS and lasted about 60 minutes after 1 train of HFS (Figure 3I-K, Table S21) and persisted for at least 24 hours after 3 trains of HFS (Figure 3L-N, Tables 23) while differences between genotypes in the fEPSP slopes of their GC responses were not statistically significant (Figure 3I-N, Table S22&24). Taken together, these results suggest that under baseline conditions *in vivo*, proportionally more GCs are active in *Prdm8^-/-^* mice than WT; in other words, the net effect of MCs on GCs is inhibitory under conditions when GC activity is influenced by physiological EC inputs and DG network dynamics. In contrast, the immediate and long-term potentiation of GC responses to HFS are impaired in the *Prdm8^-/-^* DG, suggesting the net influence of MCs on GC activity is excitatory at high frequencies.

**Figure 3.**
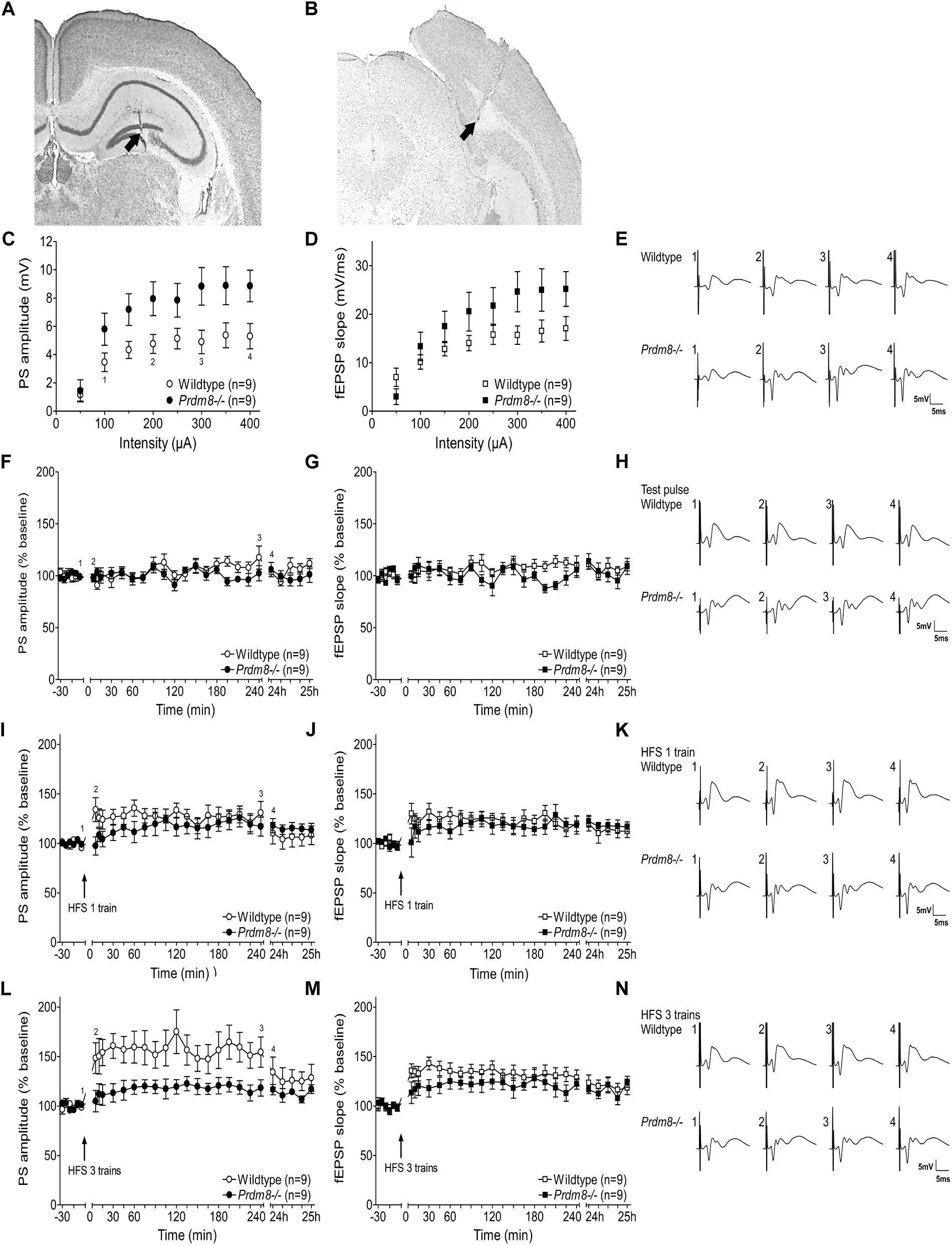
Long-term potentiation of granule cell (GC) responses to high frequency stimulation (HFS) is impaired in *Prdm8^-/-^* dentate gyrus (DG) *in vivo* but more GCs are active under baseline conditions. (A, B) Nissl staining showing the recording electrode in the GC layer of the DG (A, black arrow) and stimulating electrode in perforant path (B, black arrow). (C) The population spike amplitudes (PS)(p=0.025) and (D) slopes of field excitatory post-synaptic potential (fEPSPs)(mV/ms)(p<0.001) were larger in the stimulus-responses (input-outputs) of GCs in *Prdm8^-/-^* animals than WT (n=9|9); (E) example waveforms. (F-H) There were no baseline differences between the normalized GC responses of *Prdm8^-/-^* and WT animals (n=9|9) in (F) PS amplitude (p=0.227) or (G) fEPSP slope (p=0.242); (H) example waveforms. (I) One train of HFS at 400Hz potentiated the PS amplitudes (p<0.001) of GC responses in WT, not *Prdm8^-/-^* animals (n=9|9), but (J) fEPSP slopes (p=0.166) and (K) waveforms were no different. (L) Long-term potentiation of PS amplitudes (p=0.003) in GC responses was observed in WT, not *Prdm8^-/-^* animals (n=9|9), for up to 24 hours; (M) no short- or long-term differences in fEPSP slope (p=0.114) or (N) waveform were observed between genotypes. Statistical comparisons are summarized in Tables S17-24.

### MCs influence hippocampal-dependent spatial learning, memory and behavioural pattern separation

Given the putative role of the DG in spatial and contextual coding and pattern separation (*7, 21*), and the impaired LTP of MPP-GC responses we observed in *Prdm8^-/-^* mice, we next assessed the role of MCs in (i) hippocampal-dependent spatial learning and memory using the Morris Water Maze (MWM)(*22*); and (ii) pattern separation using the behavioural object pattern separation (OPS) task (*23*). The OPS task is a modification of the object location task in which "pattern separation load” can be varied by adjusting the distance of one of two familiar objects during the test phase, enabling the quantification of differences in behavioural pattern separation between groups (the load is increased by moving the object a shorter distance because the similarity between training and test conditions is greater). Unlike the MWM, there is no reward or stress involved in the OPS task to confound results, and it is very similar to tasks that test behavioural pattern separation in humans (*4*). Additionally, we performed several control behavioural tasks. Firstly, we used the contextual fear conditioning task, another associative learning task that is more dependent on amygdala than hippocampus (*24*), to assess baseline anxiety-related behaviour, since mice dislike enforced swimming and the results of the MWM may be confounded by baseline differences between animals (*25*). Secondly, we used the novel object recognition (NOR) task since it relies solely on extrahippocampal perirhinal cortex, and does not incur stress or involve reward (*26*). Finally, *Bhlhb4^-/-^* mice were used as controls in experiments where visual cues may have been relevant, since *Bhlhb4^-/-^* mice do not lack MCs (Figure S5) but share the same night blindness phenotype as *Prdm8^-/-^* mice (*27*). Additionally, all mice were screened using the SHIRPA protocol of behavioural measures.

The most striking finding was that *Prdm8^-/-^* mice were impaired in pattern separation but their behaviour on non-hippocampus-dependent associative learning tasks was preserved (Figure 4A,B). Compared with control mice, *Prdm8^-/-^* mice were less able to discriminate changes in object position with the highest pattern separation loads on the OPS task. These behavioural differences did not occur due to any positional bias in exploration between groups or visual impairment, since *Bhlhb4^-/-^* mice performed as well as WT controls (Tables S25-27). Additionally, *Prdm8^-/-^* mice took longer to learn the location of a hidden platform on the MWM compared with control mice, and they were unable to recollect the location of the target platform when it was removed. Again, these deficits were not secondary to visual impairment; nor were they due to any impairment in swimming ability (Figure 4C-F, Tables S28-30). *Prdm8^-/-^* mice could successfully discriminate novel from familiar objects in the NOR task, they were no more fearful than WT mice and could learn to associate the contextual fear conditioning chamber with foot shock, indicating they were not globally impaired in their learning and memory; nor did they have any major deficits identified by SHIRPA screening (Figure 4G-J, Tables S31-36). Together, these results showed that *Prdm8^-/-^* mice are selectively impaired in behavioural pattern separation and hippocampal-dependent spatial learning and memory, and provided strong behavioural evidence to support computational models of the DG that have implicated MCs in pattern separation (*21*).

**Figure 4.**
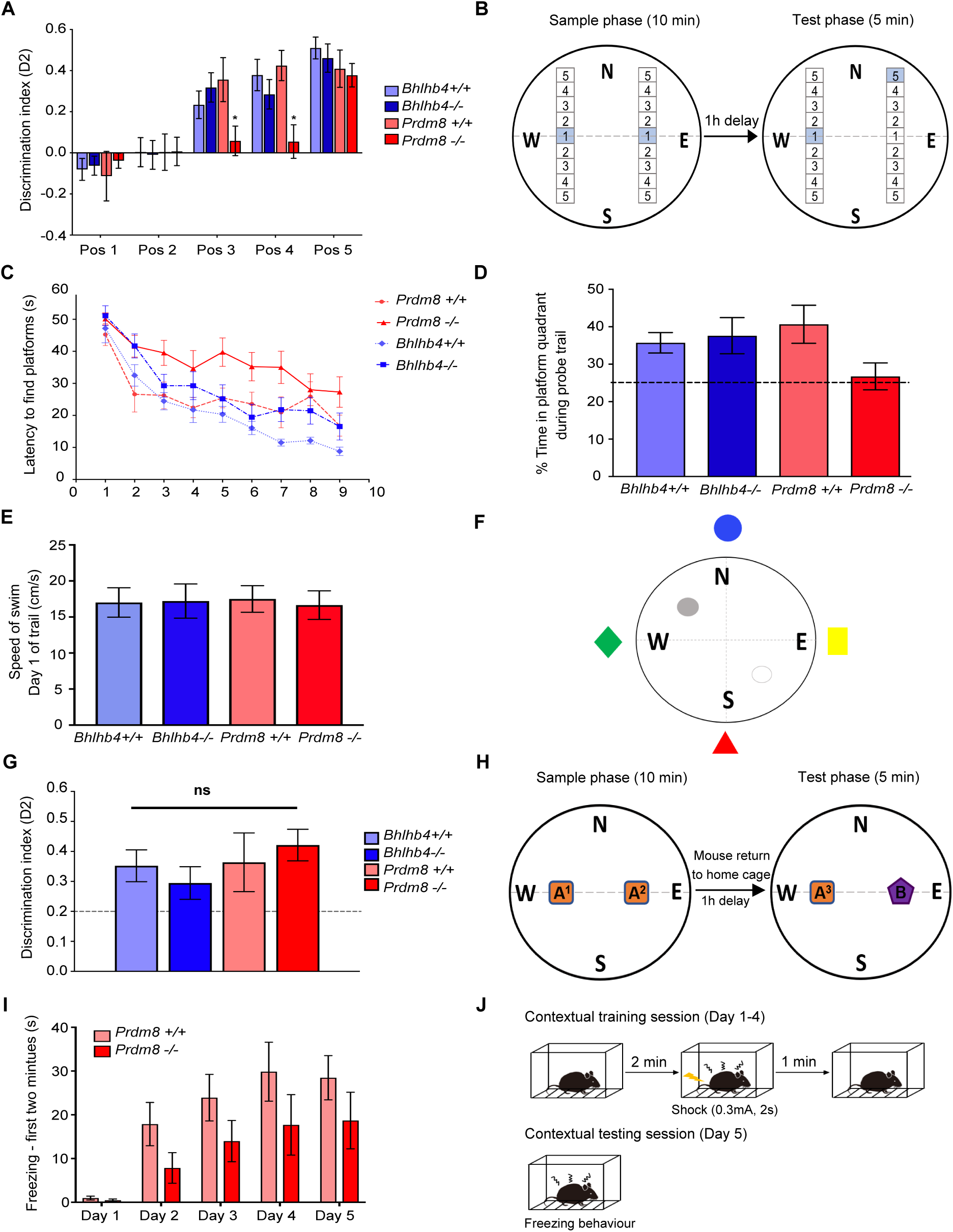
*Prdm8^-/-^* mice are significantly impaired in object pattern separation (OPS). (A) *Prdm8^-/-^* mice (n=11) were impaired in their discrimination of an object moved to positions 3 (p=0.049) and 4 (p=0.027), not position 5 (p=0.991), of the OPS task compared with controls (*Prdm8^+/+^*=8; *Bhlhb4^-/-^=*15; *Bhlhb4^+/+^*=10); mean discrimination index (D2) ±standard error of the mean (SEM) is plotted per genotype and position. (B) Schematic of OPS task. (C) *Prdm8^-/-^* mice (n=11) were slower to learn the Morris water maze (MWM) over 9 training days compared with (*Prdm8^+/+^*) controls (n=10)(p=0.330); control *Bhlhb4^-/-^* mice (with night blindness) behaved like *Bhlhb4^+/+^* controls (n=12|11)(p=0.933). (D) *Prdm8^-/-^* mice were impaired in their recall of the hidden platform (day 10 probe trial) compared with controls (n=11|10)(p=0.038). (E) Swimming speed was no different between genotypes. (F) Schematic of MWM. (G) *Prdm8^-/-^* mice (n=11) were not impaired in novel object recognition (NOR) compared to controls (*Prdm8^+/+^*=8; *Bhlhb4^-/-^*=15; *Bhlhb4^+/+^*=10)(p=0.127); D2 ±SEM is plotted per genotype. (H) Schematic of NOR task. (I) *Prdm8^-/-^* mice were slower to learn the contextual fear conditioning task (CF) and froze for less time than (*Prdm8^+/+^*) controls (n=9|12) on each training day (p=0.657). (J) Schematic of CF task. Statistical comparisons are summarized in Tables S25-S36.

### The absence of MCs from the DG protects against seizures

Existing computational models of the DG that have implicated MCs in pattern separation are based on the intrinsic electrophysiological properties and connectivity of the DG’s cell populations at a microscopic level (*21*). In contrast, neural mass models are used to simulate macroscopic spatiotemporal network dynamics and the transition from interictal to ictal activity in epilepsy (*28*). To understand how MCs might influence this transition, we developed a neural mass model following the implementation by López-Cuevas *et al* (*28*) and based on pre-existing biophysical models of the DG (Figure 5A, Table S37)(*21*). In the model, HIPP cells receive PP inputs and provide feed-forward inhibition to GCs and BCs provide GCs with recurrent feedback inhibition (*21*). We further considered GC→HIPP connections (absent in (*21*)) since HIPP interneurons are now known to be targeted to a greater extent by GCs than PP axons (*29*). When GCs were activated by sufficiently strong stimuli, the neural mass model generated 3-4Hz sharp-wave oscillations like the spike-wave discharges observed in mouse models of epilepsy (*30*). Seizure-like activity was no longer produced when MCs were removed from the model, even when the stimulus intensity was above the threshold that elicited seizure-like activity in the intact circuitry (Figure 5B-D). Furthermore, seizure-like activity was prevented by removing MC→GC connections but not MC→BC connections, suggesting the MC→GC excitation loop was required for seizure generation in the DG (Figures 5E-H).

**Figure 5.**
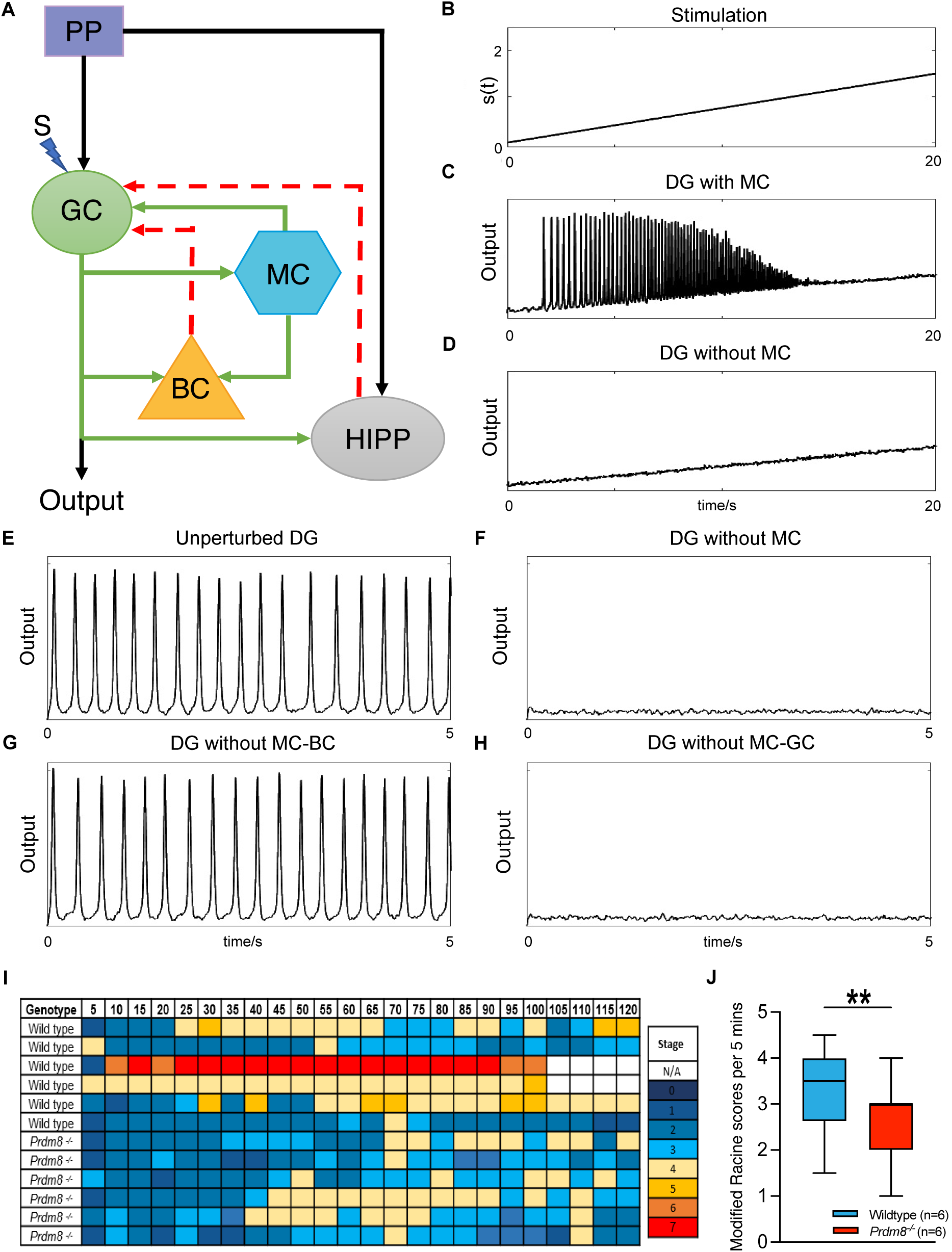
The absence of mossy cells (MC) from dentate gyrus (DG) circuitry protects against seizures. (A) Schematic of neural mass model. The perforant path (PP) excites granule cells (GC) and hilar perforant path-associated (HIPP) cells. GC excite MC, basket cells (BC), and HIPP cells. MC excite GC and BC. Both BC and HIPP cells inhibit GC. S represents an external stimulus delivered to GC. (B-D) In the model, increasing stimulation strength induced seizure-like activity when MC were present, but not they were when absent. (B) When stimulated, outputs from the DG model resembled seizure activity. (E, F) Removal of MC from the model prevented the emergence of seizure-like activity. (G) Removal of MC→BC connections did not prevent seizure-like activity. (H) Removal of MC→GC connections prevented seizure-like activity. (H) Modified Racine scores for seizure severity were recorded every 5 minutes for two hours following intraperitoneal injections of kainic acid in *Prdm8^-/-^* and wild type (WT) mice (n=6|6). Two WT mice were terminated after 100 minutes in accordance with endpoints in the Home Office project license. (J) Median scores were higher at each 5-minute interval in WT mice compared with *Prdm8^-/-^* mice (p<0.007). Supporting data is available in Tables S37-40.

Although no spontaneous behavioural or electrographic seizures were observed during *in vivo* experiments, we next investigated whether *Prdm8^-/-^* mice behaved differently to WT in the 2 hours following the induction of seizures by intraperitoneal injections of kainic acid. We found that significantly more WT mice reached the highest stages of the modified Racine scale (corresponding with more severe generalised seizures) than *Prdm8^-/-^* mice; whereas the duration and onset of seizures were no different between genotypes (Figure 5I,J, Tables S38-40). Together with the findings of the neural mass model, these data suggest that MCs influence both seizure generation and generalisation. Since MCs are vulnerable to excitotoxicity, we conclude that MC death is caused by seizures and not vice versa; indeed, MC death is protective against seizures. The consequence is that MC death disrupts the physiological balance between excitation and inhibition in DG circuitry, resulting in impaired pattern separation, spatial learning and memory.

## Discussion

The sparse firing of GCs is essential to memory encoding and pattern separation in the DG (*7*). This is largely determined by the intrinsic properties of GCs and the dense inhibitory feedforward and feedback microcircuits within DG circuitry (*10, 31, 32*). Interneurons of the DG shape the input-output relationship of GCs, and they exert dynamic levels of spatiotemporal control over GC activity by providing feedforward inhibition at the start of a spike train and feedback inhibition of later spikes (*31, 33*). Our results suggest that MCs activate interneuron microcircuits which mediate the feedback inhibition of GCs, particularly at gamma frequencies linked to associative memory encoding (*19*). GC synapses with MCs and interneurons demonstrate strong frequency-dependent facilitation that increases at higher stimulation frequencies in the slow gamma range (50>20>10>1Hz)(*32*). Additionally, GCs exhibit temporal pattern separation that is more pronounced at gamma frequencies and depends on GABAA-mediated neurotransmission in the whole DG network (*34*). The frequency-dependency of pattern separation is driven by feedback inhibition, and not feedforward inhibition, in a biophysical computational model of the DG (*32*). Hence, we conclude that the absence of MCs from DG circuitry predominantly affects GC responses to slow gamma frequency stimulation, because MCs normally recruit inhibitory feedback microcircuits that are most active at these frequencies; when the MC-dependent feedback inhibition of GCs is removed, we observe behavioural deficits in pattern separation, spatial learning and memory.

In contrast, the auto-associative theory of spatial and episodic memory says that patterns of neural activity are stored in the hippocampus through strengthening interconnections between excitatory neurons (*35*). But, the drawback of such connections is the hippocampus becomes vulnerable to runaway excitation and seizures (*36*). High stimulation frequencies normally induce LTP of synaptic transmission at MPP-GC synapses, but we found LTP was impaired when MCs were absent. Our computational model and behavioral experiments further showed the absence of MCs was protective against seizures. We infer the net influence of MCs on GC responses is excitatory under these conditions. It has been hypothesized that MCs inhibit GCs close to their cell body via di-synaptic connections with local interneurons, but that they excite GCs distal to their soma via direct MC→GC projections along the septotemporal axes of ipsilateral and contralateral hippocampus (*17*). This anatomical hypothesis could explain why the absence of MCs is protective against seizure generalization in *Prdm8^-/-^* mice. Yet, we found GC responses were less inhibited in *Prdm8^-/-^* mice compared with WT mice under baseline conditions *in vivo* and to slow gamma stimulation *in vitro*; conversely after HFS, GC responses to MPP stimulation were potentiated in WT mice and not *Prdm8^-/-^* mice. Instead, our results suggest the balance between MC-mediated excitation and inhibition of GCs is frequency-dependent and not due to their local vs distal connectivity.

One important difference between *Prdm8^-/-^* mice and animal models of epilepsy is that MCs are selectively absent from the *Prdm8^-/-^* DG, whereas varying proportions of MCs and different interneuron populations are killed by seizures, resulting in wider disruption of DG circuitry in animal models of epilepsy (*37*). Indeed, some MCs may persist and become "irritable", which may be attributable to their high rate of spontaneous EPSPs and ability to self-potentiate, as well as aberrant excitatory inputs from mossy fibre sprouting (*11, 38*). Under these conditions, it is difficult to predict how experimental manipulations of DG circuitry will affect network dynamics and seizures. Hence, ’context’ is important. Here, we were able to investigate the individual effects of MCs on DG network dynamics by using the *Prdm8^-/-^* mouse model and our neural mass model, and conclude that seizures cause MC death and not vice versa.

In conclusion, our study shows that the contribution of MCs to information processing in the DG is not static. Rather, MCs regulate DG outputs in a highly dynamic manner, whereby context and afferent frequency determine whether their impact is inhibitory or excitatory. This contribution provides inherent support to key functions of the DG in processes such as pattern separation. On this basis, we predict that diseases such as TLE, which cause MC death (a cellular defect that can be detected macroscopically from hippocampal subfield analysis of MRI brain scans), will cause deficits in pattern separation, spatial learning and memory. Furthermore, deficits in pattern separation that occur with increasing age, mild cognitive impairment and dementia may well arise from pathologies which affect MCs and the inhibitory feedback microcircuits they regulate.

## Acknowledgments

**Funding:**

The Wellcome Trust SJ1197, WT084585] (KK, DA)

The Wellcome Trust 206401/Z/17/Z (ECW, ZIB)

The Wellcome Trust Institutional Strategic Support Fund (ISSF) to Cardiff University 204824/Z/16/Z (MAL)

National Eye Research Centre SJ1050, RJ5819, RJ6043 (DA, NS, SW)

China Scholarship Council and University of Bristol U135572-105 (YL)

Canadian Institutes of Health Research (CIHR) FDN143227 (CMT, PWF)

Ontario Brain Institute (OBI) to Province of Ontario Neurodevelopmental Disorders (POND) network (JL, MH)

European Research Council (ERC) 341089 (ZAB)

German Research Foundation SFB 874/B1 (project No.:122679504), SFB 1280/A04 (project No.: 316803389) (CS, DMV)

**Author contributions:**

Conceptualization: DA

Methodology: MAL, JL, PWF, MH, ZAB, ECW, ZAB, ZIB, DMV, DA

Investigation: KK, CS, YL, SW, MAL, NS, CMT, JL, ZAB, DA

Visualization: KK, CS, YL, SW, MAL, NS, CMT, JL, DA

Funding acquisition: KK, YL, SW, PWF, MH, ZAB, ECW, ZIB, DMV, DA

Project administration: DA

Supervision: PWF, MH, ZIB, DMV, DA

Writing – original draft: KK, CS, MAL, DA

Writing – review & editing: KK, CS, YL, SW, MAL, CMT, JL, PWF, MH, ZAB, ECW, ZIB, DMV, DA

**Competing interests:**

DA is on the clinical advisory board of Siloton Ltd.

**Data and materials availability:**

All data available in the main text or supplementary materials. Code for neural mass model is available on github: https://github.com/ml0pe5/DG_model.

## Supplementary Materials

Materials and Methods Fig S1-S6

Tables S1-S40

References 39-56

