## supplementary for "Why do hippocampal mossy cells matter? It depends on the frequency and context"

**This PDF file includes:**

Materials and Methods

Figs. S1 to S6

Tables S1 to S40

### Materials and Methods

#### *Animals*

Gene targeting and genotyping of the *Prdm8*$^{eGFP}$ (hereafter called *Prdm8*$^{-/-}$ mice) and *Bhlhb4* knock out mouse lines have been reported previously (*15, 39*). Both mouse lines were maintained on mixed C57BL/6J:129S genetic backgrounds and housed under a 12:12 hour light: dark cycle with food and water available ad libitum. Experiments were carried out in accordance with the UK Animals (Scientific Procedures) Act of 1986 and approved by the Animal Welfare and Ethical Review Body of the University of Bristol and Toronto Centre for Phenogenomics Animal Care Committee, except where otherwise stated.

#### *Magnetic resonance imaging and analysis*

Adult (8-12 weeks) *Prdm8*$^{-/-}$ mice (n=6) and sex- and litter-matched WT mice (n=7) were imaged by MRI using a 3D FSE T2 weighted sequence (*40*). Fixed brains were prepared for 7 Tesla MRI as previously described (*41*) and then brain volumes were segmented and compared using morphometric software analysis with a multi-atlas approach (*42*) combining mouse brain atlases from MICe (Mouse Imaging Centre, Toronto Centre for Phenogenomics) and the University of Queensland (*43-45*). First, an analysis of every brain structure in the atlas was compared against genotype with a false discovery rate of 10%, while covarying for overall brain volume. Secondly, an analysis of the hippocampus against genotype was performed using t-statistics at an uncorrected $p<0.05$ threshold. Differences in brain volumes between genotypes were plotted in graphical format and superimposed on brain images.

**Histology**

*Immunostaining*

Fixed brains were taken from at least 3 to 6 sex- and litter-matched pairs of adult *Prdm8*$^{-/-}$ and WT mice for immunolabelling experiments. Parasagittal 20µm cryosections were stained with the following antibodies: Calretinin (rabbit anti-CR, 1:1000), Calbindin (mouse anti-CB, 1:500), PRDM8 (rabbit or guinea pig anti-PRDM8, 1:1000), NeuN (rabbit anti-NeuN, 1:100), GFAP (rabbit anti-GFAP, 1:1000), Parvalbumin (guinea pig anti-PV, 1:700), Somatostatin (rat anti-SOM, 1:100), GluR2/3 (rabbit anti-GluR2/3 1:100). Sections were counterstained with 4',6-diamidino-2-phenylindole (DAPI)(Vector Laboratories Inc UK). All sections were imaged using a Leica Confocal microscope (Leica SP5) with a maximum magnification of x60.

*Cell counting*

For cell quantification, the mean number of cells expressing each marker was calculated from at least 3 consecutive parasagittal sections taken at 2.725mm, 2.35mm and 2.15mm lateral from Bregma to sample the dorsal and ventral DG throughout the septal to temporal pole of the hippocampus. All sections were z stacked at steps of 1.5µm with 20x magnification. The z stacks were collapsed to visualize cells that were counted using the Cell Counter plug-in for ImageJ software offline. An analysis of mean cell count against genotype for each marker was performed using the unpaired t-test and plotted with GraphPad Prism version 7 software.

*Nissl staining*

Coronal 20µm cryosections were taken from *Bhlhb4*$^{-/-}$ (n=4) and *Prdm8*$^{-/-}$ (n=4) fixed brain tissue as described above as well as from sex- and litter-matched WT mice. Sections were placed in 70%

ethanol for 5 minutes, briefly dipped in $H_2O$, then placed in cresyl violet acetate solution (Signa-Aldrich) for up to 1 hour. The sections were then briefly washed in 70% ethanol followed by 100% ethanol (twice) before leaving in Histo-Clear (National Diagnostics) for 15 minutes. Sections were then mounted with Histomount (National Diagnostics) and left overnight to dry before viewing under a light microscope.

***In vitro electrophysiology***

*Slice preparation*

Field recordings were performed at 30-32˚C on 400μm transverse slices perfused with artificial CSF (aCSF) containing: 124mM NaCl, 26mM $NaHCO_3$, 3mM KCl, 1.4mM $NaH_2PO_4$, 1mM $MgSO_4$, 10mM D-glucose and 2mM $CaCl_2$, equilibrated with 95% $O_2$/5% $CO_2$. Evoked field excitatory post-synaptic potentials (fEPSPs) were recorded using a pulled borosilicate glass microelectrode with resistance 2-5MΩ positioned near the proximal dendrites of GCs within the middle molecular layer (MML) of the dentate gyrus. To induce fEPSPs, stimulations were delivered via a tungsten bipolar stimulating electrode placed in the medial perforant pathway (MPP). All field data was acquired using an Axopatch 200B amplifier (Molecular Devices) connected to WinLTP acquisition software, filtered at 5kHz and digitized at a sampling rate of 20kHz (Digidata 1322A, molecular devices).

*Baseline recordings*

Paired pulse depression (paired pulse ratio<1) is characteristic of MPP-GC synapses. To measure the paired pulse ratio (PPR), paired pulses were delivered at inter-stimulus intervals of: 20ms, 50ms, 100ms, 200ms, 500ms and 1000ms. The rising slope (mV/ms) of fEPSPs following pulse 1 and pulse 2 were recorded offline and the average of four repetitions stimulated every 30 seconds were recorded for each interval. To calculate the PPR, the slope value of the second response was divided by the slope value of the first response.

Input-output responses were measured by initially stimulating at a 'minimal' intensity every 30 seconds (sufficient to produce an evident fEPSP from background noise) and were slowly increased from 0V in 5V increments until saturation of the response amplitudes and a maximal fEPSP was obtained. To analyze fEPSPs, the rising slope (mV/ms) of each response was used as a reliable index for extracellular synaptic current. For analysis, 20-60% of the rising slope was calculated for each fEPSP offline and the average of four consecutive responses were recorded for each stimulation intensity.

*Subthreshold stimulation*

To investigate synaptic plasticity, stimulations were reduced to <30% of the maximal fEPSP amplitude attained and applied to the MPP. Stimulations of 5Hz (10 pulses, 200ms intervals, 1 second duration), 20Hz (10 pulses, 50ms intervals, 500ms duration) and 50Hz (10 pulses, 20ms intervals, 200ms duration) were delivered with 12-minute intervals between each frequency stimulation train. The slope of fEPSPs following subthreshold stimulations were recorded and the mean slope values (mV/ms) +/- the standard error of the mean (SEM) were normalized to baseline for plotting. In addition, fEPSP peak amplitudes were analyzed during the delivery of subthreshold stimulations (5Hz, 20Hz, 50Hz). To confirm that recordings had resulted from MPP stimulations (*46*), the cholinergic agonist carbachol (50μM) was added to the aCSF perfusate for 30 minutes post protocol and then washed out. Carbachol depresses the fEPSP responses of granule cells to MPP stimulation in hippocampal slices. Finally, recordings were repeated in the presence of the

$GABA_A$ receptor antagonist picrotoxin (PTX) (50mM). To prevent feedback-driven excessive firing, CA3 was removed before PTX was introduced to the recording chamber.

*Analysis*

Statistical comparisons between groups of different genotypes were performed using ANOVA or t-tests where appropriate in SPSS (IBM Statistics v23). For mixed ANOVA analysis, the Greenhouse-Geisser correction was applied if the sphericity of variance was violated. A significance level of $p<0.05$ was used in all comparisons.

***In vivo electrophysiology and analysis***

All *in vivo* electrophysiology experiments were conducted in accordance with the European Communities Council Directive of 22nd September 2010 (2010/63/EU) for care of laboratory animals and the guidelines of the German Animal Protection Law, and were approved by the North Rhine-Westphalia State Authority (Bezirksamt, Arnsberg).

*Chronic electrode implantation*

Adult male *Prdm8*$^{-/-}$ (n=9) and sex-matched wild type mice (n=9) underwent chronic implantation of electrodes (125 µm diameter, polyurethane-coated stainless-steel wire, BioMedical Instruments, Germany) under anesthesia (intraperitoneal Nembutal, 60 mg/kg, Merial GmbH, Germany) as described previously (*47, 48*). Briefly, screws connected to stainless steel wires served as ground and reference electrodes. Coordinates were based on a stereotactic mouse brain atlas (*49*). A monopolar recording electrode was implanted into the granule cell layer of the DG (2 mm posterior to bregma, 1.6 mm lateral from midline) and a bipolar stimulation electrode in the perforant path (3.8-3.9 mm posterior to bregma, 2.5-3 mm lateral from midline). The assembly was sealed and fixed to the skull with dental acrylic (Paladur®, Heraeus Kulzer GmbH, Germany) after evoked responses in the DG remained stable. Pre- and postoperative analgesia was given by intraperitoneal Meloxicam injection (0.2 mg/kg, Metacam®, Boehringer Ingelheim Vetmedica GmbH, Germany). After surgery mice were housed individually.

*Baseline recordings*

Ten to fourteen days after surgery, experiments were carried out in 20cm (length) x 20cm (width) x 30cm (height) recording chambers where the animals could move freely. Implanted electrodes were connected via a flexible cable and swivel connector to a stimulation unit (World Precision Instruments, USA) and amplifier (A-M Systems, USA). At the beginning of each experiment, the input-output relationship was examined (an evaluation of 8 stimulation intensities from 50 to 400 µA in 50 µA steps). The stimulation intensity that induces an evoked response of ~60% of the maximum evoked response determined during the input-output evaluation was used to evoke responses during the subsequent experiments. The amplitude of the population spike (PS) of the evoked potential was measured from the peak of the first positive deflection to the peak of the following negative deflection. The slope of the fEPSP was calculated between the first deflection point and the second maximum of the potential.

To assess baseline synaptic transmission, responses in the DG were evoked by applying low frequency test pulses (0.025 Hz, biphasic square wave pulses 0.2 ms duration per half wave) in the perforant path. Baseline population spike (PS) and fEPSP values were obtained by averaging evoked responses (5 sweeps at 40s intervals) every 5 minutes for the first 30 minutes. These baseline values were then used as the reference for all subsequent recorded time points, which were calculated as a percentage of the mean of these first six reference values. Further recordings were made every 5 minutes for the next 15 minutes, then every 15 minutes for up to 4.5 hours. After 24 hours, recordings were performed for another hour.

*Pattern stimulation protocols*

Animals that showed stable responses during the first 4.5 hours of recording and for the hour of recording 24 hours later were used for subsequent experiments. High-frequency stimulation (HFS) was applied to induce long term potentiation (LTP) as one or three trains of 6 bursts (each burst containing 6 pulses) delivered at 400 Hz (inter-train interval 20s, the inter-burst interval 200ms).

*Verification of electrode position*

Implanted electrode positions in the DG and perforant path were verified histologically at the end of the recordings following a published protocol for Nissl staining (*50*). Mice with incorrectly implanted electrodes were excluded from further analysis.

*Analysis*

ANOVA with repeated measures was performed to compare the evoked responses in groups of mice of different genotype at baseline or following HFS using Statistica software (TIBCO Software Inc.). All data were expressed as means ± SEM for each genotype.

***Behavioral paradigms***

*Modified SHIRPA*

A modified SHIRPA protocol (*51*) was used as a first-line phenotyping screen of adult *Prdm8*$^{-/-}$ mice (n=11) with their sex- and litter-matched controls (n=10) and adult *Bhlhb4*$^{-/-}$ mice (n=12) with their sex- and litter-matched controls (n=11). Details of the standardized protocol have been published previously (*52*). All mouse cohorts were investigated at the same time during their light cycle to account for changes in behavior due to circadian rhythms. Statistical comparisons were made of continuous measures using t-statistics and presented as the mean ± SEM for each genotype. Categorical data was analyzed using the $C^2$ test. All results were corrected for multiple comparisons.

*Morris Water Maze*

Hippocampus-dependent spatial reference memory was assessed using an open field water maze as described previously (*49*). After habituation, the performance of adult *Prdm8*$^{-/-}$ mice (n=11) was compared with sex- and litter-matched wild type mice (n=10) and with adult *Bhlhb4*$^{-/-}$ mice (n=12) to control for their night blindness phenotype (*39*) All experiments were conducted in the morning, commencing at 0900hours. Briefly, mice were trained to locate a hidden platform (diameter 10cm) submerged approximately 1cm below the surface of a pool (temperature 20-24°C, diameter 1m) with water opacified by white food coloring. Cues were positioned at the North (N, blue circle), West (W, green diamond), South (S, red triangle) and East (E, yellow rectangle) ends of the pool. To control for the possibility of quadrant effects, half of each group of mice were trained to find a platform submerged in the NW quadrant and half were trained to find a platform submerged in the SE quadrant. Each mouse received 4 trials per day for 9 days, with an interval time of approximately 20 seconds. Mice were placed into the pool facing the side wall at one of 8 possible start locations (N, NE, E, SE, S, SW, W, NW), generated randomly across trials by random.org (*53*) and allowed to swim until they found the platform or for a maximum of 60 seconds. Any mouse that failed to find the platform in 60 seconds was placed on the platform for 30 seconds before beginning the next trial. On the 10th day of testing, the platform was removed from the pool and the mice were allowed to swim for 60 seconds. The amount of time that animals spent in each quadrant of the maze was recorded, and the proportion of time spent in the correct quadrant was calculated from this data.

To determine to what extent the visual deficits of *Prdm8*$^{-/-}$ and *Bhlhb4*$^{-/-}$ mice impaired their performance on the water maze compared with wild type controls, they were trained to find a visible platform that was 1cm above water level and clearly indicated by the presence of a beacon (a whiteboard marker pen stuck to the surface of the platform). The position of the visible platform moved clockwise for each trial, although the start location of the mice varied randomly between one of 8 possibilities as described above. The mice received 4 trials per day for 6 days, with an interval time of approximately 20 seconds. Mice were allowed to swim until they found the platform or for a maximum of 60 seconds.

Latency to find the hidden platform (recorded in seconds), distance swam (measured in cm), and the speed of the mice (cm/s) were recorded using a video camera suspended directly above the pool and the Viewpoint videotrack® v3 system (Viewpoint Life Sciences, Montreal, Canada) with Automated Behavioral Analysis for Windows XP software. This software automatically calculates parameters such as speed, trajectory, and position for each animal without bias. Comparisons between groups of different genotypes across training days were made by ANOVA with repeated measures using GraphPad Prism version 5.00 for Windows (GraphPad Software, San Diego California USA). Differences between groups during the probe trial were analyzed by t-test.

*Contextual fear conditioning*

The conditioning context was a stainless-steel conditioning chamber (31 cm × 24 cm × 21 cm; Med Associates, St. Albans, VT) with a stainless steel shock-grid floor. Shock grid bars (diameter 3.2 mm) were spaced 7.9 mm apart. The grid floor was positioned over a stainless-steel drop-pan, which was lightly cleaned with 70% ethyl alcohol to provide background odor. The front, top, and back of the chamber were made of clear acrylic and the two sides made of modular aluminum.

During training, *Prdm8*$^{-/-}$ mice (n=9) and their sex- and litter-matched wild type controls (n=12) were placed in the conditioning chamber and were presented with a single foot shock (0.3mA, 2 seconds) after 2 minutes. They remained in the chamber for an additional minute, then were returned to their home cage. The training was repeated for 4 days. On the fifth day, mice were placed in the same conditioning chamber for 3 minutes without a foot shock. Mouse freezing behavior was monitored via four overhead cameras. Freezing was assessed using an automated scoring system (Actimetrics, Wilmette, IL), which digitized the video signal at 4 Hz and compared movement frame by frame to determine the amount of freezing. An ANOVA with repeated measures was performed to compare the behavior between groups of different genotypes over 5 days and a t-test was used to compare groups during the probe trial on day 5.

*Object Pattern Separation (OPS) Task*

The object pattern separation (OPS) task was used with some modifications (*23*) on adult *Prdm8*$^{-/-}$ mice (n=11) and their sex- and litter-matched (*Prdm8*$^{+/+}$) wild type controls (n=8). Adult *Bhlhb4*$^{-/-}$ mice (n=15) and sex- and litter-matched (*Bhlhb4*$^{+/+}$) wild type mice (n=10) were used as controls for any potential confounding effect of the night blindness phenotype. Experiments were performed in a circular transparent acrylic open topped arena (diameter 40cm, height 50cm) that was purpose-built by the mechanical workshop, University of Bristol, in a brightly lit room. To provide a spatial cue, a black PVC film covered half the circular wall of the arena. This arena was placed within a 1m x 1m x 0.5m wooden frame, surrounded by a black cloth to a height of 1.5m to prevent the visualization of external cues during the experiment, and the floor of the arena was covered in fine saw dust. The cloth on the south wall was used as the entry and exit for animals in the arena and was closed during all phases.

There were 5 objects used for the OPS task made from plastic building blocks (approximate dimensions: 8cm x 4cm x 6cm) which were fixed to a rectangular base using concealed Blue-tack to ensure accurate and consistent object relocation during the task. Objects placed in position 1 on the East and West sides of the arena were positioned symmetrically along the horizontal midline and ~12cm away from the east and west walls, intersecting the division of the transparent and black PVC edges of the arena. Objects placed in position 5 on the East and West sides were placed 3cm away from the adjacent walls to allow for appropriate exploration. Positions 2-4 were predefined at equidistant intervals between positions 1 and 5. For example, position 3 was measured to be halfway between position 1 and position 5. Between each trial, the objects were structurally altered to maintain the exploratory motivation of the mice. Four identical replicates of the objects were used: two for the sample phase and two for the test phase. Between experiments, objects were cleaned with 100% ethanol to remove any confounding odor cues. In addition to this, sawdust was not replaced between sessions but redistributed to remove any odor cues but not affect olfactory saturation (*54*).

Each mouse was exposed to 5 trials of the OPS task over 5 weeks (with a 7-day interval between sessions). Each trial involved a 10-minute sample phase when the mice were allowed to explore two identical objects located in position 1 on the east and west sides of the arena, then returned to their home cages for an hour (as longer time delays significantly impair pattern separation ability in this task)(*23*). After one of the two identical objects was placed along the North to South axis to stay in position 1 or move to another novel predefined location (positions 2-5), the mice were placed back in the arena for a 5-minute test phase. Exposures to positions 1-5 in the task were counterbalanced using pseudo-randomization to ensure that all mice saw every position during the test phase and that these positions were randomly distributed across the West vs East and North vs South axes.

The experimenter monitored and scored mouse behavior in real time from a monitor attached to a mounted overhead camera and using a customized designed MS-DOS programme (created by Prof. Tim Bussey, University of Cambridge) and rescored offline from the VCR recording where necessary.

"Exploration" was defined as nose-directed sniffing <1cm away from an object. Behaviors, such as leaning and/or resting on an object whilst viewing the arena were not classed as "exploration" behavior. Any mouse that failed to explore the objects for more than 10 seconds during the test phase were not included in subsequent analyses.

Analysis of the data was performed offline in Microsoft Excel (2013) and MATLAB R2017a. To determine whether the mice were able to discriminate between the two objects during the test phase, a discrimination ratio (D2) was calculated using the formula below:

$$Discrimination\ Ratio\ (D2) = \frac{D1(Novel\ exploration(s) - familiar\ exploration(s))}{(Novel\ exploration\ (s) + Familiar\ exploration\ (s)}$$

D2 was calculated for each individual mouse and group mean. Discrimination scores between 0 and +1 showed a preference for novel exploration; between 0 and -1 showed a preference for familiar. A discrimination score of zero represented chance (no preference). Additionally, total exploration time in the sample and test phases were calculated to ensure the exploration time was sufficient to allow for appropriate discrimination.

Statistical comparisons of normally distributed data were made by ANOVA using SPSS and a Tukey's post hoc test for multiple comparisons.

*Novel Object Recognition (NOR)*

The Novel Objection Recognition (NOR) task was performed on adult *Prdm8*$^{-/-}$ mice (n=14) and their sex- and litter-matched (*Prdm8*$^{+/+}$) wild type controls (n=13). Adult *Bhlhb4*$^{-/-}$ mice (n=15) and sex- and litter-matched (*Bhlhb4*$^{+/+}$) wild type mice (n=10) were used as controls for any potential confounding effect of the night blindness phenotype. Experiments were performed in the same circular arena described above for the OPS task.

Following the habituation period, each animal was placed in the arena for 10 minutes and allowed to explore two identical objects, $A^1$ and $A^2$, then returned to their home cage for 1 hour. One object was then replaced with a novel object B and the other object was replaced with an identical object $A^3$ to those presented in the sample phase, which served as the 'familiar' object. To control object bias, the object that was the familiar object for half of the cohort, served as the novel for the remainder (*55*). To control positional bias, half of the cohort saw the novel object on the left, whereas the other half saw it on the right. In so doing, exposures were counterbalanced throughout (*56*). In total, 4 identical replicates of the objects were used: two for the sample phase and two for the test phase. During the test phase, each animal was placed back in the arena and allowed to explore both objects freely for 5 minutes.

Exploration of the objects in a familiar vs novel location was defined, scored and analysed as for the OPS task. As before, statistical comparisons of normally distributed data were made by ANOVA using SPSS. A post-hoc Bonferonni correction was applied for multiple comparisons. To compare test scores with chance, a one sample t-test was used.

**Induction of epilepsy**

Six litter-matched pairs of adult male *Prdm8*$^{-/-}$ and wild type mice were placed in individual Plexiglass cylinders (60csm height x 30cm diameter) for observation. The behavior of the mice was recorded using a mounted video camera as above. Each mouse was then given an intraperitoneal injection of kainic acid (30mg/kg, Hello Bio HB0355) to induce seizures and their behavior was video recorded for the following 2 hours. Behavioral seizures were scored every 5 minutes in real time by the experimenter according to a modified Racine scale (*54*). This scale consisted of seven stages, ranging from zero to seven, with seven being the most severe (Table S38). The most severe seizure stage observed during each 5-minute interval was the recorded score. The video recordings were then scored by a second independent observer who was blind to the genotype of the mice and the scores were compared for interobserver variability. One mouse with severe continuous seizures and another with irreversible status epilepticus status were given anti-seizure medication (diazepam) to stop the seizures after 100 minutes in accordance with the endpoints described in the Home Office project license. All mice were euthanized at the end of the 2-hour observation period.

**Neural mass modelling of the dentate gyrus**

We implemented a neural mass model like the model proposed by López-Cuevas *et al* (*28*). The dynamics of the DG was described by the following first-order differential equations:

$$\dot{v}_{gc} = x_{gc}$$

$$\dot{x}_{gc} = Aa\{P_1(t) + s(t) + S(C_1 v_{mc} - C_2 v_{bc} - C_3 v_{hc})\} - 2a x_{gc} - a^2 v_{gc}$$

$$\dot{v}_{mc} = x_{mc}$$

$$\dot{x}_{mc} = AaS\big(C_4 v_{gc}\big) - 2a x_{mc} - a^2 v_{mc}$$

$$\dot{v}_{bc} = x_{bc}$$

$$\dot{x}_{bc} = GgS\big(C_5 v_{gc} + C_6 v_{mc}\big) - 2g x_{bc} - g^2 v_{bc}$$

$$\dot{v}_{hc} = x_{hc}$$

$$\dot{x}_{hc} = Bb\{P_2(t) + S\big(C_7 v_{gc}\big)\} - 2bx_{hc} - b^2 v_{hc}$$

where $v_{gc}$, $v_{mc}$, $v_{bc}$, and $v_{hc}$ are the output potentials of the neuronal populations (granule cells (GC), mossy cells (MC), basket cells (BC), and HIPP cells (HC), respectively). $x_{gc}$, $x_{mc}$, $x_{bc}$, and $x_{hc}$ are auxiliary variables, $P_1(t)$ and $P_2(t)$ represent inputs to the circuit delivered by the perforant path (PP), $s(t)$ accounts for external stimulation delivered to the GC, $S$ is a sigmoid function,

$$S = \frac{2e_0}{1 + e^{r(v_0 - v)}}$$

and $A$, $a$, $G$, $g$, $B$, $b$, $C_1$-$C_7$, $e_0$, $r$ and $v_0$ are parameters (see Table S37 for their biophysical meaning and values). The underlying circuitry was based on Danielson *et al* which had parameters that could be translated to a neural mass model (*21*). For simplicity, $P_1(t)$ and $P_2(t)$ were modelled as two independent noise generators (Gaussian noise, zero mean, standard deviation $\sigma = p/\sqrt{dt}$, where $dt$ is the integration time step and $p$ is a parameter). The external stimulation $s(t)$ was used to induce seizure-like activity in the DG. We integrated the equations using the Euler-Maruyama method with $dt = 0.001$ s. Using the parameters in Table S37, we observed that $s(t)$ elicits seizure-like activity when $s(t) > 0.04$. At sufficiently large stimulation values, $s(t) > 1$, the BC activity saturates and suppresses seizure-like activity in the circuit. To model the removal of the connections from MC to GC we set $C_1 = 0$, whereas the removal of the connections from MC to BC corresponded to $C_6 = 0$. The removal of MC from the circuit was modelled as $C_1 = C_6 = 0$.

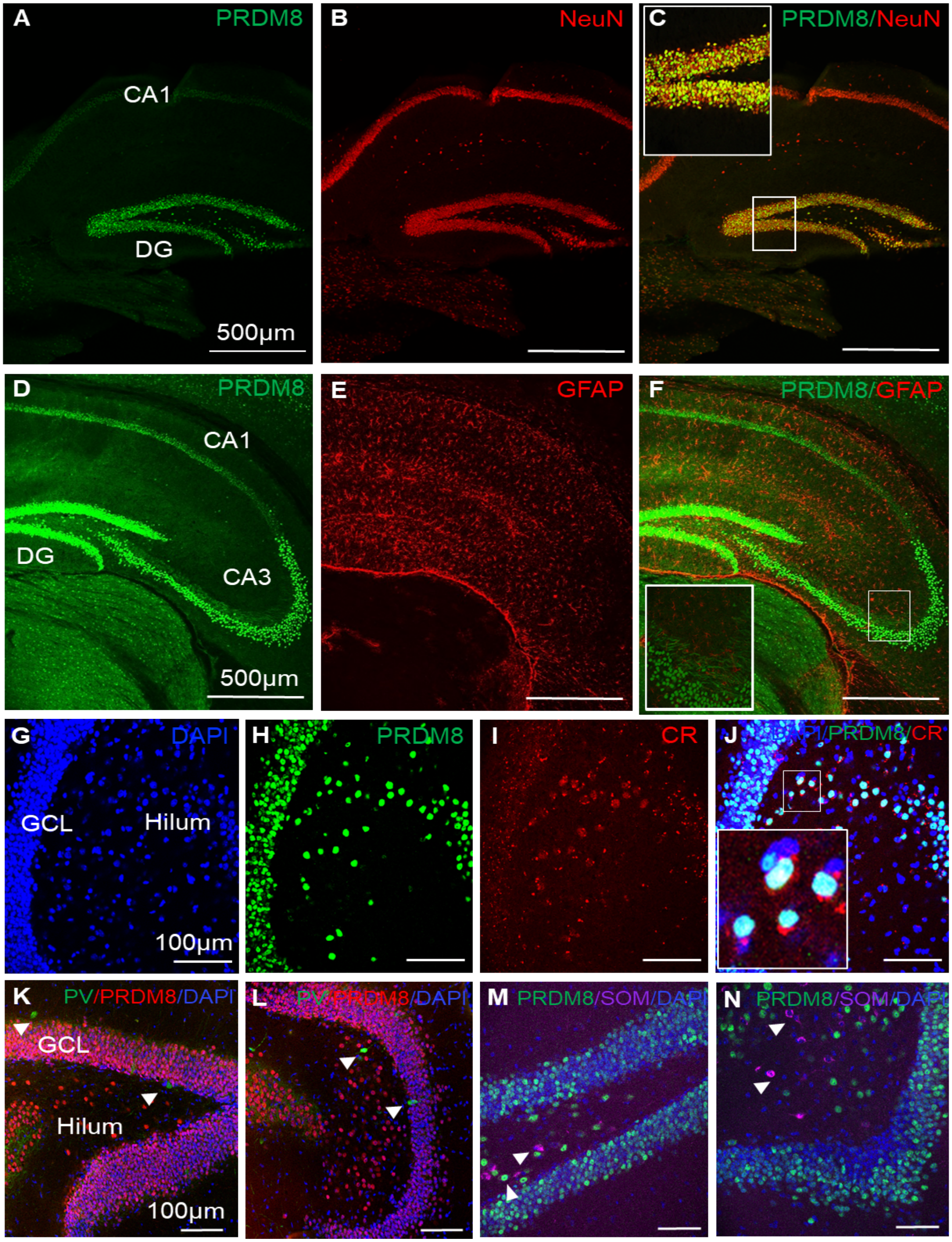
A PRDM8
CA1
DG
500μm
B NeuN
C PRDM8/NeuN
D PRDM8
CA1
DG
CA3
500μm
E GFAP
F PRDM8/GFAP
G DAPI
GCL
Hilum
100μm
H PRDM8
I CR
J DAPI/PRDM8/CR
K PV/PRDM8/DAPI
GCL
Hilum
100μm
L PV/PRDM8/DAPI
M PRDM8/SOM/DAPI
N PRDM8/SOM/DAPI

374

**Fig. S1.**

**PRDM8 is expressed in principal neuron populations within the adult mouse hippocampus.** (A-N) Representative confocal micrographs of sections from adult wild type hippocampus co-stained with antibodies to PRDM8 and cell-type specific markers. (A–C) PRDM8 was expressed in $NeuN^+$ principal neurons within the pyramidal cell layers of CA1 and CA3, and the granule cell layer (GCL) and hilar region of the DG (C inset magnified 4x). (D-F) PRDM8 was not expressed by $GFAP^+$ (Glial Fibrillary Acidic Protein) astrocytes in the DG, CA1 or CA3 (F inset magnified 4x). (G-J) In the DG, PRDM8 was expressed in granule cells in the GCL and $calretinin^+$ (CR) mossy cells in the hilum (J inset magnified 4x) but not in (K, L) parvalbumin+ (PV) or (M, N) Somatostatin+ (SOM) interneurons (white arrowheads point to single-labelled interneurons).

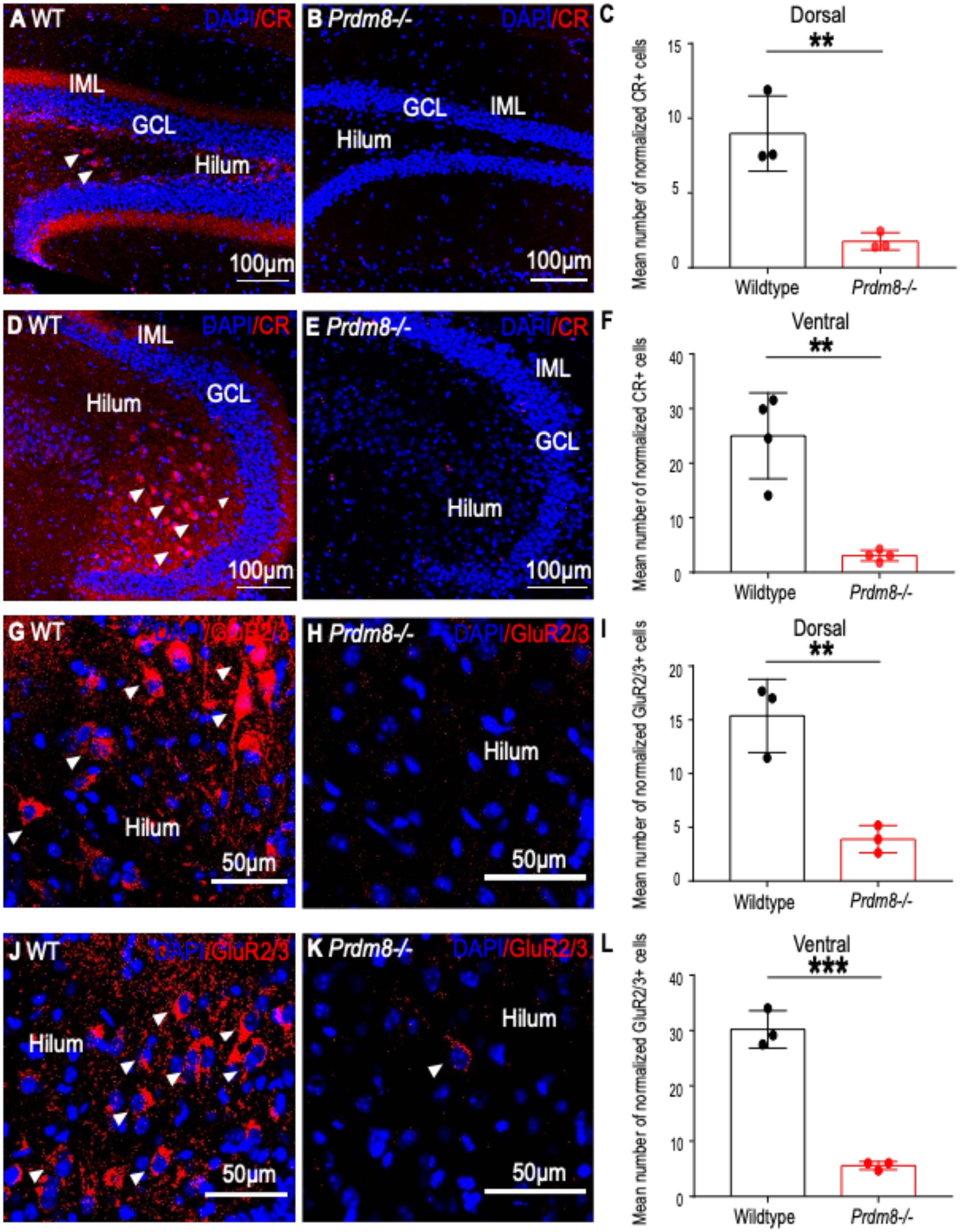
A WT
DAPI/CR
IML
GCL
Hilum
100µm
B Prdm8-/-
DAPI/CR
GCL
IML
Hilum
100µm
C
Dorsal
**
Mean number of normalized CR+ cells
15
10
5
0
Wildtype
Prdm8-/-
D WT
IML
DAPI/CR
Hilum
GCL
100µm
E Prdm8-/-
DAPI/CR
IML
GCL
Hilum
100µm
F
Ventral
**
Mean number of normalized CR+ cells
40
30
20
10
0
Wildtype
Prdm8-/-
G WT
DAPI/GluR2/3
Hilum
50µm
H Prdm8-/-
DAPI/GluR2/3
Hilum
50µm
I
Dorsal
**
Mean number of normalized GluR2/3+ cells
20
15
10
5
0
Wildtype
Prdm8-/-
J WT
DAPI/GluR2/3
Hilum
50µm
K Prdm8-/-
DAPI/GluR2/3
Hilum
50µm
L
Ventral
***
Mean number of normalized GluR2/3+ cells
40
30
20
10
0
Wildtype
Prdm8-/-

386

**Fig. S2.**

**Mossy cells are absent from the *Prdm8*$^{-/-}$ hippocampal dentate gyrus (DG).** (A-L) Representative confocal micrographs of sections from adult *Prdm8*$^{-/-}$ and wild type (WT) mice of dorsal and ventral DG labelled with antibodies to calretinin (CR) and GluR2/3, counterstained with DAPI (4'-6-diamidino-2-dihydrochloride). (A-F) CR labels mossy cell (MC) bodies (white arrowheads) and axon projections to the inner molecular layer (IML) (red band) of dorsal (A) and ventral DG (D). CR$^{+}$ mossy cell bodies and axon projections were absent from the DG of *Prdm8*$^{-/-}$ mice and CR$^{+}$ cell counts were significantly reduced compared with the (A-C) dorsal (n=3|3, p=0.009) and (D-F) ventral (n=4|4, p=0.002) DG of WT mice. Note: CR also labels some interneurons and immature granule cells (GCs). (G-L) GluR2/3 is an alternative marker of MCs; GluR2/3 labels MC bodies in the DG hilum but not their axon projections (G, J). GluR2/3 is also expressed by mature GCs in the granule cell layer (GCL). GluR2/3 expression was virtually absent from the *Prdm8*$^{-/-}$ DG hilum and GluR2/3$^{+}$ hilar cell counts were significantly reduced compared with the (G-I) dorsal (n=3|3, p=0.006) and (J-L) ventral DG (n=3|3, p<0.001) of WT controls. Statistical comparisons are summarized in Table S1. **=p<0.01; ***=p<0.001.

403 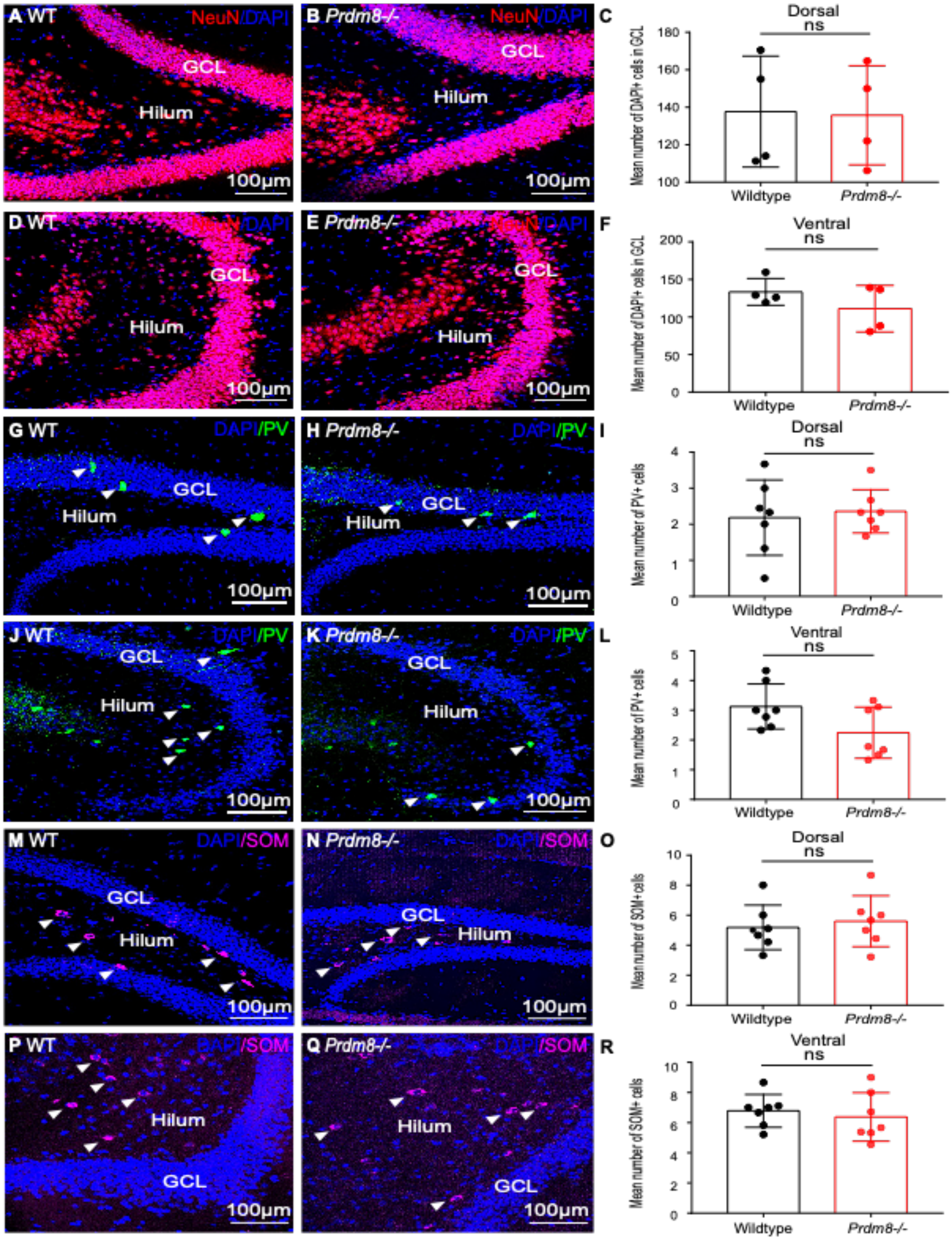
A WT
NeuN/DAPI
GCL
Hilum
100μm
B Prdm8-/-
NeuN/DAPI
GCL
Hilum
100μm
C
Dorsal
ns
Mean number of DAPI+ cells in GCL
180
160
140
120
100
Wildtype
Prdm8-/-
D WT
GCL
Hilum
100μm
E Prdm8-/-
GCL
Hilum
100μm
F
Ventral
ns
Mean number of DAPI+ cells in GCL
200
150
100
50
0
Wildtype
Prdm8-/-
G WT
DAPI/PV
GCL
Hilum
100μm
H Prdm8-/-
DAPI/PV
GCL
Hilum
100μm
I
Dorsal
ns
Mean number of PV+ cells
4
3
2
1
0
Wildtype
Prdm8-/-
J WT
DAPI/PV
GCL
Hilum
100μm
K Prdm8-/-
DAPI/PV
GCL
Hilum
100μm
L
Ventral
ns
Mean number of PV+ cells
5
4
3
2
1
0
Wildtype
Prdm8-/-
M WT
DAPI/SOM
GCL
Hilum
100μm
N Prdm8-/-
DAPI/SOM
GCL
Hilum
100μm
O
Dorsal
ns
Mean number of SOM+ cells
10
8
6
4
2
0
Wildtype
Prdm8-/-
P WT
DAPI/SOM
Hilum
GCL
100μm
Q Prdm8-/-
DAPI/SOM
Hilum
GCL
100μm
R
Ventral
ns
Mean number of SOM+ cells
10
8
6
4
2
0
Wildtype
Prdm8-/-

**Fig. S3.**

**Loss of *Prdm8* function does not affect the granule cell layer (GCL) or major interneuron populations in the adult mouse hippocampal dentate gyrus (DG).** (A-R) Representative confocal micrographs of sections from adult *Prdm8*$^{-/-}$ and wild type (WT) mice of the dorsal and ventral DG labelled with antibodies to NeuN, Parvalbumin (PV) and Somatostatin (SOM) and counterstained with DAPI (4'-6-diamidino-2-dihydrochloride). (A-F) There were no differences in NeuN and DAPI staining of the (A,B) dorsal or (D,E) ventral DG of *Prdm8*$^{-/-}$ mice compared with WT. (C,F) Counts of the number of DAPI$^{+}$ neurons in the (C) dorsal (n=4|4, $p>0.05$) and (F) ventral GCL (n=4|4, $p>0.05$) were no different in *Prdm8*$^{-/-}$ mice compared with WT controls. (G-L) Additionally, there were no significant differences in the staining and number of PV$^{+}$ interneurons (white arrow heads) in the (G-I) dorsal (n=7|7, $p>0.05$) and (J-L) ventral DG (n=7|7, $p>0.05$) of *Prdm8*$^{-/-}$ mice compared with WT controls. (M-R) There were no significant differences in the staining or number of SOM$^{+}$ interneurons (white arrow heads) in the (M-O) dorsal (n=7|7, $p>0.05$) and (P-R) ventral DG (n=7|7, $p>0.05$) of *Prdm8*$^{-/-}$ mice compared with WT controls. Statistical comparisons are summarized in Table S1. n.s = non-significant.

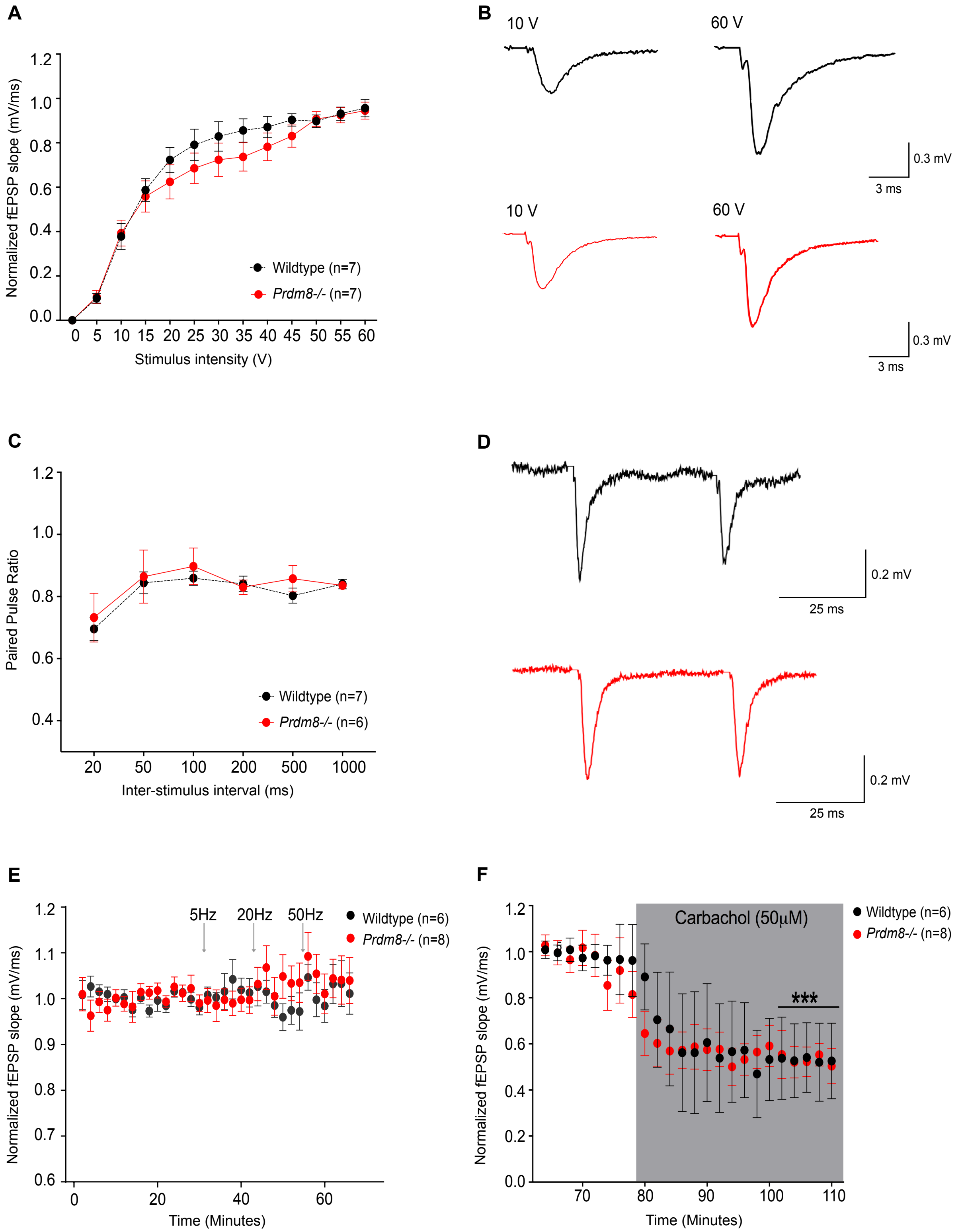
A
Normalized fEPSP slope (mV/ms)
1.2
1.0
0.8
0.6
0.4
0.2
0.0
0 5 10 15 20 25 30 35 40 45 50 55 60
Stimulus intensity (V)
Wildtype (n=7)
*Prdm8-/-* (n=7)
B
10 V
60 V
0.3 mV
3 ms
10 V
60 V
0.3 mV
3 ms
C
Paired Pulse Ratio
1.2
1.0
0.8
0.6
0.4
20 50 100 200 500 1000
Inter-stimulus interval (ms)
Wildtype (n=7)
*Prdm8-/-* (n=6)
D
0.2 mV
25 ms
0.2 mV
25 ms
E
Normalized fEPSP slope (mV/ms)
1.2
1.1
1.0
0.9
0.8
0.7
0.6
0 20 40 60
Time (Minutes)
5Hz
20Hz
50Hz
Wildtype (n=6)
*Prdm8-/-* (n=8)
F
Normalized fEPSP slope (mV/ms)
1.2
1.0
0.8
0.6
0.4
0.2
0.0
70 80 90 100 110
Time (Minutes)
Carbachol (50μM)
***
Wildtype (n=6)
*Prdm8-/-* (n=8)

**Fig. S4.**

**The absence of mossy cells from *$Prdm8^{-/-}$* dentate gyrus (DG) does not influence basal granule cell (GC) responses or short-term plasticity to medial perforant pathway (MPP) stimulation *in vitro*.** (A) There were no differences between genotypes (n=7|7) in the normalized slope values of field excitatory post-synaptic potentials (fEPSPs) in GC responses to increasing (0-60V) MPP stimulation (p=0.323); (B) example waveforms after 10V and 60V stimulation in WT (black lines) and *$Prdm8^{-/-}$* (red lines) acute hippocampal slices. (C) Paired pulse (PP) depression (PP ratio (PPR)<1) was evident in GC responses to stimulation at all interstimulus intervals (20-1000ms), confirming the electrode was stimulating MPP not lateral perforant path (LPP). There were no differences in PPR between genotypes (n=6|7)(p=0.872); (D) example waveforms after MPP stimulation at 50ms intervals in WT (black lines) and *$Prdm8^{-/-}$* (red lines) slices. (E) The normalized slope values of fEPSPs recorded in GCs for 12 minutes after 5, 20 and 50Hz stimulations were no different between genotypes (n=8|6)(p=0.896). (F) GC responses were depressed when 50μM carbachol was added to the perfusate during minutes 80-110, confirming the electrode was stimulating MPP not LPP in *$Prdm8^{-/-}$* (p<0.001) and WT slices (p<0.001)(n=8|6). Statistical comparisons are summarized in Tables S2-5. *** =p<0.001.

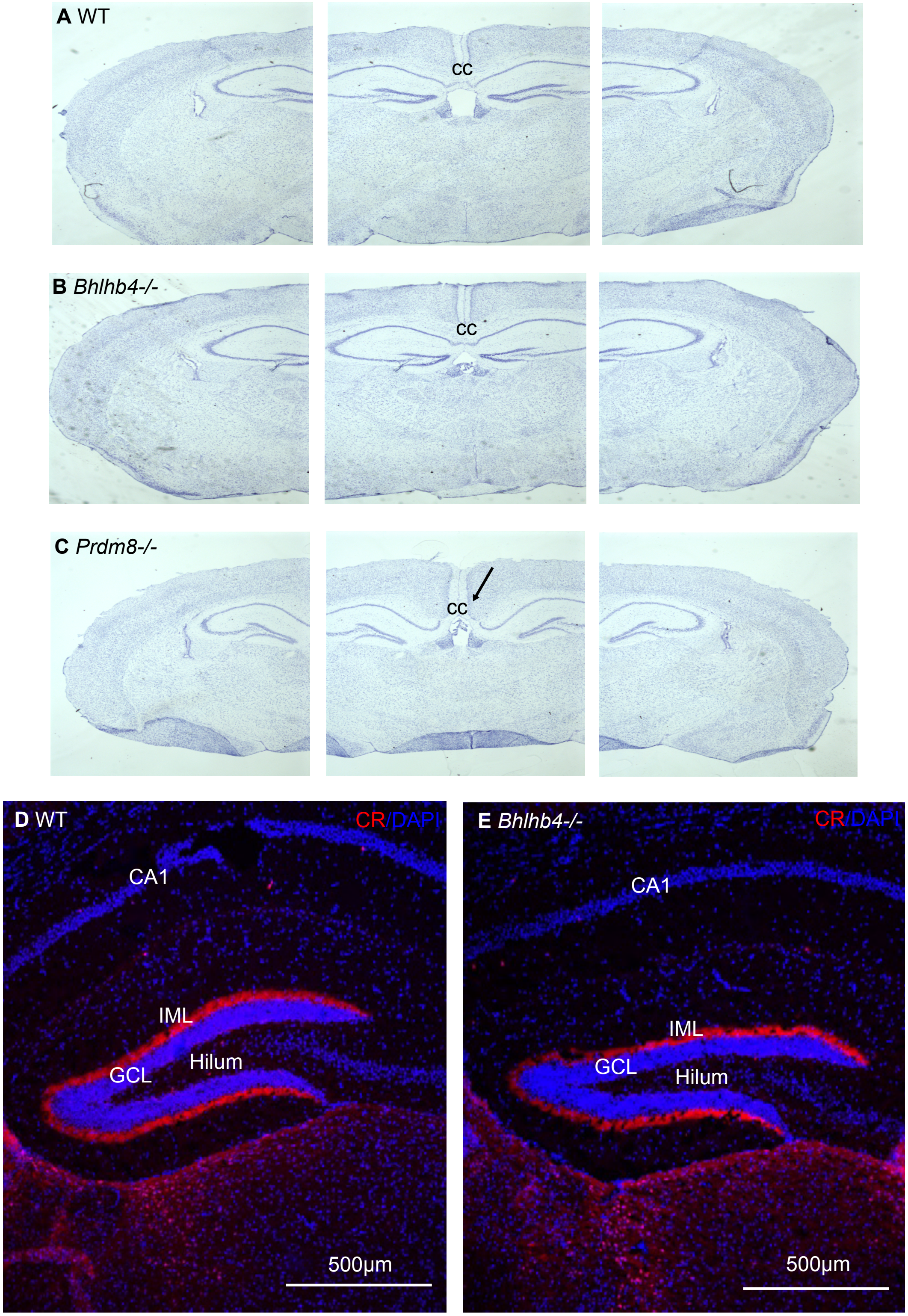
A WT
cc
B Bhlhb4-/-
cc
C Prdm8-/-
cc
D WT
CR/DAPI
CA1
IML
Hilum
GCL
500µm
E Bhlhb4-/-
CR/DAPI
CA1
IML
GCL
Hilum
500µm

**Fig. S5.**

**Mossy cells are not absent from the *Bhlhb4*$^{-/-}$ hippocampal dentate gyrus.** (A-C) Representative coronal sections from (A) wild type (WT), (B) *Bhlhb4*$^{-/-}$ and (C) *Prdm8*$^{-/-}$ mice after stained with Nissl to show the corpus callosum and hippocampal commissure. These structures were absent in (C) although the phenotype was not fully penetrant in *Prdm8*$^{-/-}$ mice: only 1 of 4 *Prdm8*$^{-/-}$ mice had evidence of this phenotype. (D, E) Representative confocal micrographs of sections from the dorsal hippocampal dentate gyrus (DG) of WT and *Bhlhb4*$^{-/-}$ mice labelled with antibodies to Calretinin (CR) and counterstained with DAPI (4’-6-diamidino-2-dihydrochloride). CR$^+$ mossy cell bodies and axonal projections to the inner molecular layer (IML) were present in both WT and *Bhlhb4*$^{-/-}$ mouse hippocampus.

A

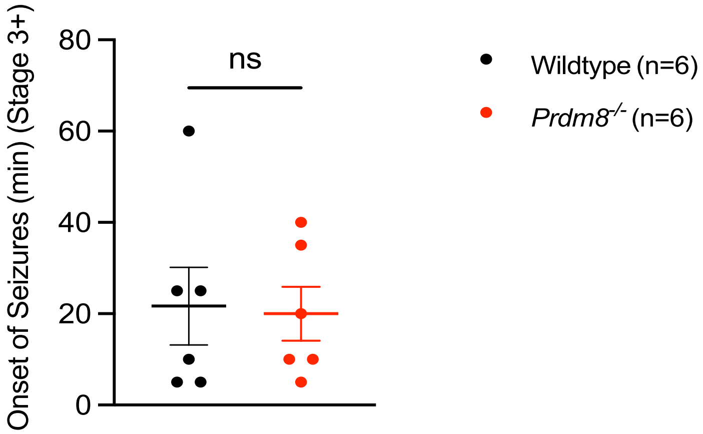
Onset of Seizures (min) (Stage 3+)
80
60
40
20
0
ns
Wildtype (n=6)
Prdm8-/- (n=6)

B

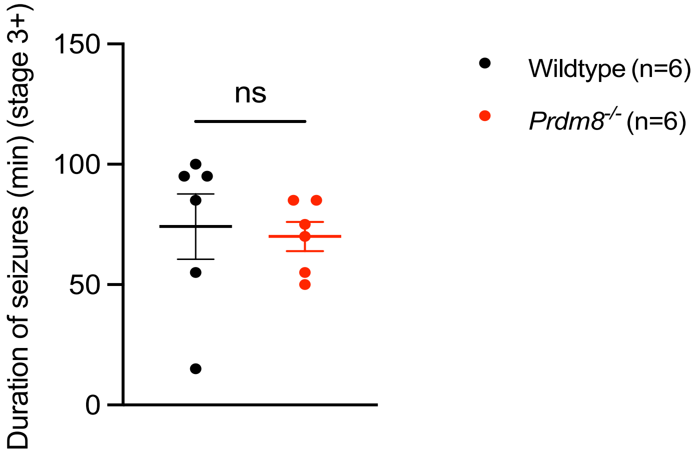
Duration of seizures (min) (stage 3+)
150
100
50
0
ns
Wildtype (n=6)
Prdm8-/- (n=6)

**Fig. S6.**

**The onset and duration of seizures induced by intraperitoneal injection of kainic acid are no different in wild type and *Prdm8$^{-/-}$* mice.** (A, B) There were no differences between wild type and *Prdm8$^{-/-}$* mice (n=6|6) in (A) the time of onset of modified Racine stage 3+ seizures (in minutes) (p=0.876) or (B) the duration of modified Racine stage 3+ seizures (in minutes) (p=0.785) Statistical comparisons are summarized in Table S40. n.s = non-significant.

**Table S1**

**Quantification of cell populations in the *Prdm8*$^{-/-}$ and wild type dentate gyrus.** Cells labelled with DAPI (4’-6-diamidino-2-dihydrochloride) and antibodies to parvalbumin, somatostatin, GluR2/3 and calretinin were counted in the dorsal and ventral DG of *Prdm8*$^{-/-}$ and litter-matched wild type animals (n=3-7), then compared using the two-tailed unpaired t-test. GluR2/3 and calretinin cell counts were significantly lower in the dorsal and ventral DG of *Prdm8*$^{-/-}$ mice compared with wild type (WT). N=number; SEM=standard error of the mean; df=degrees of freedom.

| **Cell marker** | **Group** | | **N** | **Mean ± SEM** | **t value, df** | | **p value** | |
|---|---|---|---|---|---|---|---|---|
| | | | | | **Dorsal** | **Ventral** | **Dorsal** | **Ventral** |
| Calretinin | WT | Dorsal | 3 | 8.98±1.46 | t=4.828 df=4 | t=5.538 df=6 | 0.009 | 0.002 |
| | | Ventral | 4 | 25.02±3.93 | | | | |
| | *Prdm8*$^{-/-}$ | Dorsal | 3 | 1.76±0.34 | | | | |
| | | Ventral | 4 | 3.06±0.49 | | | | |
| GluR2/3 | WT | Dorsal | 3 | 15.37±1.97 | t=5.463 df=4 | t=12.33 df=4 | 0.006 | <0.001 |
| | | Ventral | 3 | 30.21±1.95 | | | | |
| | *Prdm8*$^{-/-}$ | Dorsal | 3 | 3.90±0.74 | | | | |
| | | Ventral | 3 | 5.58±0.42 | | | | |
| DAPI | WT | Dorsal | 4 | 137.71±14.81 | t=0.099 df=6 | t=1.248 df=6 | 0.925 | 0.258 |
| | | Ventral | 4 | 133.46±8.91 | | | | |
| | *Prdm8*$^{-/-}$ | Dorsal | 4 | 135.75±13.21 | | | | |
| | | Ventral | 4 | 111.13±15.51 | | | | |
| Parvalbumin | WT | Dorsal | 7 | 2.18±0.39 | t=0.384 df=12 | t=2.03 df=12 | 0.708 | 0.065 |
| | | Ventral | 7 | 3.13±0.29 | | | | |
| | *Prdm8*$^{-/-}$ | Dorsal | 7 | 2.36±0.23 | | | | |
| | | Ventral | 7 | 2.25±0.33 | | | | |
| Somatostatin | WT | Dorsal | 7 | 5.19±0.56 | t=0.484 df=12 | t=0.552 df=12 | 0.637 | 0.591 |
| | | Ventral | 7 | 6.79±0.41 | | | | |
| | | Dorsal | 7 | 5.60±0.64 | | | | |

| **Cell marker** | **Group** | | **N** | **Mean ± SEM** | **t value, df** | | **p value** | |
|---|---|---|---|---|---|---|---|---|
| | | | | | **Dorsal** | **Ventral** | **Dorsal** | **Ventral** |
| | *Prdm8*$^{-/-}$ | Ventral | 7 | 6.38±0.61 | | | | |

**Table S2**

**Statistical comparisons of input/output granule cell (GC) responses to increasing medial perforant path (MPP) stimulation of *Prdm8*$^{-/-}$ and wild type hippocampal slices *in vitro* (n=7|7).** Mixed ANOVA was used to compare mean field excitatory postsynaptic potential (fEPSP) slope values in response to increasing MPP stimulation from 0 to 60V (mV/ms) but there were no statistically significant differences between genotypes. SS=sum of squares; MS=mean square; df=degrees of freedom

| | **SS** | **df** | **MS** | **F** | **p value** |
|---|---|---|---|---|---|
| Stimulus intensity | 16.154 | 2.330 | 6.933 | 169.421 | <0.001 |
| Genotype | 0.100 | 1 | 0.100 | 0.696 | 0.420 |
| Stimulus intensity*Genotype | 0.114 | 2.330 | 0.049 | 1.196 | 0.323 |
| Stimulus Intensity Error | 1.144 | 27.958 | 0.041 | | |
| Genotype Error | 1.718 | 12 | 0.143 | | |

**Table S3**

**Statistical comparisons of paired pulse relationship of the mean field excitatory postsynaptic potential (fEPSP) slope values recorded in granule cells in response to paired pulse stimulation of the medial perforant path (MPP) in wild type and *Prdm8*$^{-/-}$ hippocampal slices *in vitro* (n=7|6).** Mixed ANOVA was used to compare the paired pulse ratio of mean fEPSP slope values (mV/ms) recorded in granule cells in response to paired pulse stimulations of the MPP at increasing time intervals (20ms, 50ms, 100ms, 200ms, 500ms and 1000ms) in wild type vs *Prdm8*$^{-/-}$ hippocampal slices but there were no statistical differences between genotypes. SS=sum of squares; MS=mean square; df=degrees of freedom.

| | **SS** | **df** | **MS** | **F** | **p value** |
|---|---|---|---|---|---|
| Paired Pulse Interval | 0.184 | 2.124 | 0.087 | 5.740 | 0.010 |
| Genotype | <0.001 | 1 | <0.001 | <0.001 | 1.000 |
| Paired Pulse Interval * Genotype | 0.005 | 2.124 | 0.002 | 0.151 | 0.872 |
| Paired Pulse Interval Error | 0.289 | 19.119 | 0.015 | | |
| Genotype Error | 0.340 | 9 | 0.038 | | |

**Table S4**

**Statistical comparisons of mean field excitatory postsynaptic potential (fEPSP) slope values recorded in granule cells in response to stimulation of the medial perforant path (MPP) at 5Hz, 20Hz and 50Hz in *Prdm8*$^{-/-}$ and wild type hippocampal slices *in vitro* (n=8|6).** Mixed ANOVA was used to compare baseline fEPSP slope values (mV/ms) of granule cell responses to 5Hz, 20Hz and 50Hz subthreshold stimulations for 12 minutes following induction but there were no statistically significant differences between genotypes. SS=sum of squares; MS=mean square; df=degrees of freedom.

| | SS | df | MS | F | p value |
|---|---|---|---|---|---|
| Time x Genotype | 0.122 | 32 | 0.004 | 0.694 | 0.896 |
| Time | 0.152 | 32 | 0.005 | 0.861 | 0.687 |
| Genotype | 0.011 | 1 | 0.011 | 0.177 | 0.682 |
| Residual | 2.117 | 384 | 0.006 | | |

**Table S5**

**Statistical comparisons of mean field excitatory postsynaptic potential (fEPSP) slope values recorded in granule cells in response to medial perforant path (MPP) stimulation before and after the application of carbachol *in vitro* to *Prdm8*$^{-/-}$ and wild type (WT) hippocampal slices (n=8|6).** Using the 2-tailed paired t-test, fEPSP slope values (mV/ms) were found to be significantly depressed after the application of carbachol (50µM) to wild type and *Prdm8*$^{-/-}$ hippocampal slices compared with baseline responses, confirming that the stimulation electrode was stimulating the MPP and not the lateral perforant path. SD=standard deviation; SE=standard error; df=degrees of freedom.

| Genotype | Condition | Paired Differences | | | | | t | df | p value |
|---|---|---|---|---|---|---|---|---|---|
| | | Mean | SD | SE | 95% Confidence Intervals | | | | |
| | | | | | Lower | Upper | | | |
| WT | Baseline vs carbachol | 0.458 | 0.020 | 0.008 | 0.437 | 0.479 | 56.022 | 5 | <0.001 |
| *Prdm8*$^{-/-}$ | Baseline vs carbachol | 0.433 | 0.046 | 0.019 | 0.385 | 0.482 | 22.874 | 5 | <0.001 |

**Table S6**

**Within group statistical comparisons of short-term dynamics of medial perforant path-granule cell synapses to 5Hz (n=13|10), 20Hz (n=8|6) and 50Hz (n=8|6) stimulation in *Prdm8*$^{-/-}$ and WT hippocampal slices *in vitro*.** Normalized peak amplitudes were compared within-groups by two-way mixed ANOVA. The Greenhouse-Geisser correction (epsilon<0.75) was applied where Maulchy's test for Sphericity significance was<0.05. There were no significant interactions between stimulation frequency and group. SS=sum of squares; MS=mean square; df=degrees of freedom.

| Tests of Within-Subjects Effects | | | | | | | |
|---|---|---|---|---|---|---|---|
| **Frequency** | **Source** | **SS** | **df** | **MS** | **F** | **p value** | **Partial Eta Squared** |
| 5Hz WT vs *Prdm8*$^{-/-}$ | Stim | 1.045 | 1.828 | 0.572 | 109.019 | <0.001 | 0.838 |
| | stim*group | 0.007 | 1.828 | 0.004 | 0.729 | 0.477 | 0.034 |
| | Error (stim) | 0.201 | 38.396 | 0.005 | | | |
| 20Hz WT vs *Prdm8*$^{-/-}$ | Stim | 2.523 | 2.822 | 0.894 | 212.232 | <0.001 | 0.946 |
| | stim*group | 0.007 | 2.822 | 0.003 | 0.622 | 0.596 | 0.049 |
| | Error (stim) | 0.143 | 33.867 | 0.004 | | | |
| 50Hz WT vs *Prdm8*$^{-/-}$ | Stim | 2.629 | 2.344 | 1.122 | 134.906 | <0.001 | 0.918 |
| | stim*group | 0.060 | 2.344 | 0.025 | 3.064 | 0.055 | 0.203 |
| | Error (stim) | 0.234 | 28.126 | 0.008 | | | |

**Table S7**

**Between group statistical comparisons of short-term dynamics of medial perforant path-granule cell synapses to 5Hz (n=13|10), 20Hz (n=8|6) and 50Hz (n=8|6) stimulation in *Prdm8*$^{-/-}$ and WT hippocampal slices *in vitro*.** Normalized peak amplitudes were compared between-groups by two-way mixed ANOVA. There was no difference between genotypes in granule cell responses to 5Hz stimulation, but there were significant differences between genotypes to stimulations at 20Hz and 50Hz. SS=sum of squares; MS=mean square; df=degrees of freedom.

| **Tests of Between-Subjects Effects** | | | | | | | |
|---|---|---|---|---|---|---|---|
| **Frequency** | **Source** | **SS** | **df** | **MS** | **F** | **p value** | **Partial Eta Squared** |
| 5Hz WT vs *Prdm8*$^{-/-}$ | Intercept | 95.947 | 1 | 95.947 | 944.794 | <0.001 | 0.978 |
| | group | 0.010 | 1 | 0.010 | 0.096 | 0.760 | 0.005 |
| | Error | 2.133 | 21 | 0.102 | | | |
| 20Hz WT vs *Prdm8*$^{-/-}$ | Intercept | 36.057 | 1 | 36.057 | 972.534 | <0.001 | 0.988 |
| | group | 0.403 | 1 | 0.403 | 10.875 | 0.006 | 0.475 |
| | Error | 0.445 | 12 | 0.037 | | | |
| 50Hz WT vs *Prdm8*$^{-/-}$ | Intercept | 23.645 | 1 | 23.645 | 303.203 | <0.001 | 0.962 |
| | group | 0.514 | 1 | 0.514 | 6.592 | 0.025 | 0.355 |
| | Error | 0.936 | 12 | 0.078 | | | |

**Table S8**

**Within group statistical analysis of short-term dynamics of medial perforant path-granule cell synapses to 5Hz (n=13|9), 20Hz (n=8|9) and 50Hz (n=8|9) *in vitro* stimulations in *Prdm8*$^{-/-}$ slices compared with wild type (WT) hippocampal slices in the presence of picrotoxin (PTX).** Normalized peak amplitudes were compared within groups by two-way mixed ANOVA with Greenhouse-Geisser correction. There was a significant interaction at 50Hz. SS=sum of squares; MS=mean square; df=degrees of freedom.

| **Tests of Within-Subjects Effects** | | | | | | | |
|---|---|---|---|---|---|---|---|
| **Frequency** | **Source** | **SS** | **df** | **MS** | **F** | **p value** | **Partial Eta Squared** |
| 5Hz *Prdm8*$^{-/-}$ vs WT+PTX | stim | 0.782 | 2.573 | 0.304 | 85.918 | <0.001 | 0.811 |
| | stim*group | 0.006 | 2.573 | 0.002 | 0.666 | .555 | 0.032 |
| | Error (stim) | 0.182 | 51.466 | 0.004 | | | |
| 20Hz *Prdm8*$^{-/-}$ vs WT+PTX | stim | 2.995 | 1.387 | 2.159 | 62.232 | <0.001 | 0.806 |
| | stim*group | 0.018 | 1.387 | 0.013 | 0.371 | 0.618 | 0.024 |
| | Error (stim) | 0.722 | 20.807 | 0.035 | | | |
| 50Hz *Prdm8*$^{-/-}$ vs WT+PTX | stim | 6.608 | 1.367 | 4.832 | 28.560 | <0.001 | 0.656 |
| | stim*group | 1.935 | 1.367 | 1.415 | 8.365 | 0.005 | 0.358 |
| | Error (stim) | 3.470 | 20.512 | 0.169 | | | |

**Table S9**

**Between group statistical analysis of short-term dynamics of medial perforant path-granule cell synapses to 5Hz (n=13|9), 20Hz (n=8|9) and 50Hz (n=8|9) *in vitro* stimulations in *Prdm8$^{-/-}$* slices compared with wild type hippocampal slices in the presence of picrotoxin (PTX).** Normalized peak amplitudes were compared between groups by two-way mixed ANOVA with Greenhouse-Geisser correction. There was a significant difference at 50Hz. SS=sum of squares; MS=mean square; df=degrees of freedom.

| **Tests of Between-Subjects Effects** | | | | | | | |
|---|---|---|---|---|---|---|---|
| **Frequency** | **Source** | **SS** | **df** | **MS** | **F** | **p value** | **Partial Eta Squared** |
| 5Hz *Prdm8$^{-/-}$* vs WT+PTX | Intercept | 93.533 | 1 | 93.533 | 990.120 | <0.001 | 0.980 |
| | group | 0.006 | 1 | 0.006 | 0.058 | 0.812 | 0.003 |
| | Error | 1.889 | 20 | 0.094 | | | |
| 20Hz *Prdm8$^{-/-}$* vs WT+PTX | Intercept | 56.767 | 1 | 56.767 | 1092.191 | <0.001 | 0.986 |
| | genotype | 0.024 | 1 | 0.024 | 0.461 | 0.507 | 0.030 |
| | Error | 0.780 | 15 | 0.052 | | | |
| 50Hz *Prdm8$^{-/-}$* vs WT+PTX | Intercept | 51.586 | 1 | 51.586 | 491.610 | <0.001 | 0.970 |
| | group | 0.962 | 1 | 0.962 | 9.171 | 0.008 | 0.379 |
| | Error | 1.574 | 15 | 0.105 | | | |

**Table S10**

**Within group statistical comparisons of short-term dynamics of medial perforant path-granule cell synapses to 5Hz (n=10|9), 20Hz (n=6|9) and 50Hz (n=6|9) stimulation in wild type (WT) hippocampal slices *in vitro* with and without picrotoxin (PTX).** Normalized peak amplitudes were compared within-groups by two-way mixed ANOVA with the Greenhouse-Geisser correction. There was a significant interaction at 50Hz. SS=sum of squares; MS=mean square; df=degrees of freedom.

| **Tests of Within-Subjects Effects** | | | | | | | |
|---|---|---|---|---|---|---|---|
| **Frequency** | **Source** | **SS** | **df** | **MS** | **F** | **p value** | **Partial Eta Squared** |
| 5Hz WT vs WT+PTX | Stim | 0.795 | 1.944 | 0.409 | 62.341 | <0.001 | 0.786 |
| | stim*group | 0.014 | 1.944 | 0.007 | 1.114 | 0.339 | 0.061 |
| | Error (stim) | 0.217 | 33.041 | 0.007 | | | |
| 20Hz WT vs WT+PTX | Stim | 2.791 | 1.232 | 2.265 | 54.034 | <0.001 | 0.806 |
| | stim*group | 0.012 | 1.232 | 0.009 | 0.224 | 0.693 | 0.017 |
| | Error (stim) | 0.671 | 16.019 | 0.042 | | | |
| 50Hz WT vs WT+PTX | Stim | 6.855 | 1.308 | 5.243 | 26.739 | <0.001 | 0.673 |
| | stim*group | 1.142 | 1.308 | 0.873 | 4.455 | 0.041 | 0.255 |
| | Error (stim) | 3.333 | 16.999 | 0.196 | | | |

**Table S11**

**Between group statistical comparisons of short-term dynamics of medial perforant path-granule cell synapses to 5Hz (n=10|9), 20Hz (n=6|9) and 50Hz (n=6|9) stimulation in WT hippocampal slices (n=9|6) *in vitro* with and without picrotoxin (PTX).** Normalized peak amplitudes were compared between-groups by two-way mixed ANOVA. There were significant differences between genotypes to stimulations at 20Hz and 50Hz. SS=sum of squares; MS=mean square; df=degrees of freedom.

| **Tests of Between-Subjects Effects** | | | | | | | |
|---|---|---|---|---|---|---|---|
| **Frequency** | **Source** | **SS** | **df** | **MS** | **F** | **p value** | **Partial Eta Squared** |
| 5Hz | Intercept | 81.671 | 1 | 81.671 | 876.229 | <0.001 | 0.981 |
| WT vs WT+PTX | group | 0.026 | 1 | 0.026 | 0.276 | 0.606 | 0.016 |
| | Error | 1.585 | 17 | 0.093 | | | |
| 20Hz | Intercept | 39.636 | 1 | 39.636 | 862.266 | <0.001 | 0.985 |
| WT vs WT+PTX | group | 0.629 | 1 | 0.629 | 13.693 | 0.003 | 0.513 |
| | Error | 0.598 | 13 | 0.046 | | | |
| 50Hz | Intercept | 34.658 | 1 | 34.658 | 493.651 | <0.001 | 0.974 |
| WT vs WT+PTX | group | 2.687 | 1 | 2.687 | 38.268 | <0.001 | 0.746 |
| | Error | 0.913 | 13 | 0.070 | | | |

**Table S12**

**Within group statistical comparisons of short-term dynamics of medial perforant path-granule cell synapses to 5Hz, 20Hz and 50Hz stimulation in *Prdm8*$^{-/-}$ and WT hippocampal slices (n=6|9) *in vitro* in the presence of picrotoxin (PTX).** Normalized peak amplitudes were compared within-groups by two-way mixed ANOVA with the Greenhouse-Geisser correction. There were no significant interactions between stimulation frequency and group. SS=sum of squares; MS=mean square; df=degrees of freedom.

| **Tests of Within-Subjects Effects** | | | | | | | |
|---|---|---|---|---|---|---|---|
| **Frequency** | **Source** | **SS** | **df** | **MS** | **F** | **p value** | **Partial Eta Squared** |
| 5Hz | stim | 0.604 | 2.174 | 0.278 | 58.321 | $<0.001$ | 0.818 |
| WT+PTX vs *Prdm8*$^{-/-}$+PTX | stim*group | 0.012 | 2.174 | 0.005 | 1.135 | 0.339 | 0.080 |
| | Error (stim) | 0.135 | 28.262 | 0.005 | | | |
| 20Hz | stim | 3.268 | 1.258 | 2.597 | 47.332 | .000 | 0.785 |
| WT+PTX vs *Prdm8*$^{-/-}$+PTX | stim*group | 0.034 | 1.258 | 0.027 | 0.491 | .536 | 0.036 |
| | Error (stim) | 0.897 | 16.358 | 0.055 | | | |
| 50Hz | stim | 13.908 | 1.301 | 10.687 | 33.998 | $<0.001$ | 0.723 |
| WT+PTX vs *Prdm8*$^{-/-}$+PTX | stim*group | 0.061 | 1.301 | 0.047 | 0.148 | 0.770 | 0.011 |
| | Error (stim) | 5.318 | 16.918 | 0.314 | | | |

**Table S13**

**Between group statistical comparisons of short-term dynamics of medial perforant path-granule cell synapses to 5Hz, 20Hz and 50Hz stimulation in *Prdm8*$^{-/-}$ and WT hippocampal slices (n=6|9) *in vitro* in the presence of picrotoxin (PTX).** Normalized peak amplitudes were compared within-groups by two-way mixed ANOVA. There were no significant differences between genotypes to stimulations at 5Hz, 20Hz and 50Hz. SS=sum of squares; MS=mean square; df=degrees of freedom.

| **Tests of Between-Subjects Effects** | | | | | | | |
|---|---|---|---|---|---|---|---|
| **Frequency** | **Source** | **SS** | **df** | **MS** | **F** | **p value** | **Partial Eta Squared** |
| 5Hz<br>WT+PTX vs *Prdm8*$^{-/-}$+PTX | Intercept | 61.064 | 1 | 61.064 | 842.865 | <0.001 | 0.985 |
| | group | 0.042 | 1 | 0.042 | 0.573 | 0.462 | 0.042 |
| | Error | 0.942 | 13 | 0.072 | | | |
| 20Hz<br>WT+PTX vs *Prdm8*$^{-/-}$+PTX | Intercept | 45.435 | 1 | 45.435 | 630.585 | <0.001 | 0.980 |
| | group | 0.121 | 1 | 0.121 | 1.686 | 0.217 | 0.115 |
| | Error | 0.937 | 13 | 0.072 | | | |
| 50Hz<br>WT+PTX vs *Prdm8*$^{-/-}$+PTX | Intercept | 49.908 | 1 | 49.908 | 362.921 | <0.001 | 0.965 |
| | group | 0.213 | 1 | 0.213 | 1.550 | 0.235 | 0.107 |
| | Error | 1.788 | 13 | 0.138 | | | |

**Table S14**

**Statistical comparisons of normalized peak amplitudes of field excitatory postsynaptic potentials (fEPSP) recorded in granule cells *in vitro* to 5Hz stimulation of the medial perforant path.** Normalized peak amplitudes were compared by one-way ANOVA with Post-hoc Tukey's correction for multiple comparisons. Group comparisons were: (i) *Prdm8*$^{-/-}$ vs wild type (WT) (n=13|10); (ii) *Prdm8*$^{-/-}$ vs WT + PTX (n=13|9); (iii) WT vs WT + PTX (n=10|9); (iv) *Prdm8*$^{-/-}$ + PTX vs WT + PTX (n=6|9). MD=mean difference; SE=standard error.

| | | | | | | 95% Confidence Interval | |
|---|---|---|---|---|---|---|---|
| **Peak** | **Genotype** | **Genotype** | **Mean** | **SE** | **p value** | **Lower** | **Upper** |
| Peak 2 | WT | *Prdm8*$^{-/-}$ | 0.012 | 0.041 | 0.955 | -0.089 | 0.113 |
| | | WT + PTX | 0.026 | 0.045 | 0.835 | -0.084 | 0.136 |
| | | *Prdm8*$^{-/-}$ + PTX | 0.019 | 0.451 | 0.907 | -0.094 | 0.132 |
| | *Prdm8*$^{-/-}$ | WT | -0.012 | 0.041 | 0.955 | -0.113 | 0.089 |
| | | WT + PTX | 0.014 | 0.042 | 0.943 | -0.090 | 0.118 |
| | | *Prdm8*$^{-/-}$ + PTX | 0.002 | 0.025 | 0.998 | -0.060 | 0.063 |
| | WT + PTX | WT | -0.026 | 0.045 | 0.835 | -0.136 | 0.084 |
| | | *Prdm8*$^{-/-}$ | -0.014 | 0.042 | 0.943 | -0.118 | 0.090 |
| | | *Prdm8*$^{-/-}$ +PTX | -0.007 | 0.046 | 0.989 | -0.122 | 0.109 |
| | *Prdm8*$^{-/-}$ +PTX | WT | -0.191 | 0.451 | 0.907 | -0.132 | 0.094 |
| | | *Prdm8*$^{-/-}$ | -0.002 | 0.024 | 0.998 | -0.063 | 0.060 |
| | | WT + PTX | 0.007 | 0.046 | 0.989 | -1.092 | 0.122 |
| Peak 3 | WT | *Prdm8*$^{-/-}$ | -0.004 | 0.046 | 0.995 | -0.117 | 0.108 |
| | | WT + PTX | -0.024 | 0.050 | 0.878 | -0.147 | 0.099 |
| | | *Prdm8*$^{-/-}$ + PTX | 0.017 | 0.049 | 0.936 | -0.105 | 0.133 |
| | *Prdm8*$^{-/-}$ | WT | 0.004 | 0.046 | 0.995 | -0.108 | 0.117 |
| | | WT + PTX | -0.020 | 0.047 | 0.908 | -0.136 | 0.096 |
| | | *Prdm8*$^{-/-}$ + PTX | 0.008 | 0.031 | 0.968 | -0.070 | 0.086 |
| | | WT | 0.024 | 0.050 | 0.878 | -0.100 | 0.147 |

| | | | | | | 95% Confidence Interval | |
|---|---|---|---|---|---|---|---|
| **Peak** | **Genotype** | **Genotype** | **Mean** | **SE** | **p value** | **Lower** | **Upper** |
| | WT + PTX | *Prdm8*$^{-/-}$ | 0.020 | 0.047 | 0.908 | -0.010 | 0.136 |
| | | *Prdm8*$^{-/-}$ +PTX | 0.041 | 0.050 | 0.691 | -0.084 | 0.166 |
| | *Prdm8*$^{-/-}$ +PTX | WT | -0.168 | 0.486 | 0.936 | -1.387 | 0.084 |
| | | *Prdm8*$^{-/-}$ | -0.008 | 0.031 | 0.968 | -0.086 | 0.070 |
| | | WT + PTX | 0.04§ | 0.050 | 0.691 | -0.166 | 0.084 |
| Peak 4 | WT | *Prdm8*$^{-/-}$ | -0.023 | 0.046 | 0.870 | -0.136 | 0.090 |
| | | WT + PTX | -0.041 | 0.050 | 0.687 | -0.164 | 0.082 |
| | | *Prdm8*$^{-/-}$ + PTX | 0.002 | 0.051 | 0.999 | -0.126 | 0.130 |
| | *Prdm8*$^{-/-}$ | WT | 0.023 | 0.046 | 0.870 | -0.090 | 0.136 |
| | | WT + PTX | -0.018 | 0.047 | 0.919 | -0.135 | 0.098 |
| | | *Prdm8*$^{-/-}$ + PTX | 0.011 | 0.034 | 0.943 | -0.072 | 0.094 |
| | WT + PTX | WT | 0.041 | 0.050 | 0.687 | -0.082 | 0.164 |
| | | *Prdm8*$^{-/-}$ | 0.018 | 0.047 | 0.919 | -0.098 | 0.135 |
| | | *Prdm8*$^{-/-}$ +PTX | 0.043 | 0.051 | 0.999 | -0.126 | 0.130 |
| | *Prdm8*$^{-/-}$ +PTX | WT | -0.002 | 0.051 | 0.999 | -0.130 | 0.126 |
| | | *Prdm8*$^{-/-}$ | -0.011 | 0.034 | 0.943 | -0.094 | 0.072 |
| | | WT + PTX | 0.043 | 0.052 | 0.686 | -0.174 | 0.087 |
| Peak 5 | WT | *Prdm8*$^{-/-}$ | -0.012 | 0.045 | 0.962 | -0.124 | 0.100 |
| | | WT + PTX | -0.028 | 0.049 | 0.841 | -0.150 | 0.094 |
| | | *Prdm8*$^{-/-}$ + PTX | 0.007 | 0.511 | 0.989 | -0.121 | 0.136 |
| | *Prdm8*$^{-/-}$ | WT | 0.012 | 0.045 | 0.962 | -0.010 | 0.124 |
| | | WT + PTX | -0.016 | 0.047 | 0.938 | -0.131 | 0.099 |
| | | *Prdm8*$^{-/-}$ + PTX | 0.008 | 0.035 | 0.975 | -0.080 | 0.095 |
| | WT + PTX | WT | 0.028 | 0.049 | 0.841 | -0.094 | 0.150 |
| | | *Prdm8*$^{-/-}$ | 0.016 | 0.047 | 0.938 | -0.099 | 0.131 |
| | | *Prdm8*$^{-/-}$ +PTX | 0.035 | 0.521 | 0.782 | -0.096 | 0.166 |
| | *Prdm8*$^{-/-}$ +PTX | WT | -0.007 | 0.511 | 0.989 | -0.136 | 0.121 |
| | | *Prdm8*$^{-/-}$ | -0.008 | 0.035 | 0.975 | -0.095 | 0.080 |
| | | WT + PTX | -0.351 | 0.522 | 0.782 | -0.166 | 0.096 |

| | | | | | | 95% Confidence Interval | |
|---|---|---|---|---|---|---|---|
| **Peak** | **Genotype** | **Genotype** | **Mean** | **SE** | **p value** | **Lower** | **Upper** |
| Peak 6 | WT | *Prdm8*$^{-/-}$ | -0.030 | 0.046 | 0.800 | -0.144 | 0.085 |
| | | WT + PTX | -0.032 | 0.051 | 0.805 | -0.157 | 0.093 |
| | | *Prdm8*$^{-/-}$ + PTX | -0.185 | 0.052 | 0.933 | -0.150 | 0.113 |
| | *Prdm8*$^{-/-}$ | WT | 0.030 | 0.046 | 0.800 | -0.085 | 0.144 |
| | | WT + PTX | -0.002 | 0.048 | 0.999 | -0.120 | 0.116 |
| | | *Prdm8*$^{-/-}$ + PTX | 0.002 | 0.037 | 0.998 | -0.089 | 0.094 |
| | WT + PTX | WT | 0.032 | 0.051 | 0.805 | -0.093 | 0.157 |
| | | *Prdm8*$^{-/-}$ | 0.002 | 0.048 | 0.999 | -0.116 | 0.120 |
| | | *Prdm8*$^{-/-}$ +PTX | 0.013 | 0.053 | 0.966 | 0.147 | 0.121 |
| | *Prdm8*$^{-/-}$ +PTX | WT | 0.019 | 0.052 | 0.933 | -0.113 | 0.150 |
| | | *Prdm8*$^{-/-}$ | -0.002 | 0.037 | 0.998 | -0.094 | 0.089 |
| | | WT + PTX | -0.013 | 0.053 | 0.966 | -0.147 | 0.121 |
| Peak 7 | WT | *Prdm8*$^{-/-}$ | -0.023 | 0.049 | 0.883 | -0.144 | 0.098 |
| | | WT + PTX | -0.027 | 0.054 | 0.866 | -0.160 | 0.105 |
| | | *Prdm8*$^{-/-}$ + PTX | 0.011 | 0.058 | 0.980 | -0.134 | 0.156 |
| | *Prdm8*$^{-/-}$ | WT | 0.023 | 0.049 | 0.883 | -0.098 | 0.144 |
| | | WT + PTX | -0.004 | 0.050 | 0.996 | -0.129 | 0.121 |
| | | *Prdm8*$^{-/-}$ + PTX | 0.019 | 0.040 | 0.879 | -0.080 | 0.118 |
| | WT + PTX | WT | 0.027 | 0.054 | 0.866 | -0.105 | 0.160 |
| | | *Prdm8*$^{-/-}$ | 0.004 | 0.050 | 0.996 | -0.121 | 0.129 |
| | | *Prdm8*$^{-/-}$ +PTX | 0.039 | 0.059 | 0.793 | -0.110 | 0.187 |
| | *Prdm8*$^{-/-}$ +PTX | WT | -0.011 | 0.058 | 0.980 | -0.157 | 0.134 |
| | | *Prdm8*$^{-/-}$ | -0.020 | 0.040 | 0.879 | -0.118 | 0.080 |
| | | WT + PTX | -0.039 | 0.059 | 0.793 | -0.187 | 0.110 |
| Peak 8 | WT | *Prdm8*$^{-/-}$ | -0.010 | 0.045 | 0.973 | -0.121 | 0.101 |
| | | WT + PTX | -0.037 | 0.049 | 0.738 | -0.157 | 0.084 |
| | | *Prdm8*$^{-/-}$ + PTX | 0.014 | 0.052 | 0.963 | -0.118 | 0.145 |
| | *Prdm8*$^{-/-}$ | WT | 0.010 | 0.045 | 0.973 | -0.101 | 0.121 |
| | | WT + PTX | -0.027 | 0.046 | 0.834 | -0.141 | 0.087 |

| | | | | | | 95% Confidence Interval | |
|---|---|---|---|---|---|---|---|
| **Peak** | **Genotype** | **Genotype** | **Mean** | **SE** | **p value** | **Lower** | **Upper** |
| | | *Prdm8*$^{-/-}$ + PTX | 0.012 | 0.040 | 0.948 | -0.087 | 0.111 |
| | WT + PTX | WT | 0.037 | 0.049 | 0.738 | -0.084 | 0.157 |
| | | *Prdm8*$^{-/-}$ | 0.027 | 0.046 | 0.834 | -0.087 | 0.141 |
| | | *Prdm8*$^{-/-}$ +PTX | 0.050 | 0.053 | 0.622 | -0.084 | 0.184 |
| | *Prdm8*$^{-/-}$ +PTX | WT | -0.014 | 0.052 | 0.963 | -0.145 | 0.118 |
| | | *Prdm8*$^{-/-}$ | -0.012 | 0.040 | 0.948 | -0.111 | 0.087 |
| | | WT + PTX | -0.050 | 0.053 | 0.622 | -0.184 | 0.084 |
| Peak 9 | WT | *Prdm8*$^{-/-}$ | -0.019 | 0.049 | 0.925 | -0.140 | 0.103 |
| | | WT + PTX | -0.032 | 0.054 | 0.828 | -0.165 | 0.101 |
| | | *Prdm8*$^{-/-}$ + PTX | 0.015 | 0.061 | 0.967 | -0.138 | 0.168 |
| | *Prdm8*$^{-/-}$ | WT | 0.019 | 0.049 | 0.925 | -0.103 | 0.140 |
| | | WT + PTX | -0.013 | 0.051 | 0.964 | -0.139 | 0.113 |
| | | *Prdm8*$^{-/-}$ + PTX | -0.012 | 0.040 | 0.948 | -0.111 | 0.087 |
| | WT + PTX | WT | 0.032 | 0.054 | 0.828 | -0.101 | 0.165 |
| | | *Prdm8*$^{-/-}$ | 0.013 | 0.051 | 0.964 | -0.113 | 0.139 |
| | | *Prdm8*$^{-/-}$ +PTX | 0.047 | 0.062 | 0.736 | -0.109 | 0.203 |
| | *Prdm8*$^{-/-}$ +PTX | WT | -0.050 | 0.061 | 0.967 | -0.168 | 0.138 |
| | | *Prdm8*$^{-/-}$ | 0.022 | 0.043 | 0.872 | -0.086 | 0.129 |
| | | WT + PTX | -0.047 | 0.062 | 0.736 | -0.203 | 0.109 |
| Peak 10 | WT | *Prdm8*$^{-/-}$ | -0.016 | 0.047 | 0.934 | -0.132 | 0.099 |
| | | WT + PTX | -0.032 | 0.051 | 0.802 | -0.158 | 0.094 |
| | | *Prdm8*$^{-/-}$ + PTX | 0.028 | 0.058 | 0.881 | -0.118 | 0.175 |
| | *Prdm8*$^{-/-}$ | WT | 0.016 | 0.047 | 0.934 | -0.099 | 0.131 |
| | | WT + PTX | -0.016 | 0.048 | 0.941 | -0.135 | 0.103 |
| | | *Prdm8*$^{-/-}$ + PTX | 0.032 | 0.041 | 0.726 | -0.071 | 0.134 |
| | WT + PTX | WT | 0.032 | 0.051 | 0.802 | -0.094 | 0.158 |
| | | *Prdm8*$^{-/-}$ | 0.016 | 0.048 | 0.941 | -0.103 | 0.135 |
| | | *Prdm8*$^{-/-}$ +PTX | 0.060 | 0.060 | 0.575 | -0.890 | 0.210 |
| | | WT | -0.028 | 0.058 | 0.881 | -0.175 | 0.118 |

| | | | | | | 95% Confidence Interval | |
|---|---|---|---|---|---|---|---|
| **Peak** | **Genotype** | **Genotype** | **Mean** | **SE** | **p value** | **Lower** | **Upper** |
| | *Prdm8*$^{-/-}$ +PTX | *Prdm8*$^{-/-}$ | -0.032 | 0.041 | 0.726 | -0.134 | 0.071 |
| | | WT + PTX | -0.060 | 0.060 | 0.575 | -0.210 | 0.089 |

**Table S15**

**Statistical comparisons of normalized peak amplitudes of field excitatory postsynaptic potentials (fEPSP) recorded in granule cells *in vitro* to 20Hz stimulation of the medial perforant path.** Normalized peak amplitudes were compared by one-way ANOVA with Post-hoc Tukey's correction for multiple comparisons. Group comparisons were: (i) *Prdm8*$^{-/-}$ vs WT (n=8|6); (ii) *Prdm8*$^{-/-}$ vs WT + PTX (n=8|9); (iii) WT vs WT + PTX (n=6|9); (iv) *Prdm8*$^{-/-}$ + PTX vs WT + PTX (n=6|9). MD=mean difference; SE=standard error.

| | | | | | | **95% Confidence Interval** | |
|---|---|---|---|---|---|---|---|
| **Peak** | **Genotype** | **Genotype** | **Mean** | **SE** | **p value** | **Lower** | **Upper** |
| Peak 2 | WT | *Prdm8*$^{-/-}$ | -0.042 | 0.035 | 0.453 | -0.130 | 0.045 |
| | | WT + PTX | -0.043 | 0.034 | 0.425 | -0.129 | 0.043 |
| | | *Prdm8*$^{-/-}$ + PTX | 0.123 | 0.114 | 0.536 | -0.167 | 0.414 |
| | *Prdm8*$^{-/-}$ | WT | 0.042 | 0.035 | 0.453 | -0.045 | 0.130 |
| | | WT + PTX | -0.001 | 0.031 | 1.000 | -0.080 | 0.078 |
| | | *Prdm8*$^{-/-}$ + PTX | -0.010 | 0.029 | 0.941 | -0.085 | 0.065 |
| | WT + PTX | WT | 0.043 | 0.034 | 0.425 | -0.043 | 0.129 |
| | | *Prdm8*$^{-/-}$ | 0.001 | 0.031 | 1.000 | -0.078 | 0.080 |
| | | *Prdm8*$^{-/-}$ +PTX | -0.014 | 0.104 | 0.989 | -0.280 | 0.251 |
| | *Prdm8*$^{-/-}$ +PTX | WT | 0.123 | 0.114 | 0.536 | -0.167 | 0.414 |
| | | *Prdm8*$^{-/-}$ | 0.010 | 0.029 | 0.941 | -0.065 | 0.085 |
| | | WT + PTX | 0.014 | 0.104 | 0.989 | -0.251 | 0.280 |
| Peak 3 | WT | *Prdm8*$^{-/-}$ | -0.063 | 0.031 | 0.129 | -0.141 | 0.015 |
| | | WT + PTX | -0.096 | 0.030 | 0.012* | -0.173 | -0.020 |
| | | *Prdm8*$^{-/-}$ + PTX | -0.130 | 0.066 | 0.150* | -0.300 | 0.039 |
| | *Prdm8*$^{-/-}$ | WT | 0.063 | 0.031 | 0.129 | -0.015 | 0.141 |
| | | WT + PTX | -0.033 | 0.028 | 0.468 | -0.104 | 0.037 |
| | | *Prdm8*$^{-/-}$ + PTX | -0.014 | 0.035 | 0.913 | -0.105 | 0.076 |
| | | WT | 0.096 | 0.030 | 0.012* | 0.020 | 0.173 |

| | | | | | | **95% Confidence Interval** | |
|---|---|---|---|---|---|---|---|
| **Peak** | **Genotype** | **Genotype** | **Mean** | **SE** | **p value** | **Lower** | **Upper** |
| | WT + PTX | *Prdm8*$^{-/-}$ | 0.033 | 0.028 | 0.468 | -0.037 | 0.104 |
| | | *Prdm8*$^{-/-}$ +PTX | 0.026 | 0.060 | 0.902 | -0.128 | 0.180 |
| | *Prdm8*$^{-/-}$ +PTX | WT | 0.130 | 0.066 | 0.150* | -0.039 | 0.299 |
| | | *Prdm8*$^{-/-}$ | 0.014 | 0.035 | 0.913 | -0.076 | 0.105 |
| | | WT + PTX | -0.026 | 0.060 | 0.902 | -0.180 | 0.128 |
| Peak 4 | WT | *Prdm8-/-* | -0.078 | 0.034 | 0.074 | -0.164 | 0.007 |
| | | WT + PTX | -0.116 | 0.033 | 0.005** | -0.199 | -0.033 |
| | | *Prdm8*$^{-/-}$ + PTX | -0.071 | 0.046 | 0.304 | -0.189 | 0.048 |
| | *Prdm8*$^{-/-}$ | WT | 0.078 | 0.034 | 0.074 | -0.007 | 0.164 |
| | | WT + PTX | -0.038 | 0.030 | 0.438 | -0.114 | 0.039 |
| | | *Prdm8*$^{-/-}$ + PTX | 0.025 | 0.037 | 0.778 | -0.069 | 0.119 |
| | WT + PTX | WT | 0.116 | 0.033 | 0.005** | 0.033 | 0.199 |
| | | *Prdm8*$^{-/-}$ | 0.038 | 0.030 | 0.438 | -0.039 | 0.114 |
| | | *Prdm8*$^{-/-}$ +PTX | 0.085 | 0.042 | 0.137 | -0.023 | 0.193 |
| | *Prdm8*$^{-/-}$ +PTX | WT | 0.071 | 0.046 | 0.304 | -0.048 | 0.189 |
| | | *Prdm8*$_{-/-}$ | -0.025 | 0.037 | 0.778 | -0.119 | 0.069 |
| | | WT + PTX | -0.085 | 0.042 | 0.137 | -0.193 | 0.023 |
| Peak 5 | WT | *Prdm8*$^{-/-}$ | -0.118 | 0.036 | 0.010* | -0.210 | -0.027 |
| | | WT + PTX | -0.135 | 0.035 | 0.003** | -0.225 | -0.046 |
| | | *Prdm8-/-* + PTX | -0.083 | 0.055 | 0.306 | -0.223 | 0.057 |
| | *Prdm8*$^{-/-}$ | WT | 0.118 | 0.036 | 0.010* | 0.027 | 0.210 |
| | | WT + PTX | -0.017 | 0.033 | 0.866 | -0.099 | 0.066 |
| | | *Prdm8*$^{-/-}$ + PTX | 0.048 | 0.043 | 0.508 | -0.061 | 0.157 |
| | WT + PTX | WT | 0.135 | 0.035 | 0.003** | 0.046 | 0.225 |
| | | *Prdm8*$^{-/-}$ | 0.017 | 0.033 | 0.866 | -0.066 | 0.099 |
| | | *Prdm8*$^{-/-}$ +PTX | 0.076 | 0.050 | 0.303 | -0.051 | 0.204 |
| | *Prdm8*$^{-/-}$ +PTX | WT | 0.083 | 0.055 | 0.306 | -0.057 | 0.223 |
| | | *Prdm8*$^{-/-}$ | -0.048 | 0.043 | 0.508 | -0.157 | 0.061 |
| | | WT + PTX | -0.076 | 0.050 | 0.303 | -0.204 | 0.051 |

| | | | | | | 95% Confidence Interval | |
|---|---|---|---|---|---|---|---|
| **Peak** | **Genotype** | **Genotype** | **Mean** | **SE** | **p value** | **Lower** | **Upper** |
| Peak 6 | WT | *Prdm8*$^{-/-}$ | -0.103 | 0.035 | 0.019* | -0.191 | -0.016 |
| | | WT + PTX | -0.133 | 0.034 | 0.002** | -0.218 | -0.047 |
| | | *Prdm8*$^{-/-}$ + PTX | -0.071 | 0.050 | 0.351 | -0.198 | 0.056 |
| | *Prdm8*$^{-/-}$ | WT | 0.103 | 0.035 | 0.019* | 0.016 | 0.191 |
| | | WT + PTX | -0.029 | 0.031 | 0.617 | -0.108 | 0.049 |
| | | *Prdm8*$^{-/-}$ + PTX | 0.040 | 0.037 | 0.536 | -0.055 | 0.135 |
| | WT + PTX | WT | 0.133 | 0.034 | 0.002** | 0.047 | 0.218 |
| | | *Prdm8*$^{-/-}$ | 0.029 | 0.031 | 0.617 | -0.049 | 0.108 |
| | | *Prdm8*$^{-/-}$ +PTX | 0.081 | 0.045 | 0.202 | -0.035 | 0.197 |
| | *Prdm8*$^{-/-}$ +PTX | WT | 0.071 | 0.050 | 0.351 | -0.056 | 0.198 |
| | | *Prdm8*$^{-/-}$ | -0.040 | 0.037 | 0.536 | -0.135 | 0.055 |
| | | WT + PTX | -0.081 | 0.045 | 0.202 | -0.197 | 0.035 |
| Peak 7 | WT | *Prdm8*$^{-/-}$ | -0.119 | 0.041 | 0.023* | -0.222 | -0.015 |
| | | WT + PTX | -0.102 | 0.040 | 0.047* | -0.203 | -0.001 |
| | | *Prdm8*$^{-/-}$ + PTX | -0.052 | 0.061 | 0.674 | -0.207 | 0.103 |
| | *Prdm8*$^{-/-}$ | WT | 0.119 | 0.041 | 0.023* | 0.015 | 0.222 |
| | | WT + PTX | 0.017 | 0.037 | 0.895 | -0.077 | 0.110 |
| | | *Prdm8*$^{-/-}$ + PTX | 0.073 | 0.045 | 0.258 | -0.042 | 0.189 |
| | WT + PTX | WT | 0.102 | 0.040 | 0.047* | 0.001 | 0.203 |
| | | *Prdm8*$^{-/-}$ | -0.017 | 0.037 | 0.895 | -0.110 | 0.076 |
| | | *Prdm8*$^{-/-}$ +PTX | 0.058 | 0.055 | 0.554 | -0.083 | 0.200 |
| | *Prdm8*$^{-/-}$ +PTX | WT | 0.052 | 0.061 | 0.674 | -0.103 | 0.207 |
| | | *Prdm8*$^{-/-}$ | -0.073 | 0.045 | 0.258 | -0.189 | 0.042 |
| | | WT + PTX | -0.058 | 0.055 | 0.554 | -0.200 | 0.083 |
| Peak 8 | WT | *Prdm8*$^{-/-}$ | -0.125 | 0.039 | 0.011* | -0.223 | -0.027 |
| | | WT + PTX | -0.139 | 0.038 | 0.004** | -0.235 | -0.043 |
| | | *Prdm8*$^{-/-}$ + PTX | -0.061 | 0.050 | 0.463 | -0.189 | 0.068 |
| | *Prdm8*$^{-/-}$ | WT | 0.125 | 0.039 | 0.011* | 0.027 | 0.223 |
| | | WT + PTX | -0.014 | 0.035 | 0.918 | -0.102 | 0.075 |

| | | | | | | 95% Confidence Interval | |
|---|---|---|---|---|---|---|---|
| **Peak** | **Genotype** | **Genotype** | **Mean** | **SE** | **p value** | **Lower** | **Upper** |
| | | *Prdm8*$^{-/-}$ + PTX | 0.063 | 0.040 | 0.291 | -0.041 | 0.167 |
| | WT + PTX | WT | 0.139 | 0.038 | 0.004** | 0.043 | 0.235 |
| | | *Prdm8*$^{-/-}$ | 0.014 | 0.035 | 0.918 | -0.075 | 0.102 |
| | | *Prdm8*$^{-/-}$ +PTX | 0.085 | 0.046 | 0.181 | -0.032 | 0.202 |
| | *Prdm8*$^{-/-}$ +PTX | WT | 0.061 | 0.050 | 0.463 | -0.068 | 0.189 |
| | | *Prdm8*$^{-/-}$ | -0.063 | 0.040 | 0.291 | -0.167 | 0.041 |
| | | WT + PTX | -0.085 | 0.046 | 0.181 | -0.202 | 0.032 |
| Peak 9 | WT | *Prdm8*$^{-/-}$ | -0.121 | 0.041 | 0.020* | -0.224 | -0.018 |
| | | WT + PTX | -0.120 | 0.040 | 0.018* | -0.220 | -0.019 |
| | | *Prdm8*$^{-/-}$ + PTX | -0.054 | 0.059 | 0.636 | -0.204 | 0.096 |
| | *Prdm8*$^{-/-}$ | WT | 0.121 | 0.041 | 0.020* | 0.018 | 0.224 |
| | | WT + PTX | 0.001 | 0.037 | 0.999 | -0.091 | 0.094 |
| | | *Prdm8*$^{-/-}$ + PTX | 0.07 | 0.049 | 0.303 | -0.051 | 0.199 |
| | WT + PTX | WT | 0.120 | 0.040 | 0.018* | 0.019 | 0.220 |
| | | *Prdm8*$^{-/-}$ | -0.001 | 0.037 | 0.999 | -0.094 | 0.091 |
| | | *Prdm8*$^{-/-}$ +PTX | 0.069 | 0.054 | 0.421 | -0.068 | 0.206 |
| | *Prdm8*$^{-/-}$ +PTX | WT | 0.054 | 0.059 | 0.636 | -0.096 | 0.204 |
| | | *Prdm8*$^{-/-}$ | -0.074 | 0.049 | 0.303 | -0.199 | 0.051 |
| | | WT + PTX | -0.069 | 0.054 | 0.421 | -0.206 | 0.068 |
| Peak 10 | WT | *Prdm8*$^{-/-}$ | -0.128 | 0.042 | 0.015* | -0.233 | -0.023 |
| | | WT + PTX | -0.148 | 0.041 | 0.004** | -0.251 | -0.046 |
| | | *Prdm8*$^{-/-}$ + PTX | - | 0.052 | 0.452 | -0.195 | 0.068 |
| | *Prdm8*$^{-/-}$ | WT | 0.128 | 0.042 | 0.015* | 0.023 | 0.233 |
| | | WT + PTX | -0.020 | 0.037 | 0.855 | -0.115 | 0.075 |
| | | *Prdm8*$^{-/-}$ + PTX | 0.062 | 0.045 | 0.370 | -0.050 | 0.178 |
| | WT + PTX | WT | 0.148 | 0.041 | 0.004** | 0.046 | 0.251 |
| | | *Prdm8*$^{-/-}$ | 0.020 | 0.037 | 0.855 | -0.075 | 0.115 |
| | | *Prdm8*$^{-/-}$ +PTX | 0.085 | 0.047 | 0.195 | -0.035 | 0.205 |
| | | WT | 0.063 | 0.052 | 0.452 | -0.068 | 0.195 |

| | | | | | | 95% Confidence Interval | |
|---|---|---|---|---|---|---|---|
| **Peak** | **Genotype** | **Genotype** | **Mean** | **SE** | **p value** | **Lower** | **Upper** |
| | *Prdm8*$^{-/-}$ +PTX | *Prdm8*$^{-/-}$ | -0.062 | 0.045 | 0.370 | -0.178 | 0.053 |
| | | WT + PTX | -0.085 | 0.047 | 0.195 | -0.205 | 0.035 |

**Table S16**

**Statistical comparisons of normalized peak amplitudes of field excitatory postsynaptic potentials (fEPSP) recorded in granule cells *in vitro* to 50Hz stimulation of the medial perforant path.** Normalized peak amplitudes were compared by one-way ANOVA with Post-hoc Tukey's correction for multiple comparisons. Group comparisons were: (i) *Prdm8*$^{-/-}$ vs WT (n=8|6); (ii) *Prdm8*$^{-/-}$ vs WT + PTX (n=8|9); (iii) WT vs WT + PTX (n=6|9); (iv) *Prdm8*$^{-/-}$ + PTX vs WT + PTX (n=6|9). MD=mean difference; SE=standard error.

| | | | | | | **95% Confidence Interval** | |
|---|---|---|---|---|---|---|---|
| **Peak** | **Genotype** | **Genotype** | **Mean** | **SE** | **p value** | **Lower** | **Upper** |
| Peak 2 | WT | *Prdm8*$^{-/-}$ | -0.023 | 0.045 | 0.863 | -0.136 | 0.090 |
| | | WT + PTX | -0.135 | 0.043 | 0.015* | -0.245 | -0.025 |
| | | *Prdm8*$^{-/-}$ + PTX | -0.322 | 0.140 | 0.081 | -0.680 | 0.035 |
| | *Prdm8*$^{-/-}$ | WT | 0.023 | 0.045 | 0.863 | -0.090 | 0.136 |
| | | WT + PTX | -0.112 | 0.040 | 0.029* | -0.213 | -0.010 |
| | | *Prdm8*$^{-/-}$ + PTX | -0.109 | 0.053 | 0.127 | -0.245 | 0.027 |
| | WT + PTX | WT | 0.135 | 0.043 | 0.015* | 0.025 | 0.245 |
| | | *Prdm8*$^{-/-}$ | 0.112 | 0.040 | 0.029* | 0.010 | 0.213 |
| | | *Prdm8*$^{-/-}$ +PTX | -0.016 | 0.128 | 0.992 | -0.342 | 0.311 |
| | *Prdm8*$^{-/-}$ +PTX | WT | 0.322 | 0.140 | 0.081 | -0.035 | 0.680 |
| | | *Prdm8*$^{-/-}$ | 0.109 | 0.053 | 0.127 | -0.027 | 0.245 |
| | | WT + PTX | 0.016 | 0.128 | 0.992 | -0.311 | 0.342 |
| Peak 3 | WT | *Prdm8*$^{-/-}$ | -0.039 | 0.067 | 0.830 | -0.207 | 0.130 |
| | | WT + PTX | -0.353 | 0.065 | 0.001** | -0.517 | -0.188 |
| | | *Prdm8*$^{-/-}$ + PTX | -0.714 | 0.283 | 0.053 | -1.437 | 0.009 |
| | *Prdm8*$^{-/-}$ | WT | 0.039 | 0.067 | 0.830 | -0.130 | 0.207 |
| | | WT + PTX | -0.314 | 0.060 | 0.001** | -0.465 | -0.162 |
| | | *Prdm8*$^{-/-}$ + PTX | -0.299 | 0.064 | 0.001** | -0.462 | -0.136 |
| | | WT | 0.353 | 0.065 | 0.001** | 0.188 | 0.517 |

| | | | | | | 95% Confidence Interval | |
|---|---|---|---|---|---|---|---|
| **Peak** | **Genotype** | **Genotype** | **Mean** | **SE** | **p value** | **Lower** | **Upper** |
| | WT + PTX | *Prdm8*$^{-/-}$ | 0.314 | 0.060 | 0.001** | 0.162 | 0.465 |
| | | *Prdm8*$^{-/-}$ +PTX | 0.031 | 0.259 | 0.992 | -0.629 | 0.690 |
| | *Prdm8*$^{-/-}$ +PTX | WT | 0.714 | 0.283 | 0.053 | -0.009 | 1.437 |
| | | *Prdm8*$^{-/-}$ | 0.299 | 0.064 | 0.001** | 0.136 | 0.462 |
| | | WT + PTX | -0.031 | 0.259 | 0.992 | -0.690 | 0.629 |
| Peak 4 | WT | *Prdm8-/-* | -0.092 | 0.055 | 0.244 | -0.233 | 0.048 |
| | | WT + PTX | -0.312 | 0.054 | 0.001** | -0.449 | -0.176 |
| | | *Prdm8*$^{-/-}$ + PTX | -0.360 | 0.122 | 0.023* | -0.672 | -0.048 |
| | *Prdm8*$^{-/-}$ | WT | 0.092 | 0.055 | 0.244 | -0.048 | 0.233 |
| | | WT + PTX | -0.220 | 0.050 | 0.001** | -0.347 | -0.094 |
| | | *Prdm8*$^{-/-}$ + PTX | -0.168 | 0.055 | 0.018* | -0.308 | -0.027 |
| | WT + PTX | WT | 0.312 | 0.054 | 0.001** | 0.176 | 0.449 |
| | | *Prdm8*$^{-/-}$ | 0.220 | 0.050 | 0.001** | 0.094 | 0.347 |
| | | *Prdm8*$^{-/-}$ +PTX | 0.092 | 0.112 | 0.691 | -0.192 | 0.377 |
| | *Prdm8*$^{-/-}$ +PTX | WT | 0.360 | 0.122 | 0.023* | 0.048 | 0.672 |
| | | *Prdm8*$_{-/-}$ | 0.168 | 0.055 | 0.018* | 0.027 | 0.308 |
| | | WT + PTX | -0.092 | 0.112 | 0.691 | -0.377 | 0.192 |
| Peak 5 | WT | *Prdm8*$^{-/-}$ | -0.137 | 0.049 | 0.029* | -0.260 | -0.013 |
| | | WT + PTX | -0.253 | 0.048 | 0.001** | -0.374 | -0.132 |
| | | *Prdm8-/-* + PTX | -0.163 | 0.066 | 0.058 | -0.331 | 0.005 |
| | *Prdm8*$^{-/-}$ | WT | 0.137 | 0.049 | 0.029* | 0.0131 | 0.260 |
| | | WT + PTX | -0.116 | 0.044 | 0.039* | -0.228 | -0.005 |
| | | *Prdm8*$^{-/-}$ + PTX | -0.025 | 0.053 | 0.885 | -0.162 | 0.111 |
| | WT + PTX | WT | 0.253* | 0.048 | 0.001** | 0.132 | 0.374 |
| | | *Prdm8*$^{-/-}$ | 0.116 | 0.044 | 0.039* | 0.005 | 0.228 |
| | | *Prdm8*$^{-/-}$ +PTX | 0.109 | 0.060 | 0.193 | -0.044 | 0.262 |
| | *Prdm8*$^{-/-}$ +PTX | WT | 0.163 | 0.066 | 0.058 | -0.005 | 0.331 |
| | | *Prdm8*$^{-/-}$ | 0.025 | 0.053 | 0.885 | -0.111 | 0.162 |
| | | WT + PTX | -0.109 | 0.060 | 0.193 | -0.262 | 0.044 |

| | | | | | | 95% Confidence Interval | |
|---|---|---|---|---|---|---|---|
| **Peak** | **Genotype** | **Genotype** | **Mean** | **SE** | **p value** | **Lower** | **Upper** |
| Peak 6 | WT | *Prdm8*$^{-/-}$ | -0.205 | 0.066 | 0.014* | -0.372 | -0.039 |
| | | WT + PTX | -0.260 | 0.064 | 0.002** | -0.422 | -0.097 |
| | | *Prdm8*$^{-/-}$ + PTX | -0.126 | 0.058 | 0.103 | -0.275 | 0.022 |
| | *Prdm8*$^{-/-}$ | WT | 0.205 | 0.066 | 0.014* | 0.039 | 0.372 |
| | | WT + PTX | -0.054 | 0.059 | 0.636 | -0.204 | 0.095 |
| | | *Prdm8*$^{-/-}$ + PTX | 0.030 | 0.071 | 0.905 | -0.152 | 0.212 |
| | WT + PTX | WT | 0.260 | 0.064 | 0.002** | 0.097 | 0.422 |
| | | *Prdm8*$^{-/-}$ | 0.054 | 0.059 | 0.636 | -0.095 | 0.204 |
| | | *Prdm8*$^{-/-}$ +PTX | 0.083 | 0.053 | 0.287 | -0.053 | 0.219 |
| | *Prdm8*$^{-/-}$ +PTX | WT | 0.126 | 0.058 | 0.103 | -0.022 | 0.275 |
| | | *Prdm8*$^{-/-}$ | -0.030 | 0.071 | 0.905 | -0.212 | 0.152 |
| | | WT + PTX | -0.083 | 0.053 | 0.287 | -0.219 | 0.053 |
| Peak 7 | WT | *Prdm8*$^{-/-}$ | -0.183 | 0.057 | 0.013* | -0.328 | -0.037 |
| | | WT + PTX | -0.187 | 0.056 | 0.009** | -0.329 | -0.045 |
| | | *Prdm8*$^{-/-}$ + PTX | -0.078 | 0.057 | 0.383 | -0.224 | 0.068 |
| | *Prdm8*$^{-/-}$ | WT | 0.183 | 0.057 | 0.013* | 0.037 | 0.328 |
| | | WT + PTX | -0.005 | 0.052 | 0.996 | -0.135 | 0.126 |
| | | *Prdm8*$^{-/-}$ + PTX | 0.081 | 0.055 | 0.334 | -0.061 | 0.222 |
| | WT + PTX | WT | 0.187 | 0.056 | 0.009** | 0.045 | 0.329 |
| | | *Prdm8*$^{-/-}$ | 0.004 | 0.052 | 0.996 | -0.126 | 0.135 |
| | | *Prdm8*$^{-/-}$ +PTX | 0.081 | 0.052 | 0.295 | -0.053 | 0.214 |
| | *Prdm8*$^{-/-}$ +PTX | WT | 0.078 | 0.057 | 0.383 | -0.068 | 0.224 |
| | | *Prdm8*$^{-/-}$ | -0.081 | 0.055 | 0.334 | -0.222 | 0.061 |
| | | WT + PTX | -0.081 | 0.052 | 0.295 | -0.214 | 0.053 |
| Peak 8 | WT | *Prdm8*$^{-/-}$ | -0.207 | 0.064 | 0.011* | -0.369 | -0.046 |
| | | WT + PTX | -0.201 | 0.062 | 0.012* | -0.359 | -0.043 |
| | | *Prdm8*$^{-/-}$ + PTX | -0.039 | 0.059 | 0.790 | -0.188 | 0.111 |
| | *Prdm8*$^{-/-}$ | WT | 0.207 | 0.064 | 0.011* | 0.046 | 0.369 |
| | | WT + PTX | 0.007 | 0.058 | 0.993 | -0.139 | 0.152 |

| | | | | | | 95% Confidence Interval | |
|---|---|---|---|---|---|---|---|
| **Peak** | **Genotype** | **Genotype** | **Mean** | **SE** | **p value** | **Lower** | **Upper** |
| | | *Prdm8*$^{-/-}$ + PTX | 0.148 | 0.064 | 0.081 | -0.016 | 0.313 |
| | WT + PTX | WT | 0.201 | 0.062 | 0.012* | 0.043 | 0.359 |
| | | *Prdm8*$^{-/-}$ | -0.007 | 0.058 | 0.993 | -0.152 | 0.139 |
| | | *Prdm8*$^{-/-}$ +PTX | 0.116 | 0.053 | 0.103 | -0.020 | 0.253 |
| | *Prdm8*$^{-/-}$ +PTX | WT | 0.039 | 0.059 | 0.790 | -0.111 | 0.188 |
| | | *Prdm8*$^{-/-}$ | -0.148 | 0.064 | 0.081 | -0.313 | 0.016 |
| | | WT + PTX | -0.116 | 0.053 | 0.103 | -0.253 | 0.020 |
| Peak 9 | WT | *Prdm8*$^{-/-}$ | -0.223 | 0.069 | 0.012* | -0.398 | -0.047 |
| | | WT + PTX | -0.203 | 0.068 | 0.019* | -0.374 | -0.031 |
| | | *Prdm8*$^{-/-}$ + PTX | -0.046 | 0.056 | 0.697 | -0.189 | 0.097 |
| | *Prdm8*$^{-/-}$ | WT | 0.223 | 0.069 | 0.012* | 0.047 | 0.398 |
| | | WT + PTX | 0.020 | 0.062 | 0.946 | -0.138 | 0.178 |
| | | *Prdm8*$^{-/-}$ + PTX | 0.145 | 0.067 | 0.109 | -0.028 | 0.318 |
| | WT + PTX | WT | 0.203 | 0.068 | 0.019* | 0.031 | 0.374 |
| | | *Prdm8*$^{-/-}$ | -0.020 | 0.062 | 0.946 | -0.178 | 0.138 |
| | | *Prdm8*$^{-/-}$ +PTX | 0.103 | 0.051 | 0.138 | -0.028 | 0.234 |
| | *Prdm8*$^{-/-}$ +PTX | WT | 0.046 | 0.056 | 0.697 | -0.097 | 0.189 |
| | | *Prdm8*$^{-/-}$ | -0.145 | 0.067 | 0.109 | -0.318 | 0.028 |
| | | WT + PTX | -0.103 | 0.051 | 0.138 | -0.234 | 0.028 |
| Peak 10 | WT | *Prdm8*$^{-/-}$ | -0.234 | 0.065 | 0.005** | -0.398 | -0.069 |
| | | WT + PTX | -0.198 | 0.064 | 0.014* | -0.359 | -0.038 |
| | | *Prdm8*$^{-/-}$ + PTX | -0.013 | 0.075 | 0.983 | -0.206 | 0.179 |
| | *Prdm8*$^{-/-}$ | WT | 0.234 | 0.065 | 0.005** | 0.069 | 0.398 |
| | | WT + PTX | 0.035 | 0.059 | 0.821 | -0.113 | 0.184 |
| | | *Prdm8*$^{-/-}$ + PTX | 0.220 | 0.070 | 0.016* | 0.041 | 0.400 |
| | WT + PTX | WT | 0.198 | 0.064 | 0.014* | 0.038 | 0.359 |
| | | *Prdm8*$^{-/-}$ | -0.035 | 0.059 | 0.821 | -0.184 | 0.113 |
| | | *Prdm8*$^{-/-}$ +PTX | 0.131* | 0.048 | 0.033* | 0.010 | 0.253 |
| | | WT | 0.011 | 0.052 | 0.977 | -0.122 | 0.144 |

| | | | | | | 95% Confidence Interval | |
|---|---|---|---|---|---|---|---|
| **Peak** | **Genotype** | **Genotype** | **Mean** | **SE** | **p value** | **Lower** | **Upper** |
| | *Prdm8*$^{-/-}$ +PTX | *Prdm8*$^{-/-}$ | 0.220 | 0.070 | 0.016* | 0.041 | 0.400 |
| | | WT + PTX | -0.131 | 0.048 | 0.033* | -0.253 | -0.010 |

**Table S17**

**Statistical analysis of population spike amplitudes in the input-output responses of granule cells to increasing perforant path stimulation (from 50 to 400 µA in steps of 50 µA) *in vivo* in *Prdm8*$^{-/-}$ and wild type mice (n=9|9).** Mean responses were compared by ANOVA with repeated measures and were significantly different between genotypes. SS=sum of squares; MS=mean square; df=degrees of freedom.

| **Effect** | **SS** | **df** | **MS** | **F** | **p value** |
|---|---|---|---|---|---|
| Intercept | 4672.748 | 1 | 4672.748 | 92.138 | <0.001 |
| Genotype | 281.932 | 1 | 281.932 | 5.559 | 0.031 |
| Error | 811.438 | 16 | 50.715 | | |
| Intensity | 482.887 | 7 | 68.984 | 28.494 | <0.001 |
| Intensity*Genotype | 40.648 | 7 | 5.807 | 2.399 | 0.025 |
| Error | 271.154 | 112 | 2.421 | | |

**Table S18**

**Statistical analysis of the mean field excitatory post-synaptic potentials (fEPSP) slope of the input-output responses of granule cells to increasing perforant path stimulation (from 50 to 400 µA in steps of 50 µA) *in vivo* in *Prdm8*$^{-/-}$ and wild type mice (n=9|9).** Mean values were compared by ANOVA with repeated measures. SS=sum of squares; MS=mean square; df=degrees of freedom.

| **Effect** | **SS** | **df** | **MS** | **F** | **p value** |
|---|---|---|---|---|---|
| Intercept | 37844.19 | 1 | 37844.19 | 83.858 | <0.001 |
| Genotype | 1009.60 | 1 | 1009.60 | 2.237 | 0.154 |
| Error | 7220.66 | 16 | 451.29 | | |
| Intensity | 3836.00 | 7 | 548.00 | 29.061 | <0.001 |
| Intensity*Genotype | 571.12 | 7 | 81.59 | 4.327 | <0.001 |
| Error | 2111.97 | 112 | 18.86 | | |

**Table S19**

**Statistical analysis of population spike amplitudes in the baseline responses of granule cells *in vivo* evoked by test-pulse stimulations of the perforant path in *Prdm8*$^{-/-}$ and wild type mice (n=9|9).** Mean amplitudes were recorded at 23 time points after the first six baseline values and compared by ANOVA with repeated measures. SS=sum of squares; MS=mean square; df=degrees of freedom.

| Effect | SS | df | MS | F | p value |
|---|---|---|---|---|---|
| Intercept | 4364499 | 1 | 4364499 | 3784.681 | <0.001 |
| Genotype | 1990 | 1 | 1990 | 1.726 | 0.207 |
| Error | 18451 | 16 | 1153 | | |
| Time | 7503 | 22 | 341 | 1.439 | 0.093 |
| Time*Genotype | 6366 | 22 | 289 | 1.221 | 0.227 |
| Error | 83438 | 352 | 237 | | |

**Table S20**

**Statistical analysis of mean field excitatory post-synaptic potentials (fEPSP) slopes in the baseline responses of granule cells *in vivo* evoked by test-pulse stimulations of the perforant path in *Prdm8*$^{-/-}$ and wild type mice (n=9|9).** Mean slopes were recorded at 23 time points recorded after the first six baseline values and compared by ANOVA with repeated measures. SS=sum of squares; MS=mean square; df=degrees of freedom.

| **Effect** | **SS** | **df** | **MS** | **F** | **p value** |
|---|---|---|---|---|---|
| Intercept | 4506226 | 1 | 4506226 | 2688.314 | <0.001 |
| Genotype | 3422 | 1 | 3422 | 2.042 | 0.172 |
| Error | 26820 | 16 | 1676 | | |
| Time | 7011 | 22 | 319 | 1.160 | 0.282 |
| Time*Genotype | 7269 | 22 | 330 | 1.203 | 0.242 |
| Error | 96718 | 352 | 275 | | |

**Table S21**

**Statistical analysis of population spike amplitudes in the responses of granule cells *in vivo* evoked by one train of high frequency 400Hz stimulation of the perforant path of *Prdm8*$^{-/-}$ and wild type (WT) mice (n=9|9).** Mean amplitudes were recorded at 23 time points after the first six baseline values and compared by ANOVA with repeated measures. There was a significant interaction between time and genotype. SS=sum of squares; MS=mean square; df=degrees of freedom.

| **Effect** | **SS** | **df** | **MS** | **F** | **p value** |
|---|---|---|---|---|---|
| Intercept | 5888145 | 1 | 5888145 | 559.973 | <0.001 |
| Genotype | 6037 | 1 | 6037 | 0.574 | 0.460 |
| Error | 168241 | 16 | 10515 | | |
| Time | 12196 | 22 | 554 | 2.142 | 0.002 |
| Time*Genotype | 13389 | 22 | 609 | 2.351 | <0.001 |
| Error | 91120 | 352 | 259 | | |

**Table S22**

**Statistical analysis of mean field excitatory post-synaptic potentials (fEPSP) slopes in the responses of granule cells *in vivo* evoked by one train of high frequency 400Hz stimulation of the perforant path in *Prdm8*$^{-/-}$ and wild type (WT) mice (n=9|9).** Mean slopes were recorded at 23 time points recorded after the first six baseline values and compared by ANOVA with repeated measures. SS=sum of squares; MS=mean square; df=degrees of freedom.

| **Effect** | **SS** | **df** | **MS** | **F** | **p value** |
|---|---|---|---|---|---|
| Intercept | 5925362 | 1 | 5925362 | 931.848 | <0.001 |
| Genotype | 2084 | 1 | 2084 | 0.328 | 0.575 |
| Error | 101740 | 16 | 6359 | | |
| Time | 4661 | 22 | 212 | 0.746 | 0.791 |
| Time*Genotype | 8131 | 22 | 370 | 1.301 | 0.166 |
| Error | 99990 | 352 | 284 | | |

**Table S23**

**Statistical analysis of population spike amplitudes in the responses of granule cells *in vivo* evoked by 3 trains of high frequency 400Hz stimulation of the perforant path of *Prdm8*$^{-/-}$ and wild type (WT) mice (n=9|9).** Mean amplitudes were recorded at 23 time points after the first six baseline values and compared by ANOVA with repeated measures. Population spike amplitudes were significantly different between genotypes and there was a significant interaction between time and genotype. SS=sum of squares; MS=mean square; df=degrees of freedom.

| **Effect** | **SS** | **df** | **MS** | **F** | **p value** |
|---|---|---|---|---|---|
| Intercept | 7324256 | 1 | 7324256 | 310.743 | <0.001 |
| Genotype | 119435 | 1 | 119435 | 5.067 | 0.039 |
| Error | 377122 | 16 | 23570 | | |
| Time | 26588 | 22 | 1209 | 3.980 | <0.001 |
| Time*Genotype | 14191 | 22 | 645 | 2.124 | 0.003 |
| Error | 106893 | 352 | 304 | | |

**Table S24**

**Statistical analysis of mean field excitatory post-synaptic potentials (fEPSP) slopes in the responses of granule cells *in vivo* evoked by 3 trains of high frequency 400Hz stimulation of the perforant path in *Prdm8*$^{-/-}$ and wild type (WT) mice (n=9|9).** Mean slopes were recorded at 23 time points recorded after the first six baseline values and compared by ANOVA with repeated measures. SS=sum of squares; MS=mean square; df=degrees of freedom.

| Effect | SS | df | MS | F | p value |
|---|---|---|---|---|---|
| Intercept | 6498262 | 1 | 6498262 | 779.061 | <0.001 |
| Genotype | 12097 | 1 | 12097 | 1.450 | 0.246 |
| Error | 133458 | 16 | 8341 | | |
| Time | 8712 | 22 | 396 | 2.424 | <0.001 |
| Time*Genotype | 5002 | 22 | 227 | 1.392 | 0.114 |
| Error | 57519 | 352 | 163 | | |

**Table S25**

**Statistical comparison of mouse exploratory behavior of on the object pattern separation (OPS) task by each position of the object (1-5).** Mean discrimination index (D2) (±standard error of the mean) was compared across genotypes using Tukey's post hoc test for multiple comparisons. Genotypes of mice for statistical comparisons were: (i) *Bhlhb4*$^{-/-}$ (n=15); (ii) *Bhlhb4*$^{+/+}$*:* litter and sex matched WT controls for *Bhlhb4*$^{-/-}$ mice (n=10); (iii) *Prdm8*$^{-/-}$ (n=11); (iv) *Prdm8*$^{+/+}$: litter and sex matched WT controls for *Prdm8*$^{-/-}$ mice (n=8). SE=standard error; SS=sum of squares; MS=mean square; df=degrees of freedom; *=mean difference is significant at the 0.05 level.

| Object location | Genotype | Genotype | Mean Difference | SE | p value | 95% Confidence Interval | |
|---|---|---|---|---|---|---|---|
| | | | | | | Lower | Upper |
| Position 1 | *Bhlhb4*$^{+/+}$ (n=10) | *Bhlhb4*$^{-/-}$ | -0.018 | 0.084 | 0.996 | -0.245 | 0.209 |
| | | *Prdm8*$^{+/+}$ | 0.033 | 0.097 | 0.986 | -0.227 | 0.292 |
| | | *Prdm8*$^{-/-}$ | -0.042 | 0.091 | 0.966 | -0.287 | 0.202 |
| | *Bhlhb4*$^{-/-}$ (n=14) | *Bhlhb4*$^{+/+}$ | 0.018 | 0.084 | 0.996 | -0.209 | 0.245 |
| | | *Prdm8*$^{+/+}$ | 0.051 | 0.090 | 0.942 | -0.192 | 0.293 |
| | | *Prdm8*$^{-/-}$ | -0.024 | 0.084 | 0.992 | -0.251 | 0.202 |
| | *Prdm8*$^{+/+}$ (n=8) | *Bhlhb4*$^{+/+}$ | -0.033 | 0.097 | 0.986 | -0.292 | 0.227 |
| | | *Bhlhb4*$^{-/-}$ | -0.051 | 0.090 | 0.942 | -0.293 | 0.192 |
| | | *Prdm8*$^{-/-}$ | -0.075 | 0.097 | 0.864 | -0.335 | 0.185 |
| | *Prdm8*$^{-/-}$ *(n=10)* | *Bhlhb4*$^{+/+}$ | 0.042 | 0.091 | 0.966 | -0.202 | 0.287 |
| | | *Bhlhb4*$^{-/-}$ | 0.024 | 0.084 | 0.992 | -0.202 | 0.251 |
| | | *Prdm8*$^{+/+}$ | 0.075 | 0.097 | 0.864 | -0.185 | 0.335 |
| Position 2 | *Bhlhb4*$^{+/+}$ (n=10) | *Bhlhb4*$^{-/-}$ | 0.102 | 0.115 | 0.812 | -0.207 | 0.412 |
| | | *Prdm8*$^{+/+}$ | <0.001 | 0.128 | 1.000 | -0.343 | 0.344 |
| | | *Prdm8*$^{-/-}$ | -0.003 | 0.125 | 1.000 | -0.338 | 0.331 |
| | *Bhlhb4*$^{-/-}$ (n=12) | *Bhlhb4*$^{+/+}$ | -0.102 | 0.115 | 0.812 | -0.412 | 0.207 |
| | | *Prdm8*$^{+/+}$ | -0.102 | 0.119 | 0.826 | -0.422 | 0.217 |
| | | *Prdm8*$^{-/-}$ | -0.106 | 0.115 | 0.796 | -0.415 | 0.204 |

| Object location | Genotype | Genotype | Mean Difference | SE | p value | 95% Confidence Interval | |
|---|---|---|---|---|---|---|---|
| | | | | | | Lower | Upper |
| | *Prdm8*$^{+/+}$ (n=8) | *Bhlhb4*$^{+/+}$ | <-0.001 | 0.128 | 1.000 | -0.344 | 0.343 |
| | | *Bhlhb4*$^{-/-}$ | 0.102 | 0.119 | 0.826 | -0.217 | 0.422 |
| | | *Prdm8*$^{-/-}$ | -0.003 | 0.128 | 1.000 | -0.347 | 0.340 |
| | *Prdm8*$^{-/-}$ (n=10) | *Bhlhb4*$^{+/+}$ | 0.003 | 0.125 | 1.000 | -0.331 | 0.338 |
| | | *Bhlhb4*$^{-/-}$ | 0.106 | 0.115 | 0.796 | -0.204 | 0.415 |
| | | *Prdm8*$^{+/+}$ | 0.004 | 0.128 | 1.000 | -0.340 | 0.347 |
| Position 3 | *Bhlhb4*$^{+/+}$ (n=9) | *Bhlhb4*$^{-/-}$ | -0.107 | 0.103 | 0.728 | -0.384 | 0.170 |
| | | *Prdm8*$^{+/+}$ | -0.162 | 0.114 | 0.494 | -0.469 | 0.145 |
| | | *Prdm8*$^{-/-}$ | 0.153 | 0.110 | 0.513 | -0.144 | 0.451 |
| | *Bhlhb4*$^{-/-}$ (n=12) | *Bhlhb4*$^{+/+}$ | 0.107 | 0.103 | 0.728 | -0.170 | 0.384 |
| | | *Prdm8*$^{+/+}$ | -0.055 | 0.110 | 0.957 | -0.351 | 0.240 |
| | | *Prdm8*$^{-/-}$ | 0.260 | 0.106 | 0.085 | -0.025 | 0.545 |
| | *Prdm8*$^{+/+}$ (n=8) | *Bhlhb4*$^{+/+}$ | 0.162 | 0.114 | 0.494 | -0.145 | 0.469 |
| | | *Bhlhb4*$^{-/-}$ | 0.055 | 0.110 | 0.957 | -0.240 | 0.351 |
| | | *Prdm8*$^{-/-}$ | 0.315* | 0.117 | 0.049* | 0.001 | 0.630 |
| | *Prdm8*$^{-/-}$ (n=9) | *Bhlhb4*$^{+/+}$ | -0.153 | 0.110 | 0.513 | -0.451 | 0.144 |
| | | *Bhlhb4*$^{-/-}$ | -0.260 | 0.106 | 0.085 | -0.545 | 0.025 |
| | | *Prdm8*$^{+/+}$ | -0.315* | 0.117 | 0.049* | -0.630 | -0.001 |
| Position 4 | *Bhlhb4*$^{+/+}$ (n=10) | *Bhlhb4*$^{-/-}$ | 0.094 | 0.106 | 0.812 | -0.190 | 0.378 |
| | | *Prdm8*$^{+/+}$ | -0.016 | 0.119 | 0.999 | -0.336 | 0.304 |
| | | *Prdm8*$^{-/-}$ | 0.334* | 0.116 | 0.031* | 0.023 | 0.645 |
| | *Bhlhb4*$^{-/-}$ (n=15) | *Bhlhb4*$^{+/+}$ | -0.094 | 0.106 | 0.812 | -0.378 | 0.190 |
| | | *Prdm8*$^{+/+}$ | -0.110 | 0.109 | 0.747 | -0.403 | 0.183 |
| | | *Prdm8*$^{-/-}$ | 0.240 | 0.106 | 0.123 | -0.044 | 0.524 |
| | *Prdm8*$^{+/+}$ (n=8) | *Bhlhb4*$^{+/+}$ | 0.016 | 0.119 | 0.999 | -0.304 | 0.336 |
| | | *Bhlhb4*$^{-/-}$ | 0.110 | 0.109 | 0.747 | -0.183 | 0.403 |
| | | *Prdm8*$^{-/-}$ | 0.350 | 0.119 | 0.027* | 0.031 | 0.670 |
| | *Prdm8*$^{-/-}$ | *Bhlhb4*$^{+/+}$ | -0.334 | 0.116 | 0.031* | -0.645 | -0.023 |
| | | *Bhlhb4*$^{-/-}$ | -0.240 | 0.106 | 0.123 | -0.524 | 0.044 |

| Object location | Genotype | Genotype | Mean Difference | SE | p value | 95% Confidence Interval | |
|---|---|---|---|---|---|---|---|
| | | | | | | Lower | Upper |
| | (n=11) | *Prdm8*$^{+/+}$ | -0.350 | 0.119 | $0.027^{*}$ | -0.670 | -0.031 |
| Position 5 | *Bhlhb4*$^{+/+}$ (n=10) | *Bhlhb4*$^{-/-}$ | 0.049 | 0.093 | 0.951 | -0.120 | 0.298 |
| | | *Prdm8*$^{+/+}$ | 0.101 | 0.108 | 0.785 | -0.188 | 0.390 |
| | | *Prdm8*$^{-/-}$ | 0.133 | 0.099 | 0.547 | -0.134 | 0.399 |
| | *Bhlhb4*$^{-/-}$ (n=15) | *Bhlhb4*$^{+/+}$ | -0.049 | 0.093 | 0.951 | -0.298 | 0.120 |
| | | *Prdm8*$^{+/+}$ | 0.052 | 0.100 | 0.953 | -0.215 | 0.319 |
| | | *Prdm8*$^{-/-}$ | 0.083 | 0.090 | 0.793 | -0.159 | 0.325 |
| | *Prdm8*$^{+/+}$ (n=8) | *Bhlhb4*$^{+/+}$ | -0.101 | 0.108 | 0.785 | -0.390 | 0.188 |
| | | *Bhlhb4*$^{-/-}$ | -0.052 | 0.100 | 0.953 | -0.319 | 0.215 |
| | | *Prdm8*$^{-/-}$ | 0.031 | 0.106 | 0.991 | -0.252 | 0.315 |
| | *Prdm8*$^{-/-}$ (n=11) | *Bhlhb4*$^{+/+}$ | -0.133 | 0.099 | 0.547 | -0.399 | 0.134 |
| | | *Bhlhb4*$^{-/-}$ | -0.083 | 0.090 | 0.793 | -0.325 | 0.159 |
| | | *Prdm8*$^{+/+}$ | -0.031 | 0.106 | 0.991 | -0.315 | 0.252 |

**Table S26**

**Statistical comparisons of mouse exploratory behavior on the object pattern separation (OPS) task.** Mean discrimination index (D2) (±standard error of the mean) was compared across genotypes by two-way ANOVA. Genotypes of mice for statistical comparisons were: (i) *Bhlhb4*$^{-/-}$ (n=15); (ii) *Bhlhb4*$^{+/+}$: litter and sex matched WT controls for *Bhlhb4*$^{-/-}$ mice (n=10); (iii) *Prdm8*$^{-/-}$ (n=11); (iv) *Prdm8*$^{+/+}$: litter and sex matched WT controls for *Prdm8*$^{-/-}$ mice (n=8). SS=sum of squares; MS=mean square; df=degrees of freedom.

| **Effect** | **SS** | **df** | **MS** | **F** | **p value** |
|---|---|---|---|---|---|
| Genotype | 0.574 | 3 | 0.191 | 3.200 | 0.025 |
| Position | 7.584 | 4 | 1.896 | 31.729 | 0.000 |
| Genotype*Position | 0.969 | 12 | 0.081 | 1.352 | 0.193 |
| Error | 11.114 | 186 | 0.60 | | |

**Table S27**

**Statistical comparisons of mouse exploration behavior of left versus right objects on the object pattern separation (OPS) task.** Two-way ANOVA was used to compare the mean mouse exploration time in seconds of objects on the left vs objects on the right during the sample phase of the OPS task. Genotypes of mice for statistical comparisons were: (i) *Bhlhb4*$^{-/-}$ (n=15); (ii) *Bhlhb4*$^{+/+}$*:* litter and sex matched wild type (WT) controls for *Bhlhb4*$^{-/-}$ mice (n=10); (iii) *Prdm8*$^{-/-}$ (n=11); (iv) *Prdm8*$^{+/+}$: litter and sex matched WT controls for *Prdm8*$^{-/-}$ mice (n=8). SS=sum of squares; MS=mean square; df=degrees of freedom.

| **Effect** | **SS** | **df** | **MS** | **F** | **p value** |
|---|---|---|---|---|---|
| Genotype | 3671.456 | 3 | 1223.819 | 10.051 | <0.001 |
| Position | 40.219 | 1 | 40.219 | 0.330 | 0.566 |
| Genotype*Position | 24.361 | 3 | 8.120 | 0.067 | 0.978 |
| Error | 52358.970 | 430 | 121.765 | | |

**Table S28**

**Statistical analysis of learning behavior on the Morris Water Maze task.** The mean time spent searching for a hidden platform over 9 days of training was compared across genotypes using ANOVA with repeated measures. Genotypes for comparison were: (i) *Prdm8*$^{-/-}$ versus litter and sex-matched wild type mice (*Prdm8*$^{+/+}$) (n=11|10) and (ii) *Bhlhb4*$^{-/-}$ versus litter and sex-matched wild type mice (*Bhlhb4*$^{+/+}$) (n=12|11). SS=sum of squares; MS=mean square; df=degrees of freedom.

| | **Effect** | **SS** | **df** | **MS** | **F** | **p value** |
|---|---|---|---|---|---|---|
| *Prdm8*$^{+/+}$ vs *Prdm8*$^{-/-}$ | Genotype | 5580 | 1 | 5580 | 5.639 | 0.028 |
| | Day | 8541 | 8 | 1068 | 11.83 | <0.001 |
| | Genotype*Day | 834.2 | 8 | 104.3 | 1.155 | 0.330 |
| | Subjects | 18800 | 19 | 989.5 | 10.96 | |
| | Residual | 13721 | 152 | 90.27 | | |
| *Bhlhb4*$^{+/+}$ vs *Bhlhb4*$^{-/-}$ | Genotype | 2380 | 1 | 2380 | 5.544 | 0.028 |
| | Day | 24789 | 8 | 3099 | 30.46 | <0.001 |
| | Genotype*Day | 304.8 | 8 | 38.10 | 0.375 | 0.933 |
| | Subjects | 9015 | 21 | 429.3 | 4.220 | |
| | Residual | 17092 | 168 | 101.7 | | |

**Table S29**

**Statistical analysis of time spent in quadrant with hidden platform on day 10 of the Morris Water Maze task.** The mean time spent searching for a hidden platform on day 10 of training was compared between groups using the two-tailed t-test. Groups for comparison were: (i) *Prdm8*$^{-/-}$ versus litter and sex-matched WT mice (*Prdm8*$^{+/+}$) (n=10|10) and (ii) *Bhlhb4*$^{-/-}$ versus litter and sex-matched WT mice (*Bhlhb4*$^{+/+}$) (n=12|11). SEM=standard error of the mean; df=degrees of freedom.

| **Genotype** | **Mean time spent in quadrant with platform (+/- SEM)** | **t-value** | **df** | **p value** | **F-ratio** | **p Variances** |
|---|---|---|---|---|---|---|
| *Prdm8*$^{-/-}$ | 26.75 ± 3.587 | 2.24 | 18 | 0.038 | 2.003 | 0.315 |
| *Prdm8*$^{+/+}$ | 40.68 ± 5.077 | | | | | |
| *Bhlhb4*$^{-/-}$ | 37.62 ± 4.824 | 0.3326 | 21 | 0.743 | 3.415 | 0.063 |
| *Bhlhb4*$^{+/+}$ | 35.73 ± 2.727 | | | | | |

**Table S30**

**Statistical analysis of swimming speed on day 1 of the Morris Water Maze task.** The mean swimming speed on day 1 of training was compared between groups using the two-tailed t-test. Groups for comparison were: (i) *Prdm8*$^{-/-}$ versus litter- and sex-matched wild type mice (*Prdm8*$^{+/+}$) (n=11|10) and (ii) *Bhlhb4*$^{-/-}$ versus litter- and sex-matched wild type mice (*Bhlhb4*$^{+/+}$) (n=12|11). SEM=standard error of the mean; df=degrees of freedom.

| **Genotype** | **Mean speed on day 1 (+/- SEM)** | **t-value** | **df** | **p value** | **F-ratio** | **p Variances** |
|---|---|---|---|---|---|---|
| *Prdm8*$^{-/-}$ | 17.99 ± 0.860 | 0.318 | 19 | 0.754 | 1.506 | 0.550 |
| *Prdm8*$^{+/+}$ | 18.35 ± 0.735 | | | | | |
| *Bhlhb4*$^{-/-}$ | 17.22 ± 0.686 | 0.210 | 21 | 0.831 | 1.358 | 0.637 |
| *Bhlhb4*$^{+/+}$ | 17.02 ± 0.615 | | | | | |

**Table S31**

**Statistical comparisons of mouse exploration behavior on the Novel Object Recognition (NOR) task.** The mean discrimination index (D2) (±standard error of the mean) for all genotypes were compared to chance with the t-test. df=degrees of freedom.

| **Group** | **N** | **D2 ± SEM** | **t** | **df** | **p value** |
|---|---|---|---|---|---|
| *Prdm8*$^{+/+}$ | 8 | 0.363 ± 0.099 | -4.978 | 7 | 0.002 |
| *Prdm8*$^{-/-}$ | 11 | 0.421 ± 0.053 | -4.614 | 10 | 0.001 |
| *Bhlhb4*$^{+/+}$ | 10 | 0.352 ± 0.053 | -8.310 | 9 | <0.001 |
| *Bhlhb4*$^{-/-}$ | 12 | 0.294 ± 0.054 | -5.766 | 11 | <0.001 |

**Table S32**

**Statistical comparisons of mouse exploration behavior on the Novel Object Recognition (NOR) task.** The mean discrimination index (D2) (±standard error of the mean) for all genotypes were compared within groups and between groups by ANOVA. Genotypes in comparison groups were: (i) *Bhlhb4*$^{-/-}$ (n=12); (ii) *Bhlhb4*$^{+/+}$: litter- and sex-matched wild type controls for *Bhlhb4*$^{-/-}$ mice (n=10); (iii) *Prdm8*$^{-/-}$ (n=11); (iv) *Prdm8*$^{+/+}$: litter- and sex-matched wild type controls for *Prdm8*$^{-/-}$ mice (n=8). SS=sum of squares; MS=mean square; df=degrees of freedom.

| **Variable** | **SS** | **df** | **MS** | **F** | **P** |
|---|---|---|---|---|---|
| Between Groups | 0.430 | 3 | 0.143 | 2.029 | 0.127 |
| Within Groups | 2.611 | 37 | 0.71 | | |

**Table S33**

**Statistical analysis of learning behavior of *Prdm8*$^{-/-}$ and wild type mice on the contextual fear conditioning task (n=9|12).** Mean freezing time was compared within groups over days of training by ANOVA with repeated measures. There was no significant interaction between days and genotypes. SS=sum of squares; MS=mean square; df=degrees of freedom.

| **Effect** | **SS** | **df** | **MS** | **F** | **p value** |
|---|---|---|---|---|---|
| Intercept | 26273.36 | 1 | 26273.36 | 41.926 | <0.001 |
| Genotype | 1847.59 | 1 | 1847.59 | 2.948 | 0.102 |
| Error | 11906.46 | 19 | 626.66 | | |
| Days | 7609.46 | 4 | 1902.37 | 10.784 | <0.001 |
| Days*genotype | 430.36 | 4 | 107.59 | 0.610 | 0.657 |
| Error | 13406.50 | 76 | 176.40 | | |

**Table S34**

**Between group statistical analysis of learning behavior of *Prdm8*$^{-/-}$ and WT mice on the contextual fear conditioning task by day of training (n=9|12).** Mean freezing time was compared between groups on each day of training by t-test. Although *Prdm8*$^{-/-}$ mice tended to freeze for less time during the contextual fear conditioning task, there was no significant differences between genotypes on each of training. SS=sum of squares; MS=mean square; df=degrees of freedom. SD=standard deviation; df=degrees of freedom.

| **Variable** | ***Prdm8*$^{-/-}$ mean** | ***Prdm8*$^{-/-}$ SD** | **WT mean** | **WT SD** | **t value** | **df** | **p value** | **F ratio variances** | **p variances** |
|---|---|---|---|---|---|---|---|---|---|
| Day 1 | 0.485 | 0.791 | 0.968 | 1.525 | -0.863 | 19 | 0.399 | 3.713 | 0.073 |
| Day 2 | 7.847 | 10.472 | 17.880 | 17.146 | -1.547 | 19 | 0.138 | 2.681 | 0.172 |
| Day 3 | 13.995 | 14.121 | 23.922 | 18.423 | -1.344 | 19 | 0.195 | 1.702 | 0.460 |
| Day 4 | 17.699 | 20.756 | 29.858 | 23.364 | -1.236 | 19 | 0.231 | 1.267 | 0.754 |
| Day 5 | 18.694 | 19.403 | 28.475 | 17.528 | -1.209 | 19 | 0.241 | 1.225 | 0.736 |

**Table S35**

**Statistical comparisons of the results of observational behavioral measures in the modified SHIRPA protocol comparing adult wild type (WT) mice with adult *Prdm8*$^{-/-}$ mice (n=10|11).** Continuous measures (mean ± standard error of the mean) were compared between genotypes using the unpaired t-test. Categorical data were compared by $X^2$ test. P values in the table are uncorrected. There were no significant differences between genotypes after correction for multiple comparisons.

| | | **WT (n=10)** | ***Prdm8*$^{-/-}$ (n=11)** | **Uncorrected p values** |
|---|---|---|---|---|
| **Behavior in viewing jar** | | | | |
| Body position | Inactive | 0 | 0 | 1.0 |
| | Active | 10 | 11 | |
| | Excessive Activity | 0 | 0 | |
| Tremor | Absent | 10 | 11 | 1.0 |
| | Present | 0 | 0 | |
| Palpebral Closure | Eyes open | 10 | 11 | 1.0 |
| | Eyes closed | 0 | 0 | |
| Coat Appearance | Tidy | 9 | 10 | 1.0 |
| | Irregularities | 1 | 1 | |
| Whiskers | Present | 10 | 11 | 1.0 |
| | Absent | 0 | 0 | |
| Lacrimation | Absent | 10 | 11 | 1.0 |
| | Present | 0 | 0 | |
| Defecation | Present | 2 | 1 | 0.59 |
| | Absent | 8 | 10 | |
| **Behavior in arena** | | | | |
| Transfer Arousal | Extended Freeze | 0 | 0 | 1.0 |
| | Brief Freeze | 5 | 5 | |
| | Immediate Movement | 5 | 6 | |

| | | WT (n=10) | *Prdm8*$^{-/-}$ (n=11) | Uncorrected p values |
|---|---|---|---|---|
| Locomotor Activity | Squares entered | 9.60 ± 0.72 | 10.64 ± 0.51 | 0.25 |
| Gait | Fluid | 10 | 11 | 1.0 |
| | Lack of fluidity | 0 | 0 | |
| Tail Elevation | Dragging | 0 | 1 | 1.0 |
| | Horizontal Extension | 9 | 10 | |
| | Elevated | 1 | 0 | |
| Startle Response* | None | 2 | 0 | 0.035 |
| | Prayer Reflex | 6 | 11 | |
| | Reaction in addition | 2 | 0 | |
| Touch Escape | No response | 1 | 0 | 0.59 |
| | Response to touch | 8 | 8 | |
| | Flees prior to touch | 1 | 3 | |
| Defecation | Number | 0.2 ± 0.13 | 0 | 1.0 |
| **Behavior above arena** | | | | |
| Positional Passivity | Tail | 10 | 11 | 1.0 |
| Skin Color | Blanched | 0 | 0 | 1.0 |
| | Pink | 10 | 11 | |
| | Red | 0 | 0 | |
| Trunk Curl | Absent | 10 | 11 | 1.0 |
| | Present | 0 | 0 | |
| Limb Grasping | Absent | 8 | 5 | 0.39 |
| | Present | 3 | 6 | |
| Pinna Reflex | Present | 1 | 1 | 1.0 |
| | Absent | 9 | 10 | |
| Corneal Reflex | Present | 10 | 11 | 1.0 |
| | Absent | 0 | 0 | |
| Contact Righting Reflex | Present | 10 | 11 | 1.0 |
| | Absent | 0 | 0 | |
| Evidence of Biting | None | 10 | 11 | 1.0 |
| | Bites | 0 | 0 | |

| | | WT (n=10) | *Prdm8*$^{-/-}$ (n=11) | Uncorrected p values |
|---|---|---|---|---|
| Vocalization | None | 4 | 3 | 0.66 |
| | Vocal | 6 | 8 | |

**Table S36**

**Statistical comparisons of the results of observational behavioral measures in the modified SHIRPA protocol comparing adult wild type (WT) mice with adult *Bhlhb4*$^{-/-}$ mice (n=11|12).** Continuous measures (mean ± standard error of the mean) were compared between genotypes using the unpaired t-test. Categorical data were compared by $X^2$ test. P values in the table are uncorrected. There were no significant differences between genotypes after correction for multiple comparisons.

| | | **WT (n=11)** | ***Bhlhb4*$^{-/-}$ (n=12)** | **Uncorrected p values** |
|---|---|---|---|---|
| **Behavior in viewing jar** | | | | |
| Body position | Inactive | 0 | 0 | 1.0 |
| | Active | 11 | 12 | |
| | Excessive Activity | 0 | 0 | |
| Tremor | Absent | 9 | 12 | 0.22 |
| | Present | 2 | 0 | |
| Palpebral Closure | Eyes open | 11 | 12 | 1.0 |
| | Eyes closed | 0 | 0 | |
| Coat Appearance | Tidy | 11 | 10 | 0.48 |
| | Irregularities | 0 | 2 | |
| Whiskers | Present | 8 | 7 | 0.67 |
| | Absent | 3 | 5 | |
| Lacrimation | Absent | 11 | 12 | 1.0 |
| | Present | 0 | 0 | |
| Defecation | Present | 0 | 0 | 1.0 |
| | Absent | 11 | 12 | |
| **Behavior in arena** | | | | |
| Transfer Arousal | Extended Freeze | 0 | 0 | 1.0 |
| | Brief Freeze | 5 | 6 | |
| | Immediate Movement | 6 | 6 | |

| | | WT (n=11) | *Bhlhb4*$^{-/-}$ (n=12) | Uncorrected p values |
|---|---|---|---|---|
| Locomotor Activity | Squares entered | 10.64 ± 0.54 | 10.5 ± 0.5 | 0.68 |
| Gait | Fluid | 11 | 12 | 1.0 |
| | Lack of fluidity | 0 | 0 | |
| Tail Elevation | Dragging | 0 | 0 | 0.22 |
| | Horizontal Extension | 11 | 9 | |
| | Elevated | 0 | 3 | |
| Startle Response | None | 0 | 3 | 0.31 |
| | Prayer Reflex | 6 | 5 | |
| | Reaction in addition | 5 | 4 | |
| Touch Escape | No response | 2 | 2 | 1.0 |
| | Response to touch | 8 | 9 | |
| | Flees prior to touch | 1 | 1 | |
| Defecation | Number | 0 | 0 | 1.0 |
| **Behavior above arena** | | | | |
| Positional Passivity | Tail | 11 | 12 | 1.0 |
| Skin Color | Blanched | 0 | 0 | 1.0 |
| | Pink | 11 | 12 | |
| | Red | 0 | 0 | |
| Trunk Curl | Absent | 11 | 12 | 1.0 |
| | Present | 0 | 0 | |
| Limb Grasping | Absent | 7 | 11 | 0.16 |
| | Present | 4 | 1 | |
| Pinna Reflex | Present | 4 | 4 | 1.0 |
| | Absent | 7 | 8 | |
| Corneal Reflex | Present | 11 | 12 | 1.0 |
| | Absent | 0 | 0 | |
| Contact Righting Reflex | Present | 11 | 12 | 1.0 |
| | Absent | 0 | 0 | |
| Evidence of Biting | None | 11 | 12 | 1.0 |
| | Bites | 0 | 0 | |

| | | WT (n=11) | *Bhlhb4*$^{-/-}$ (n=12) | Uncorrected p values |
|---|---|---|---|---|
| Vocalization | None | 6 | 7 | 1.0 |
| | Vocal | 5 | 5 | |

**Table S37**

**Model parameter values and biophysical interpretations in neural mass model of the dentate gyrus.** The values of $A$, $B$, $G$ were based on López-Cuevas *et al* (*21*) (i.e., their order of magnitude and the fact that $A$ is expected to be lower than both $B$ and $G$). $a$, $b$, $g$, $v_0$, $e_0$, and $r$ were based on Jansen and Rit (*23*) and Wendling *et al* (*24*). The relations between the connectivity strengths $C_1$-$C_6$ were based on Danielson *et al* (*22*). $C_7$, $x_1$, $x_2$, and $x_3$ were calibrated such that the inputs to the sigmoid function were physiologically reasonable, i.e., close to $v_0$ (we used $x_1 = 10$, $x_2 = 0.5$, and $x_3 = 1$).

| Parameter | Interpretation | Value |
|---|---|---|
| $\boldsymbol{A}$ | Mean excitatory synaptic gain | 5 mV |
| $\boldsymbol{B}$ | Mean slow inhibitory synaptic gain | 10 mV |
| $\boldsymbol{G}$ | Mean fast inhibitory synaptic gain | 10 mV |
| $\boldsymbol{a}$ | Inverse average time constant: excitatory feedback loop | 100 /s |
| $\boldsymbol{b}$ | Inverse average time constant: slow inhibitory feedback loop | 50 /s |
| $\boldsymbol{g}$ | Inverse average time constant: fast inhibitory feedback loop | 500 /s |
| $\boldsymbol{C_1}$ | Connectivity strength: MC to GC | $16x_1$ |
| $\boldsymbol{C_2}$ | Connectivity strength: BC to GC | $1x_1$ |
| $\boldsymbol{C_3}$ | Connectivity strength: HIPP to GC | $8x_1$ |
| $\boldsymbol{C_4}$ | Connectivity strength: GC to MC | $400x_2$ |
| $\boldsymbol{C_5}$ | Connectivity strength: GC to BC | $20x_3$ |
| $\boldsymbol{C_6}$ | Connectivity strength: MC to BC | $80x_3$ |
| $\boldsymbol{C_7}$ | Connectivity strength: GC to HIPP | 100 |
| $\boldsymbol{v_0}$ | Firing threshold potential | 6 mV |
| $\boldsymbol{e_0}$ | Half of maximum firing rate of neural masses | 2.5 /s |
| $\boldsymbol{r}$ | Slope of the sigmoid transformation | 0.56 /mV |
| $\boldsymbol{p}$ | Noise level from the PP to GC and HIPP | 0.2 /s |

**Table S38**

**Modified Racine scale of seizure severity**

| Seizure stage | Behavioral expression |
|---|---|
| 0 | No change in behavior |
| 1 | Immobile |
| 2 | Head bobbing |
| 3 | Single clonus, intermittent rearing (<1 sec) |
| 4 | Bilateral clonus, continuous rearing (>1 sec) |
| 5 | Continuous rearing and falling |
| 6 | Generalised tonic seizures |
| 7 | Status epilepticus |

**Table S39**

**Comparison of median seizure scores induced in wild type mice and *Prdm8*$^{-/-}$ mice every 5 minutes for 2 hours after intraperitoneal kainic acid injections (n=6|6)**. Two-tailed Wilcoxon matched-pairs signed rank test was used to compare the maximum seizure scores between groups, which was statistically significant.

| **Wilcoxon matched pairs signed rank test** | ***Prdm8*$^{-/-}$ vs wild type** |
|---|---|
| Sum of positive, negative ranks | 26.00, -145.0 |
| Sum of signed ranks (W) | -119.0 |
| Number of pairs | 24 |
| Number of ties (ignored) | 6 |
| P value | 0.007 |

**Table S40**

**Statistical analysis of duration and time of onset of stage 3+ seizures between wild type (WT) mice and *Prdm8*$^{-/-}$ mice for two hours after intraperitoneal kainic acid injections (n=6|6).** Mean values were compared with the two-tailed unpaired t-test but there were no statistical differences between groups.

| | **Duration of stage 3+ seizures (minutes)** | **Onset of seizures stage 3+ (minutes)** |
|---|---|---|
| WT | 95 | 25 |
| | 55 | 5 |
| | 95 | 10 |
| | 100 | 5 |
| | 85 | 25 |
| | 15 | 60 |
| *Prdm8*$^{-/-}$ | 70 | 35 |
| | 50 | 20 |
| | 75 | 5 |
| | 85 | 40 |
| | 85 | 10 |
| | 55 | 10 |
| p value (2-tailed unpaired t-test) | 0.785 | 0.876 |
